# Intensive transmission in wild, migratory birds drove rapid geographic dissemination and repeated spillovers of H5N1 into agriculture in North America

**DOI:** 10.1101/2024.12.16.628739

**Authors:** Lambodhar Damodaran, Anna Jaeger, Louise H. Moncla

## Abstract

Since late 2021, a panzootic of highly pathogenic H5N1 has devastated wild birds, agriculture, and mammals. Analysis of 1,818 Hemagglutinin sequences from wild birds, domestic birds and mammals revealed that the North American panzootic was driven by ∼9 introductions into the Atlantic and Pacific Flyways, followed by rapid dissemination via wild, migratory birds. Transmission was primarily driven by Anseriformes, while non-canonical species acted as dead-end hosts. Unlike the epizootic of 2015, outbreaks in domestic birds were driven by ∼46-113 independent introductions from wild birds that persisted for up to 6 months. Backyard birds were infected ∼9 days earlier on average than commercial poultry, suggesting potential as “early warning signals” for transmission upticks. We find that wild birds are an emerging source of H5N1 in North America, necessitating enhanced, continuous surveillance. Prevention of agricultural outbreaks may now require novel strategies that reduce transmission at the wild bird/agriculture interface.

## Introduction

Highly pathogenic avian influenza (HPAI) viruses pose persistent challenges for human and animal health. Since emerging in 1996, highly pathogenic H5N1 viruses of the A/goose/Guangdong lineage have spread globally via endemic transmission among domestic birds in Asia and Africa coupled with long-distance dispersal by wild migrating birds^1,2^. In 2005, introduction of poultry-derived H5N1 viruses into wild birds in China led to viral dispersal across Northern Africa and Asia, establishing new lineages of endemic circulation in poultry^3,4^. In 2014, wild, migratory birds carried highly pathogenic H5N8 viruses from Europe to North America, sparking an outbreak that resulted in the culling of over 50.5 million commercial birds in the United States ^5^. While this outbreak substantially impacted the agriculture industry, these viruses did not establish persistent transmission in wild birds. Following aggressive culling of domestic birds, the epizootic was extinguished and North America remained free of HPAI for years.

In December 2021, clade 2.3.4.4b HPAI H5N1 viruses were introduced and spread across the Americas, causing a panzootic of significant morbidity and mortality in wild and domestic animals. These viruses were likely first introduced into North America in late 2021 by migratory birds flying across the Arctic Circle from Europe^6,7^, after which reassortment with low-pathogenicity avian influenza (LPAI) viruses endemic to North America produced a reassortant with altered neurotropism in experimentally infected mammals^8^. In contrast to past North American epizootics, morbidity and mortality has been widespread across a broad range of wild avian species not usually impacted by HPAI^9^ such as raptors, owls, passerines ^1,10^, and Sandwich Terns^11–13^. Infections have also occurred in mammal species not typically associated with HPAI, such as foxes, skunks, raccoons, harbor seals, dolphins, bears^1,14^, domestic goats, and dairy cattle^15^. The broad range of affected wild avian species raises the possibility that new reservoir hosts (hosts that would sustain continuous transmission chains) could be established, necessitating an evaluation of which species should be actively surveilled. Putative transmission among marine mammals and domestic dairy cattle pose new challenges for animal health and biosecurity, and highlight the need to understand the ecological factors that lead to spillover in mammals and agriculture^15,16^.

Historically, H5N1 transmission has been linked to enzootic transmission in domestic poultry^17,18^, paired with occasional cross-continental movement by wild birds of the Anseriformes (waterfowl such as ducks and geese) and Charadriiformes (Shorebirds) orders^19–21^. Unlike the North American epizootic in 2014-2015, widespread culling of domestic birds has not halted detections in North America, suggesting that patterns of transmission since 2022 may be distinct from past North American epizootics. Prior work has posited that clade 2.3.4.4b viruses may be better able to infect and transmit among wild bird species, leading to persistent, seasonal circulation in European wild birds and widespread reassortment^2,11,13^. In Europe, clade 2.3.4.4b viruses caused repeated incursions into European wild and domestic birds from 2016-2020, with a notable shift towards seasonal outbreaks and a broader range in infected wild species since 2020^22^. Thus, rather than acting as transient hosts that facilitate long-range movement, wild birds are now thought to play a greater role in maintaining and disseminating these viruses across Europe and Asia. However, the role of wild vs. domestic birds in driving transmission in North America has not been robustly or comprehensively studied, limiting informed enactment of surveillance and outbreak intervention strategies.

Previous genomic analysis of the United States epizootic linked outbreaks in poultry to wild birds, though the robustness of these results to differences in sampling between wild and domestic birds was not directly examined^10^. Other studies have reported surveillance and localized outbreak data to identify new incursions in wild bird populations^6,7,23^, but do not address the role of different species in contributing to cross-continental epizootic spread, transmission between species, or outbreaks in agriculture. As of 2025, highly pathogenic avian influenza viruses are classified as foreign animal diseases by the United States and Canada, with outbreak control plans primarily focused on biosecurity to reduce spread between farms, and rapidly culling domestic birds^24,25^. If the epizootic in North America reflects the changing ecology of clade 2.3.4.4b viruses towards wild bird driven transmission observed elsewhere, then surveillance, policy, and outbreak mitigation strategies may need to be fundamentally reformulated. Surveillance activities should focus on the host groups most important for viral dispersal, while control plans should be formulated to prevent incursions and spread in agriculture. Disentangling which species drove the North American epizootic, and whether transmission was driven by wild birds vs. domestic poultry, are therefore key for formulating effective surveillance and intervention strategies, but currently understudied. Finally, while it is currently thought that cases in mammals likely stem from infections in wild birds, work to formally link infections across species has been sparse.

Viral phylodynamic approaches are emerging as critical tools for outbreak reconstruction^26,27^. Viral genomes contain molecular records of transmission histories, allowing them to be used to trace how outbreaks begin and spread. Here we use Bayesian phylogeographic approaches paired with rigorous controls for sampling bias^28,29^ to trace how highly pathogenic H5N1 viruses were introduced and disseminated during the first 18 months of the epizootic across North America. We capitalize on a dataset of 1,824 Hemagglutinin gene sequences sampled from North American birds and mammals in 2021-2023, and curate additional metadata on geography, migratory flyways, domestic/wild status, host taxonomic order, and migratory behavior to reconstruct transmission between these groups. Using this dataset of HA sequences, we show that the epizootic in North America was driven by ∼9 independent introductions that descend from outbreaks in Europe and Asia, though only a single introduction spread successfully across the continent. The initial wave of H5N1 transmission spread from East to West by wild, migratory birds between adjacent migratory flyways. Transmission from non-canonical avian species like songbirds and owls was limited and resulted in dead-end transmission chains, suggesting that these species are unlikely to establish as reservoirs. Instead, transmission was primarily sustained by Anseriformes, suggesting these species as useful targets for surveillance activities. In contrast to the outbreak in 2014/2015, outbreaks in agriculture were seeded by ∼46-113 independent introductions from wild birds, with some onward transmission. Backyard birds were infected slightly more frequently, and earlier on average, than commercial birds, suggesting that increased backyard bird surveillance could be investigated as early warning signs for increased incidence. Together, these data pinpoint wild birds as emerging sources for H5N1 virus transmission in North America, capable of rapid, and long-distance viral dispersal and repeated reseeding of outbreaks in agriculture. Our results implicate continuous surveillance in wild aquatic migratory birds (particularly Anseriformes) as critical for contextualizing outbreaks in mammals and agriculture, and suggest that viral evolution in North America may now be increasingly governed by wild bird movement, ecology, and reassortment. Investment in interventions beyond culling that reduce interactions between domestic and wild animals may now be crucial for limiting future outbreaks in agriculture.

## Results

### Viral sequence data capture seasonal variation of HPAI detections

Most sequence data from North America represent detections in the United States and Canada (United States: 1590, Canada: 224, Central America: 8), with a “detection” classified as a positive PCR test from a collected sample. In Canada, surveillance in wild and domestic populations are coordinated by a collaborative effort from the Canadian Food Inspection Agency, Environment Canada, the Public Health Agency of Canada, and the Canadian Wildlife Health Centre, which perform year-round surveillance in live and dead wild birds, and case investigation for suspected poultry outbreaks^30^. In the United States, the United State Department of Agriculture Animal and Plant Health Inspection Service (APHIS) manages HPAI surveillance and testing in wild birds via investigation of reported morbidity and mortality events, hunter-harvested game birds/waterfowl, sentinel species/live bird collection, and environmental sampling of water bodies and surfaces^31,32^. USDA APHIS also surveilles domestic birds using several reporting methods: mandatory testing through the National Poultry Improvement Plan, coordination with state agricultural agencies, routine testing in high-risk areas, and backyard flock surveillance^33^.

Throughout the epizootic and during the time period analyzed in this study (November 2021-September 2023), most HPAI detections in the US have been reported in wild birds sampled via testing of sick and dead (5,611), hunter harvested (3,340), and live wild birds (1,127) (Figure S1a). Data on domestic bird detections are reported with information on poultry type (e.g., duck, chicken) and by whether the farm is classified as a commercial operation or backyard flock. Backyard flocks are categorized by the USDA as operations with fewer than 1,000 birds^34,35^ and by the World Organization for Animal Health (WOAH) as any birds kept in captivity for reasons other than for commercial production^36^. Among domestic birds, detections (1,177 total) came predominantly from commercial chickens (9.3%), commercial turkeys (28.5%), commercial breeding operations (species unspecified) (15.3%), and birds designated WOAH Non-Poultry which refers to backyard birds (42.3%) (Figure S1b). Other domestic bird detections occurred in game bird raising operations (2.5%) and commercial ducks (2.0%). The North American epizootic has impacted a broad range of mammalian hosts, with detections (399) reported in red foxes (24.3%), mice (24.1%), skunks (12.2%), and domestic cats (13.2%). Other mammalian hosts (26.2%) represent a wide range of species including harbor seals, bobcats, fishers, and bears (Figure S1c).

The first detection of HPAI H5N1 in North America was reported in migratory gulls in Newfoundland and Labrador Canada in November 2021, followed by the first detection in the United States in a wild American wigeon in South Carolina on December 30th, 2021^6^. From January – May 2022, a wave of 2,510 total detections were reported across 43 states and 91 different species (Figure 1). Following a lull in the summer of 2022, cases rose again in August 2022, leading to a larger epizootic wave that lasted until March 2023, totaling 8,001 detections across all 48 contiguous U.S. states and Alaska. Case detections peaked in the fall and spring, coinciding roughly with seasonal migration timing for birds migrating between North and South America^37,38^. Seasonal case variation could arise due to seasonal bird migration, fluctuating virus prevalence in wild birds, or fledging times of susceptible chicks^39^, though continued monitoring is necessary to determine whether these patterns persist in future years.

**Figure 1.**
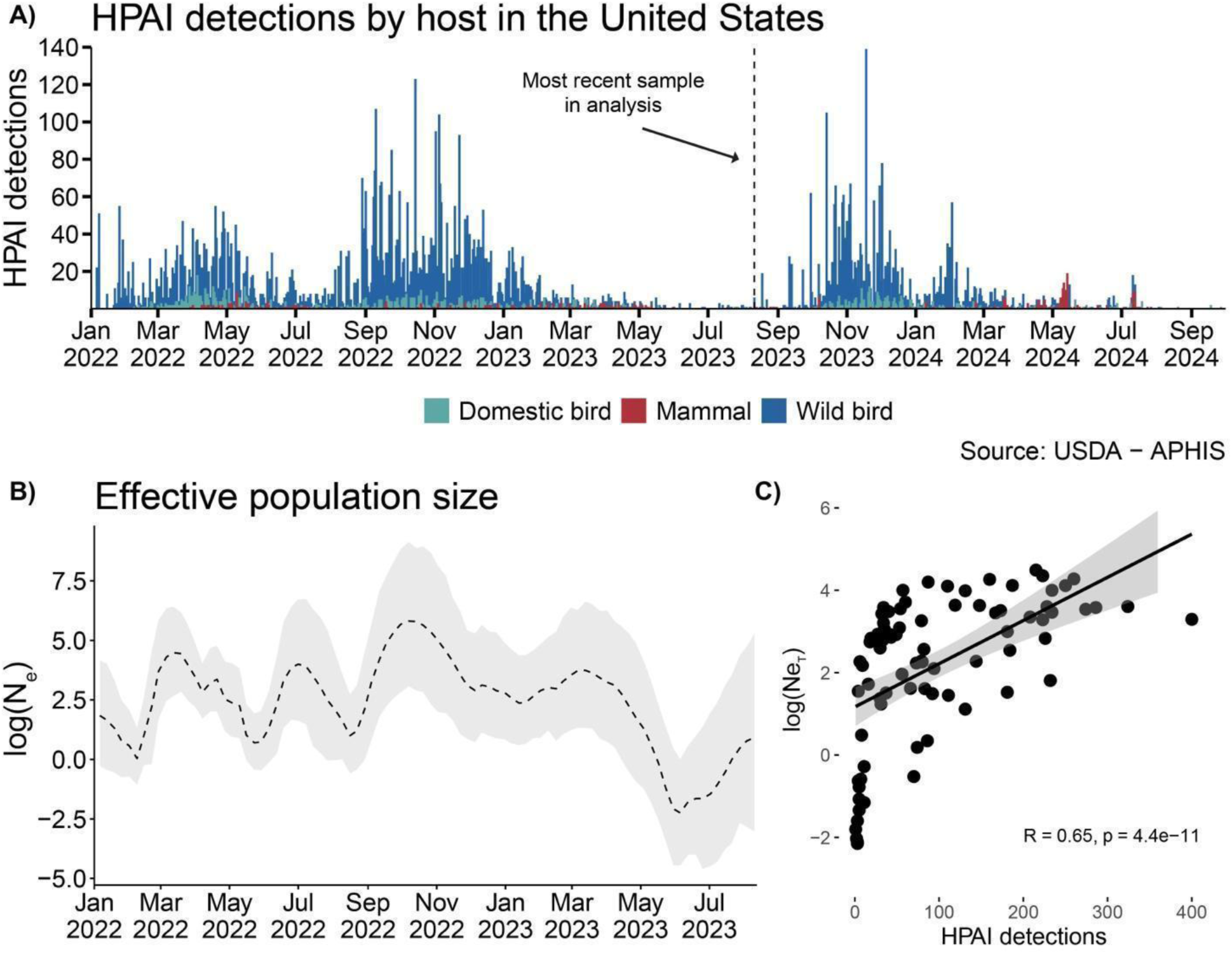
Detections of HPAI in North America show distinct epidemic waves following introduction events in late 2021. A) Detections of HPAI in wild birds, domestic birds, and non-human mammals. B) The Log-scaled Effective population size (N_e_) estimates estimated in BEAST using the Bayesian SkyGrid coalescent for sequences collected between Sep 2021 and Aug 2023. C) Correlation plot of log(N_e_) vs HPAI detections by week, spearman correlation displayed.

Sequence data sampled in North America is heavily skewed toward sequences from the United States (United States: 1590, Canada: 224, Central America: 8), and from the first 6 months of the outbreak, with 74% of all available sequences sampled from January-July 2022 (Figure S2). To evaluate whether sequence data reflect case detections, we inferred the viral effective population size (*Ne*), a measure of viral genetic diversity shown to be mathematically related to disease prevalence and the disease transmission rate^26^. We inferred *Ne* using a nonparametric population model (Skygrid), which captures relative changes in genetic diversity and the variability of growth rate in the virus population over time, providing a proxy for epidemic dynamics as previously described^26,40^. Using 6 datasets of sequences subsampled by host taxonomic order (see Methods for details), we infer that *Ne* is modestly correlated with detections (highest Spearman rank correlation: 0.65, p=4.4e-11) (Figure 1c)(Figure S3-4), and that peaks in *Ne* precede peaks in detections by ∼1 week (Figure S5), likely reflecting the lag between viral transmission and case detection. Thus, despite uneven sequence acquisition across time, the diversity of sampled sequences roughly reflect the amplitude of H5N1 cases. We therefore opted to use sequence data for the entire sampling period for broad inferences on introductions and geographic spread, but supplement these analyses with controls for sampling differences between groups. For more intensive reconstructions of transmission patterns between wild birds, commercial poultry, and backyard birds, we focus on the initial 6-month period with the most densely sampled data, coupled with experiments to assess the impacts of sampling on results. Finally, though we retained data from Canada and Central America for all subsequent analyses, our results are likely most informative about transmission within the United States due to the heavy skewing of data towards the United States.

### Highly pathogenic H5N1 was introduced multiple times from Europe and Asia

Though early case reports have described early introductions into the Atlantic and Pacific, a comprehensive and geographically representative analysis of the number and timings of introductions into North America across the continent has not been published ^6,8,23^. To reconstruct the number and timings of H5N1 introductions into North America during the first 18 months of the epizootic, we constructed a dataset that includes HA sequences from North America from domestic and wild birds (n= 1,327 unique isolates, identical sequences were removed), along with contextual sequences from other continents with ongoing outbreaks in 2021 and 2022 (Asia = 294, Europe = 300). We then inferred the number and timings of introductions into North America using a discrete trait phylogeographic model (see Methods for details)^41^.

Most sequenced infections in North America (98.5% of tips) descend from a single introduction from Europe in late 2021 (95% Highest Posterior Density (HPD), September 9th – October 7th 2021) (Figure 2a), consistent with reports of infected migratory gulls in Newfoundland and Labrador Canada in November 2021, and subsequent mortality in farmed birds in December of 2021^6,7,10^. We also recapitulate a second, short-lived introduction from Europe in 2022 as previously described by Erdelyan et al^8^. However, we also infer 7 (median = 7, 95% HPD: (6,8)) additional introductions between February and September 2022 that nest within the diversity of viruses circulating in Asia (Figure 2B-C), 5 of which have not been previously described. These introductions represent infections sampled in Alaska, Oregon, California, Wyoming and British Columbia, Canada that showed limited dissemination and persisted for short periods of time (0.024 – 6.9 months). The western location of these tips suggest potential introduction via the Pacific flyway, consistent with previous reports documenting incursions into North America from Japan via Alaska and the upper Pacific^42^ (Figure S6). Though none of these Pacific introductions had sampled descendants in the time period analyzed (up to September 2023), additional data available at time of writing indicate that one of these introductions re-emerged in late 2024 as the D1.1 lineage (annotated in Figure 2a)^43^. Though it remains unclear why this HA lineage was not detected from mid-2023 to 2024, the novel introductions documented here, and the eventual outgrowth of one of these lineages, highlights the importance of surveillance in the Pacific region for capturing viral importations. These data suggest that H5N1 viruses were introduced into North America at least 9 times, and that viral flow into the Pacific coast may be far more common than previously documented.

**Figure 2.**
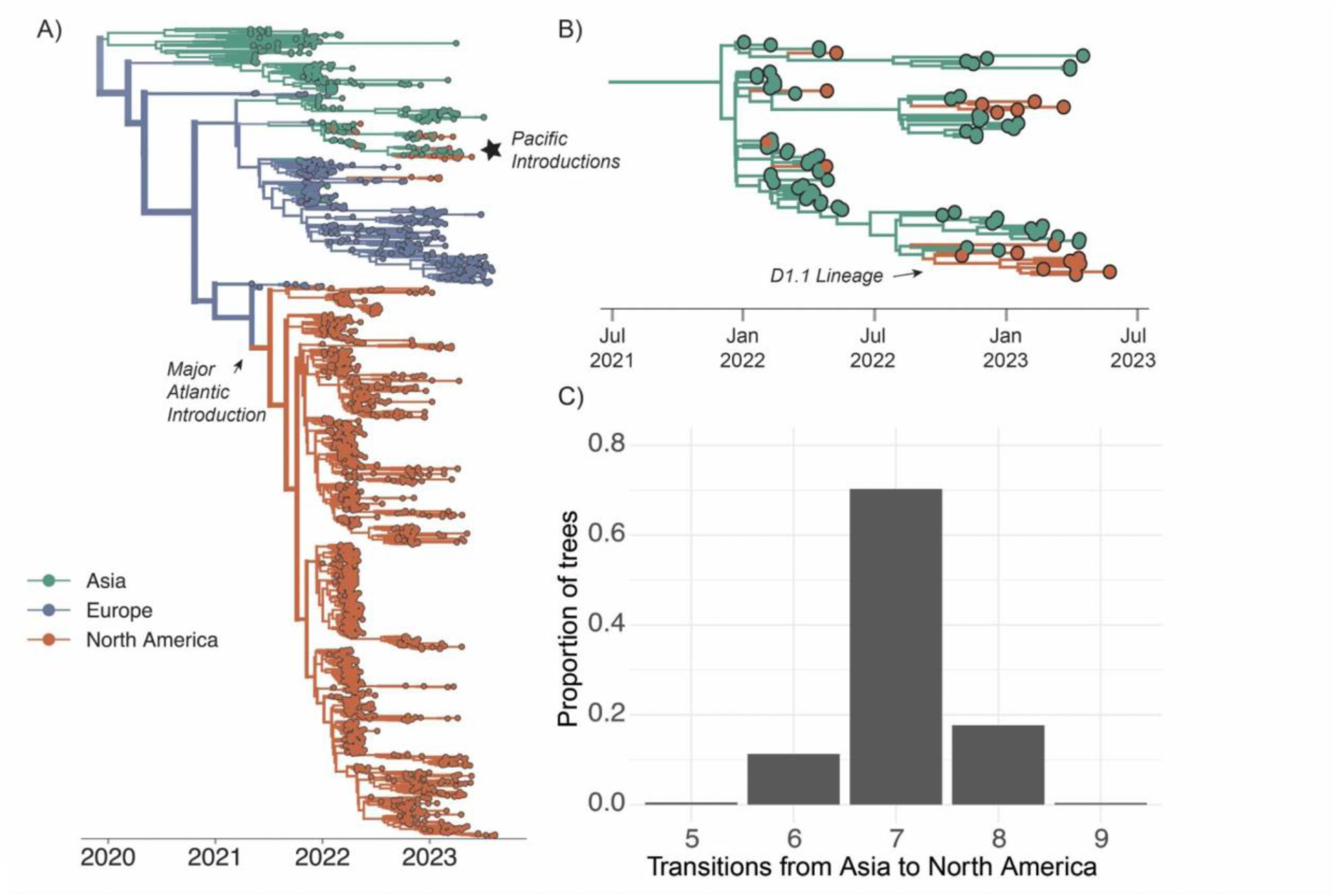
Introductions from Europe facilitated the majority of ongoing transmission while Asian introductions did not result in continent-wide onward transmission. A) Bayesian phylogenetic reconstruction of n=1,927 globally sampled sequences of HPAI clade 2.3.4.4b colored by continent of isolation. Opacity of branches corresponds to posterior support for the discrete trait inferred for a given branch, the thickness corresponds to the number of descendent tips the given branch produces. B) A close-up view of the starred section of the tree in A, focusing on introductions from Asia. C) We inferred the number of transitions from Asia to North America across the posterior set of 9,000 trees. The x-axis represents the number of introductions, and the y-axis represents the proportion of trees across the posterior set with that number of inferred transitions.

### Wild, migratory birds drove rapid expansion of H5N1 across the continent

Recent analyses of global H5N1 circulation patterns suggest wild birds as increasingly important sources for clade 2.3.4.4b virus evolution and transmission in Europe and Asia^17^. In the Americas, avian migratory routes are classified by the U.S. Fish and Wildlife Service (USFWS) into 4 major flyways: the Atlantic, Mississippi, Central, and Pacific^44^. If the epizootic were spread predominantly by wild, migratory birds, we reasoned that viruses sampled from the same, or neighboring flyways, should cluster together more closely than viruses sampled from non-adjacent flyways. To test this, we assigned avian sequences to the migratory flyway matching the US state of sampling, assembled a dataset of 250 sequences randomly subsampled for each USFWS flyway (total n=1,000), and implemented a discrete trait diffusion model to estimate transition rates between flyways as a proxy for viral movement between regions. To determine whether sequences clustered more strongly by flyway than expected by chance, we calculated the Association Index (AI), a measure of how strongly a trait is associated with a phylogenetic tree^45^. Finally, to determine whether movement between flyways was better supported than movement across other adjacent geographic areas, we quantified transitions between 4 North American regions stratified by latitude (see Methods for details).

Tips that descend from viruses circulating in Asia (those in Figure 2B) cluster together as a basal clade inferred in the Pacific flyway (orange cluster at top of tree, posterior probability = 0.98), consistent with introduction into the West Coast. The primary introduction from Europe occurred via the Atlantic Flyway, and subsequently spread rapidly across North America (Figure 3A,C). From the inferred time of introduction in the Atlantic coast (September 9th – October 7th 2021), viruses descending from this introduction had been sampled in every other flyway within ∼4.8 months, indicating markedly faster spread than observed for other avian-transmitted viruses^46^. Sequences clustered strongly by flyway (AI =10.563, p=0.00199), grouping most closely with those sampled within the same or geographically adjacent flyway (Figure 3A, Table S18).

**Figure 3.**
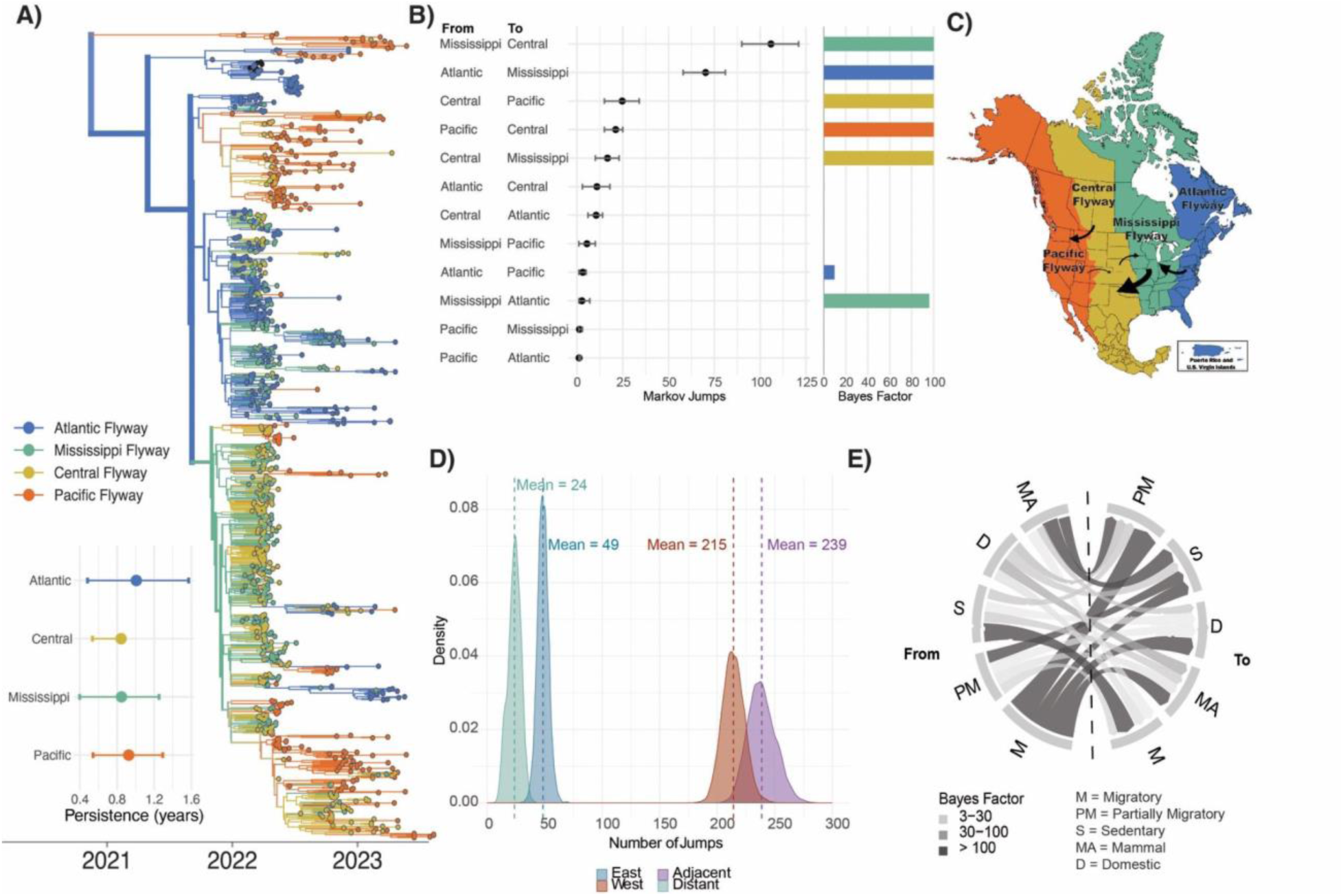
Migratory birds rapidly disseminated H5N1 via migratory flyways. A) Phylogenetic reconstruction of n=1,000 sequences colored by continental flyway. Inset is the results of the PACT analysis for persistence in each flyway (how long a tip takes to leave its sampled location going backwards on the tree), excluding the Pacific clade to show persistence following Atlantic introduction. B) Mean and 95% HPD for the number of markov jumps between USFWS flyways where the color of bar on the right of each jump pair corresponds to the source population and the height of the bar corresponds to the Bayes Factor (BF) support (where white corresponds to BF < 3 and full color corresponds to BF > 100). C) U.S. Fish and Wildlife Service waterfowl flyways map, with arrows annotated to represent rates with Bayes factor support of at least 100. Here the size of the arrow corresponds to the magnitude of the mean transition rate. D) Posterior distribution of the number of Markov jumps between flyways in the eastward or westward direction and between adjacent and distant flyways. E) Chord diagram of discrete trait diffusion based on migratory status going from the source population on the left to the sink population on the right where the thickness of the chord represents the mean transition rate and the color represents the Bayes Factor support.

Transitions (inferred as “Markov jumps”) between adjacent flyways were ∼10 times more frequent (mean = 239, 95% HPD: 216, 262) than those between distant flyways (mean = 24, 95% HPD: 12, 33)(Figure 3D, Figure S7), indicating a strong signal of dissemination via geographic proximity. Transitions between geographic groups stratified by latitude showed a similar trend towards transitions between adjacent (mean = 111, 95% HPD: 97, 125) vs distant (mean = 39, 95% HPD: 31, 46) regions, though the effect was less pronounced (Figure S9, Table S2), suggesting transmission that proceed most strongly among flyway regions. Transitions were predominantly inferred from East to West (Figure 3C, D, Table S1), with East to West jumps inferred ∼4.4 times more frequently (mean = 214, 95% HPD: (196, 232)) than West to East jumps (mean = 49, 95% HPD: (38, 57)) (Figure 3D), and 2.3-3.8 times more frequently than jumps along the North-South axis (Figure S9, Table S2).

We next calculated the Bayes Factor (BF) support for each between-flyway transition rate, and describe those very high statistical support (BF > 100, see Methods for details). We infer the highest and most well supported rates from the Mississippi to Central flyway (56.301 markov jumps/year, 95% HPD: 47.85, 64.33), Atlantic to Mississippi flyway (37.34 markov jumps/year, 95% HPD: 30.84, 43.065), and Central to Pacific flyway (13.127 markov jumps/year, 95% HPD: 7.975, 18.077)(Figure 3B, Figure S8, Table S1). Though the Pacific flyway experienced the highest number of introductions during the time period analyzed, transitions from the Pacific flyway elsewhere were inferred with low magnitude and weak support, consistent with the limited transmission we infer from West to East. Indeed, only a single statistically supported rate was inferred from the Pacific flyway to the adjacent Central flyway (BF = 3, 11.236 markov jumps/year, 95% HPD: 7.975, 13.292). Quantification of the length of time that each lineage persisted in each flyway showed slightly longer persistence within the Atlantic and Pacific flyways, though the estimates were variable (Figure 3A). We speculate that this pattern could reflect higher habitat and species richness within coastal flyways^47^, or that coastal flyways each only border 1 other flyway for viruses to transition to, potentially resulting in longer within-flyway persistence.

The strong clustering by flyways is consistent with long-range transmission by wild, migratory birds. To measure this, we next classified wild bird sequences by whether they were sampled from a wild bird considered migratory, partially migratory, or sedentary using the AVONET database^48^. We then modeled discrete trait diffusion across a subsample of 1000 sequences with equal proportions of the following 5 categories: wild migratory birds (includes most ducks and geese), wild partially migratory birds (some ducks, raptors, and vultures), wild sedentary birds (owls crows), domestic birds, and non-human mammals. Wild migratory and partially migratory birds are inferred at the root of the tree with probabilities substantially higher than expected based on their sampling frequencies (Figure S10 and Table S19), indicating a role for these species in sustained transmission across the epizootic. Transitions from wild migratory birds were inferred with the highest number and most strongly supported transition rates (BF > 3000), indicating that migrating wild birds were critical sources for infections in other species (Figure 3E, Table S3). In contrast, transitions from non-migratory wild birds were inferred with low magnitudes and weak support (Figure 3E, Table S3). Taken together, H5N1 viruses were then rapidly disseminated from East to West following a single introduction on the Atlantic coast, spreading between adjacent migratory flyways by wild, migrating wild birds. Viral dispersal was strongly associated with geographic proximity, with the strongest evidence for viral movement that proceeded westward. These results highlight the capacity of migratory birds to rapidly disseminate novel viral incursions across continental North America.

### Epizootic transmission was sustained by canonical host species

Previous outbreaks of highly pathogenic H5N1 viruses have been facilitated by wild Anseriformes (waterfowl) and Charadriiformes (shorebirds), and domestic species (Galliformes and Anseriformes)^18,49–52^, though the role of these hosts varies across outbreaks. While domestic ducks have played critical roles in bridging interactions between wild and domestic populations in past outbreaks in Asia, domestic ducks account for only 2% of all detections in the US, and the vast majority of cases in the North American panzootic have been in wild birds and Gallinaceous poultry (turkeys and chickens), potentially reflecting differences in poultry production between regions^53^. In the current panzootic, die-offs have occurred across a range of wild, non-canonical host orders, including Accipitriformes (raptors, condors, vultures), Strigiformes (owls), and Passeriformes (sparrows, crows, robins, etc.)^9^, raising the possibility that these new species could establish as reservoirs that merit surveillance. Pinpointing which species contributed to epizootic spread is thus critical for designing effective surveillance strategies. To determine whether particular host groups played outsized roles in driving transmission in the epizootic, we classified sequences by taxonomic orders that were most well-sampled and modeled transmission between them using a discrete trait model. We consolidated sequences of two orders of raptors, Accipitriformes and Falconiformes, hereafter referred to as “raptors”. We also consolidated two orders of pelagic birds, Charadriiformes and Pelecaniformes, hereafter referred to as “shorebirds”. Following classification, we defined 7 host order groups: Anseriformes, Shorebirds, Strigiformes, Passeriformes, Raptors, Galliformes, and non-human mammals.

Discrete trait approaches assume that the number of sequences in a dataset are representative of the underlying distribution of cases in an outbreak, resulting in faulty inference when this assumption is violated^29,41,54^ and bias when groups are unevenly sampled^28,29^. To account for differential sampling among these host groups, we therefore considered two, distinct subsampling approaches. The first is a proportional sampling regime in which sequences are sampled proportional to the detections in each host group each month. This common sampling regime assumes that case detections in each group are the closest proxy for the case distribution in the outbreak, and attempts to align sampling with underlying model assumptions. However, this approach may not be appropriate if case detection is heavily biased between groups. For HPAI H5N1 in North America, detections in wild birds are primarily identified when humans report sick or dead birds to wildlife health authorities or wildlife rescues (Figure S1A), which may skew detections towards birds with dedicated rescue services or birds that reside in closer proximity to humans. For example, Anseriformes and raptors comprised 50.2% and 20.3% of all sequences, respectively, which could arise from high case intensity or a higher rate of case acquisition. A second, complementary subsampling approach is to sample sequences equally, meaning that sequences are sampled from each group in perfectly equal numbers. By forcing the number of sequences from each group to be equal, the transmission inference must be driven by the underlying sequence diversity in each group rather than by sampling differences. Given the high variation among detections within each host group, we opted to pursue both sampling regimes and focus on results that were concordant in both. We first performed an Association Index test to confirm that clustering was sufficient for discrete trait inference (Table S18). Next, for each regime, we performed 3 independent subsamples, where the dataset was sampled either proportional to cases, or equally (see Methods for details). For the equal sampling regime, each dataset included 100 randomly sampled sequences per host group, except for Passeriformes for which only 57 sequences were available. To account for variation across subsampled datasets, we combined the results for the 3 independent subsamples to summarize statistical support (Figure S12, Table S4-5). Due to similar tree topologies across replicates, we visualized the phylogeny of the dataset with the highest posterior support (equal order subsample 1) below and make the results of all analyses available in supplement (see Figure S11, Table S6-S11). Finally, to measure the effects of potential sampling bias on the inferred transition rates, we performed a modified “tip shuffle” analysis. We generated 100 datasets in which the host tip assignments were randomly shuffled, re-inferred the host group at internal nodes, and infer a mean root state probability for each host across the 100 shuffled datasets. We then compared the root state probability in the empirical data to that inferred in the shuffled data as a measure of the impact of sampling on the results as previously described (see Methods for details)^55^.

The first introduction into North America is comprised of infections sampled in great black-backed gulls (large shorebirds cluster, posterior probability = 0.69), consistent with evidence of migratory gulls facilitating transmission from Europe (Figure 4A). This shorebirds cluster contains 6 sequences from harbor seals sampled from outbreaks in New England, resulting in a highly supported transition rate (BF = 537, posterior probability= 0.99) from shorebirds to non-human mammals (6.07 markov jumps/year, 95% HPD: (3.21, 8.91), aligning with suggestions that these outbreaks are linked to scavenging or environmental contamination by infected shore-birds^1,16^. After this initial cluster of infections, multiple deep, internal nodes on the phylogeny are inferred in Anseriformes with high posterior support (0.99), indicating that Anseriformes played an important role in driving sustained transmission and dispersal across North America.

**Figure 4.**
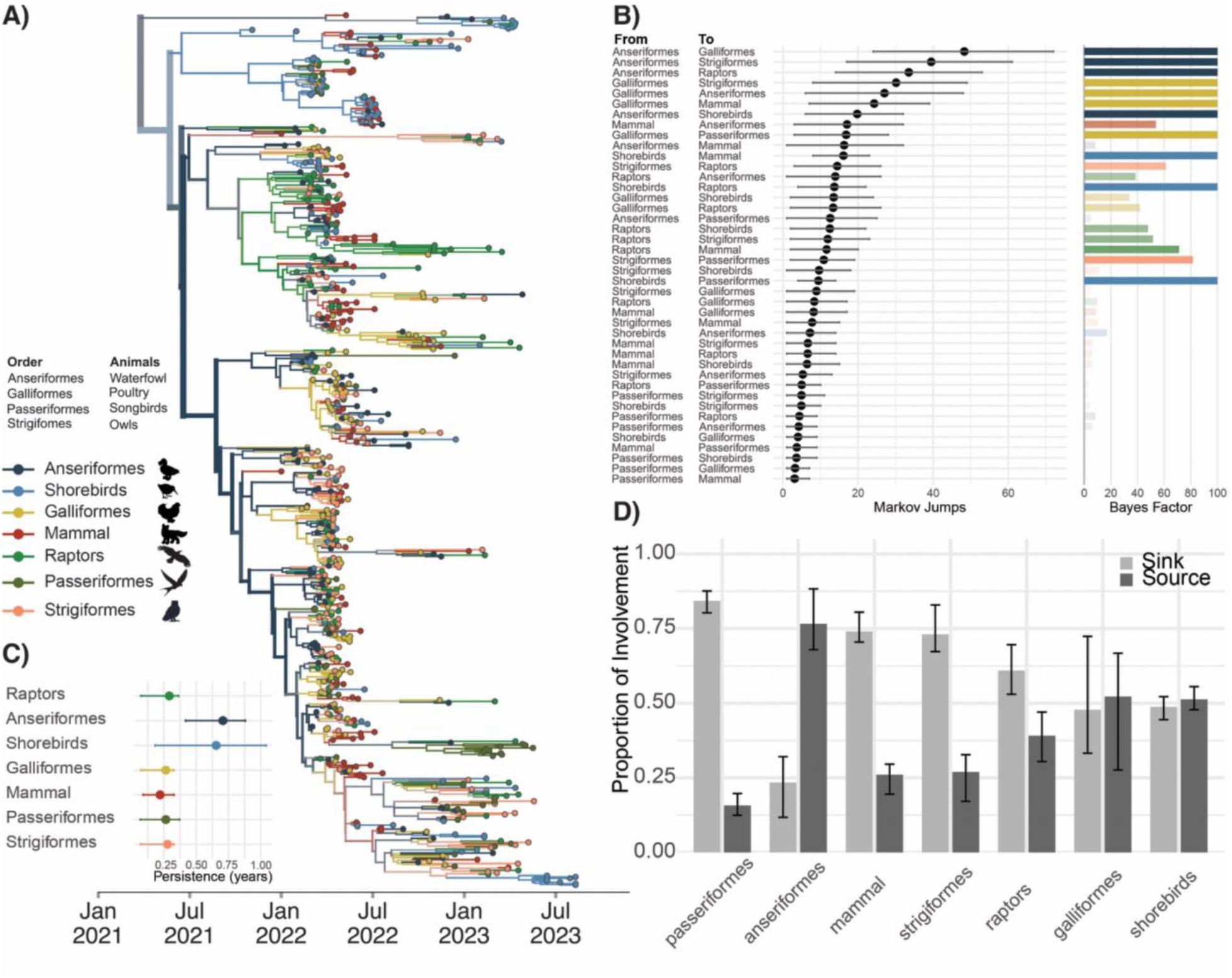
Anseriformes drove outbreak transmission, while new host species represent dead-end infections. A) Bayesian phylogenetic reconstruction of n=655 sequences subsampled by host order with equal proportions of each host. The color of tips and branches represents taxonomic order, and opacity represents the posterior support for the inferred host group. Thickness of branches correspond to the number of tips descending from a given branch. B) Mean number of Markov jumps and 95% HPD from the host group on the left (labelled “From”) to the host on the right (labelled “To”) as inferred from the combined results of three equal orders subsamples. The dot represents the mean number of Markov jumps and the black lines (whiskers) represent the 95% HPD. The corresponding bar plot shows the bayes factor support for each jump pair and color of each bar represents the “From” host. The opacity of the bar represents the Bayes factor support for the inclusion of the rate in the diffusion network. White (opacity of 0) represents any Bayes factors inferred to be less than 3, while a full color (opacity of 1) represents any Bayes factors inferred to be greater than or equal to 100. C) Results of the PACT analysis for persistence in each host order for phylogeny shown in panel A. D) For each host, we computed the proportion of Markov jumps involving that host order in which that host was inferred as a source (jump coming from that order) or as a sink (jump going to that order). The bars represent the variability across the 3 replicates of equal orders subsamples.

Across all replicates in both sampling regimes (6 total analyses), Anseriformes are inferred at the root 2-3 times more frequently than in null, shuffled datasets (Table S19), providing strong support for Anseriformes as critical drivers of epizootic transmission beyond what is expected from their frequency in the data. We infer Anseriformes as the predominant hosts seeding infections into other species (Figures 4B and 4D, Figure S13, Table S6-11), with the highest rates to Galliformes (17.81 markov jumps/year (95% HPD: (9.27, 26.02), BF = 1691, posterior probability = 0.99) and Strigiformes (13.51 markov jumps/year, 95% HPD: 5.35, 22.87, BF = 232, posterior probability = 0.99). Aligning with speculation following mortality events in bald eagles^56^, we also infer a highly supported transition rate (BF = 127, posterior probability = 0.95) from Anseriformes to Raptors, consistent with putative links between raptors and the waterfowl they predate. Each of these patterns were preserved in each independent subsample in both sampling regimes, indicating high robustness to sampling (Figure S11-12).

We also infer support for transmission from Galliformes to Anseriformes (7.86 markov jumps/year (95% HPD: (0.71, 14.97), BF = 147, posterior probability = 0.96), Strigiformes, and nonhuman mammals. In this dataset, Galliformes primarily represent domesticated poultry (98% of sequences), suggesting that transmission from domestic birds back to wild birds and mammals may also have occurred, a hypothesis we investigate in more depth below. However, lineages in Galliformes tended to be short-lived, persisting for 0.26 years on average (95% HPD: 0.07, 0.33 years). Additionally, Galliformes were inferred at the root less frequently than expected for their sampling frequency (Table S19), and tended to cluster together more strongly than expected by chance (p=0.0099, Table S18), consistent with a more limited role in transmission that may have been confined to localized agricultural outbreaks. In contrast, viral lineages persisted for the longest in Anseriformes (mean persistence time = 0.71 years, 95% HPD: 0.42, 0.88 years) (Figure 4C), and Shorebirds (mean persistence time = 0.654 years, 95% HPD: 0.18, 1.04 years). Shorebird sequences were very highly clustered with each other (AI=8.008, null=2.324, p=0.00999), suggesting some degree of separation between viruses circulating in Shorebird populations, consistent with ecological partitioning of low pathogenicity avian influenza viruses in these hosts ^57^. Tip shuffle analyses show mixed results for the Shorebirds, indicating a lack of consistent evidence for their role in transmission relative to their sampling frequency in the dataset. These data suggest that while Anseriformes, shorebirds, and Galliformes may all have contributed to transmission events to other species, that Anseriformes were the predominant drivers of longer-term persistence and spread to other hosts in the time period analyzed.

One surprising result was inference of raptors as a low-frequency, but statistically well supported source population to Anseriformes (5.18 markov jumps/year (95%HPD: (0.36, 9.27), BF = 39, posterior probability = 0.87). Previous characterizations of HPAI in Raptors during the 2014/2015 outbreak in North America showed mortality events and neurological symptoms in wild raptors^58^, while serological evidence of infections in bald eagles have indicated exposure to influenza A viruses in 5% of birds tested between 2006 and 2010^59^. In the ongoing panzootic, raptors represent the third most prevalent group in wild bird detections in Europe (12% of detections) and second most detected group in North America (20.3%)^13,60^. Tip shuffle results for both sampling regimes indicate that raptors are generally less probable at the root than expected based on their sampling frequency, indicating that the clustering of genetic sequences supports a limited role for epizootic transmission. Thus, while raptors may have been both heavily impacted and sampled at a high frequency, these data suggest that raptors were unlikely key drivers of epizootic spread. Future work to better establish the reasons for such high case numbers among raptors and to investigate their potential links to Anseriformes will be necessary for formulating wildlife management strategies.

We found limited support for non-canonical host groups (songbirds, owls, and nonhuman mammals) in seeding infections in other species. Passeriformes (songbirds), Strigiformes (owls), and mammals each primarily served as sinks for viral diversity (Figure 4B,D), with transitions inferred with low magnitude and weak support (Figure 4B). Summing the number of jumps originating from wild canonical (Anseriformes, shorebirds), wild noncanonical (Passeriformes, Strigiformes, raptors, mammals), and Galliforme (domestic) hosts confirm that noncanonical hosts primarily acted sinks that were far likelier to receive virus than propagate it onward (Figure S14), supporting short, terminal transmission chains that did not lead to long-term persistence (Figure 4C and Figure S15). Mammal sequences cluster across the entire diversity of the phylogeny (Figure 4A) and are not associated with one particular cluster of viruses, indicating that mammal infections were not confined to a particular viral lineage, supporting very short persistence times of 0.22 years (95%HPD:(0.088, 0.328)), and only one strongly supported transition rate to Anseriformes (BF = 53, posterior probability = 0.89). Instead, these findings are most compatible with a model in which wild mammals and other non-canonical species are infected by direct interaction with wild birds, likely related to scavenging and predation behavior^14^.

Taken together, these data suggest that despite high case numbers in several unusual wild hosts, non-canonical species generally played minor roles in transmission across the continent or to other species. Instead, epizootic transmission was most strongly supported in Anseriformes. We infer Anseriformes as predominant drivers of longer-term persistence and spread to other hosts across multiple independent analyses using distinct sampling regimes, supporting surveillance in these species for capturing trends in viral diversity and spread. Our data also suggest some role for shorebirds in supporting persistent viral transmission. Future work to disentangle the relationship between viral spread in these two key host groups may further resolve their utilities for surveillance and monitoring.

### Agricultural outbreaks were seeded by repeated introductions from wild birds

From 2022 to mid-2025, the US culled over 160 million domestic birds, with agricultural losses estimated between $2.5 to $3 billion USD ^61^. Understanding the degree to which outbreaks in agriculture have been driven by repeated introductions from wild birds vs. sustained transmission in agriculture is critical for improving surveillance and biosecurity practices. However, differences in sampling between wild and domestic birds challenge this goal. While domestic birds comprise 11% of all detections, they comprise 23.2% of sequences, making them overrepresented in available sequence data. In contrast, though a higher number of detections and sequences have been deposited for wild birds, wild birds are likely to be heavily under sampled due to the challenge of sampling wildlife^9,31^. Finally, while each detection in wild birds represents a single infected animal, domestic bird detections usually represent a single infected farm, where the true number of infected animals is unknown. Given these challenges, we designed a “titration” analysis to measure the impact of varying degrees of sampling on transmission inference between wild and domestic birds. We first generated a dataset composed of equal numbers of domestic and wild bird sequences (using all 270 available domestic bird sequences and 270 randomly sampled wild bird sequences) sampled between November 2021 and August 2023. By setting the number of sequences from each group equal, we force the inference to be driven by the sequence data itself, rather than the sampling regime. Next, we added in progressively more wild bird sequences until we reached a final ratio of domestic to wild sequences equal to 1:3, which approximates the ratio of detections in domestic and wild birds during the epizootic (1026 domestic detections vs. 3078 wild detections for study time period). In total, we generated 5 datasets with the following ratios of domestic to wild bird sequences: 1:1, 1:1.5,1:2,1:2.5, and 1:3 (see Methods for more details). For each dataset, we applied a discrete trait diffusion model to infer transmission between wild and domestic birds. The goal of this analysis was two-fold. First, we aimed to determine whether either domestic or wild birds would be inferred as the primary source population, and whether that inference would be robust to our choice in sampling regime. Second, by titrating in sequences at varying degrees, we hoped to assess whether the inferred number of transitions between hosts stabilized at a certain ratio. If inferred transitions increase linearly with sequences and never stabilize, this would indicate that more surveillance data is necessary. If not, this provides evidence that adding additional data does not result in altered results, suggesting that the currently available data may be sufficient for estimating dynamics within this time period.

When domestic/wild sequences were included in equal proportions, the root of the phylogeny and majority of internal nodes are inferred as wild birds, suggesting that wild birds are inferred as the primary source in the outbreak even when sampled evenly (Figure S16A). Wild birds are inferred at the root of the tree at a far higher probability than expected from their sampling (posterior probability = 0.895 in empirical data vs. 0.482 in tip shuffled data), while domestic birds are underrepresented (Table S19). Under equal sampling, this result is likely driven by higher genetic diversity among viruses sampled from wild birds, consistent with a large, source population. Within the background of wild bird sequences, domestic bird sequences form highly clustered groups (AI=23.096, p=0.0019, Table S18), consistent with some transmission between domestic birds. Transmission is inferred bi-directionally, with similar magnitudes of transmission inferred from domestic to wild birds, and from wild to domestic birds (Figure 5B, Figure S17, Table S12). If these patterns represent the true transmission history in the epizootic, we reasoned that these patterns should remain intact even when additional wild bird sequences are added to the tree. If not, then we hypothesized that some of domestic bird clusters may be disrupted as more wild bird sequences are added into the tree.

**Figure 5.**
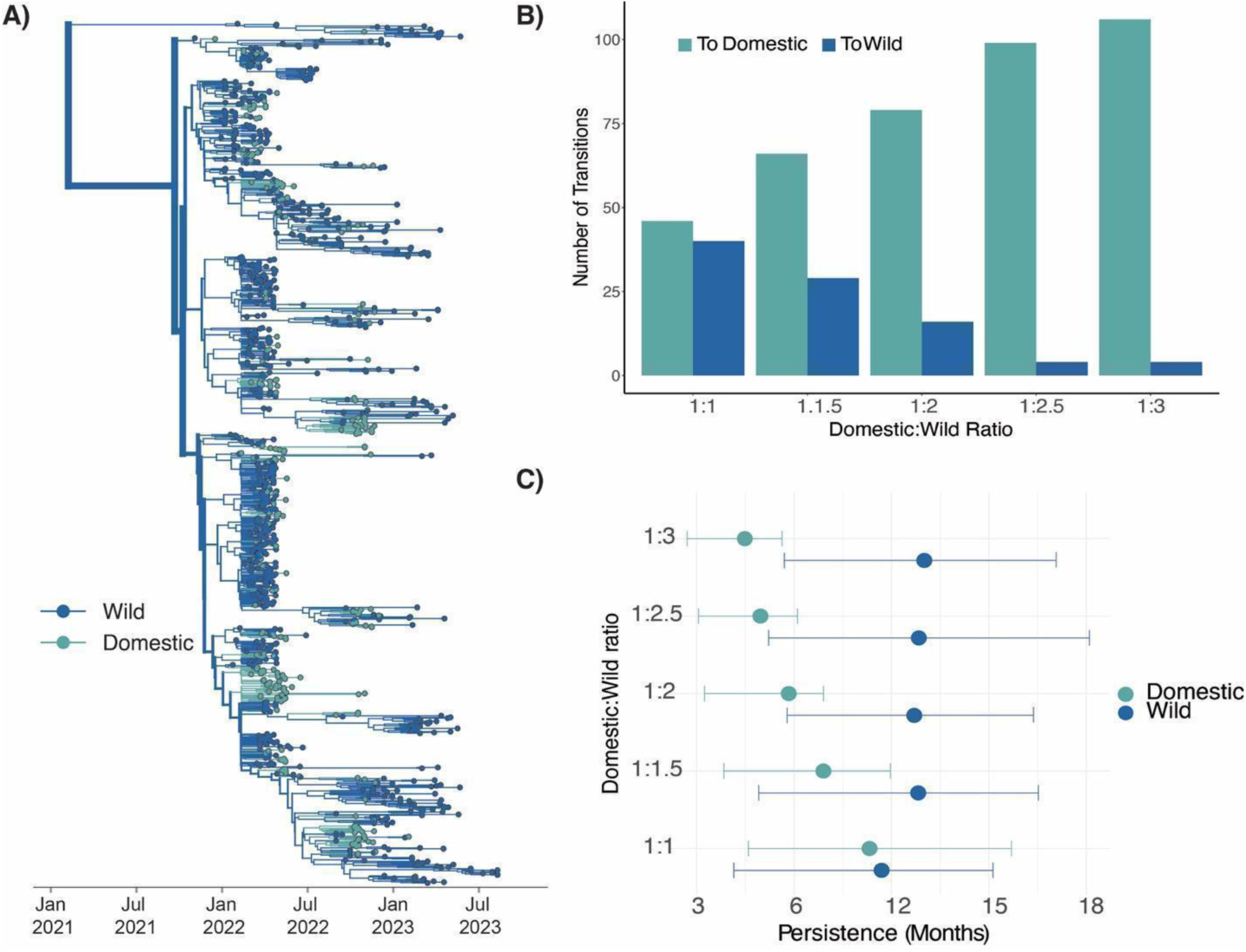
Outbreaks in domestic birds were seeded by repeated introductions from wild birds, with some onward transmission. A) Phylogenetic reconstruction where taxa and branches colored by wild or domestic host status containing a 1:3 ratio of domestic to wild bird sequences (n=1080). B) Number of transitions from a given trait to another trait inferred through ancestral state reconstruction for each titration. C) Results of the PACT analysis for persistence in domestic and wild birds for each titration.

As wild sequences were progressively added into the tree, most domestic-only clusters became smaller, broken up by wild sequences that interspersed within these clades (Figure S16A-E). The “breaking up” of these domestic clusters results in inference of more transitions from wild to domestic birds, and fewer transitions from domestic to wild birds (Figure 5B-C, Figure S17). The largest changes in the inferred transitions occurred between the 1:1 and 1:2.5 titrations, with minimal to no changes observed between transitions inferred in the 1:2.5 and 1:3 datasets, suggesting stability in the inferred transitions towards the end of the experiment (Table S12). The phylogeny of the final dataset (using a 1:3 ratio of domestic to wild sequences) shows 106 clusters of domestic sequences (Figure 5A, Figure S18-19, Table S12) each inferred as a unique introduction of H5N1 from wild birds into domestic birds. Among these domestic-only clusters, we calculate that lineages persisted for ∼4.5 months on average (95% HPD: 2.7, 5.63). In contrast, we infer limited transmission from domestic birds back to wild birds (Figure 5B, Table S12), with only 4 inferred introductions from domestic birds to wild birds (Table S12). Viral lineages in wild birds also persisted for over twice as long as those in domestic birds, persisting for ∼10 months (95% HPD: 5.7, 14.07) (Figure 5C).

Commercial turkey operations have been heavily impacted during the epizootic, comprising 53.7% of all detections on commercial farms^62^. However, the presence of wild turkeys throughout North America makes categorizing turkey sequences as domestic or wild status ambiguous. 98% of all turkey sequences are not associated with metadata on domestic/wild status, and thus were excluded from the previous analysis. However, epidemiologic data provides some hint that most deposited turkey sequences likely stem from domestic outbreaks. Among case detections during the study period, only 139 were reported in wild turkeys, representing 1.5% of all wild bird detections. In contrast, commercial turkey outbreaks comprised 28.5% of all domestic detections in the study period, suggesting that unlabelled turkey sequences are most likely to have come from domestic birds. While this data is not conclusive, we performed an additional analysis to determine whether our exclusion of turkey sequences (that are likely domestic) may have biased our results. In this analysis, any turkey sequence not labeled as “wild turkey” was categorized as “domestic” and combined with the other domestic bird sequences, resulting in a dataset in which ∼20% of domestic bird sequences were turkeys. We then created a second titration analysis using these domestic sequences to generate datasets of 1:1,1:1.5, and 1:2 ratios of domestic:wild sequences. We then evaluated the impact of the inclusion of turkey sequences on inferred transmission patterns between wild and domestic birds.

Inclusion of turkey sequences did not substantially change inferred transition rates between wild and domestic birds but did increase the inferred persistence times of domestic bird clusters (Figure S17, Table S12). As wild bird sequences were added to the tree, we observed the same “breakup” of domestic clades as in the above analysis, with more wild sequences leading to more inferred introductions into domestic birds, and very few into wild birds. In both titration experiments, the final number of inferred transmission events from domestic to wild birds was 4 (Table S12), indicating minimal transmission back to wild species, regardless of whether turkeys were included (Figure S20, Table S12). Inclusion of turkey sequences resulted in slightly longer inferred persistence times in domestic birds, increasing inferred persistence by 1.29 and 1.54 months in the 1:1.5 and 1:2 titrations (Figure S22). While turkey sequences tended to cluster closely with confirmed domestic sequences, they did form several large turkey-only clusters on the tree (Figure S21), indicating some degree of separation between turkey and non-turkey operations. To more directly estimate the role turkeys played in transmission we built a final 1:2 (domestic:wild) dataset where the number of turkey and domestic poultry were equal, which conformed to both a uniform and case proportional dataset. Surprisingly, we found that while most introductions into turkey populations stemmed from wild birds (42 transitions), we infer a high number of transmission events between turkeys and other domestic birds. We estimate ∼38 introductions from turkeys to other domestic birds, and 18 in the opposite direction (Figure S23, Table S13), indicating some degree of transmission between distinct poultry operation types. The high degree of transitions from wild birds to turkeys, and from turkeys to other domestic birds suggest a putative role for turkeys in mediating transmission between wild birds and other types of domestic poultry operations. In line with the high number of turkey detections during the epizootic, we infer comparatively larger clusters in turkeys than in other domestic birds, suggesting that turkeys may have played a key role in transmission among domestic populations.

Together, these data suggest a few important conclusions. First, wild birds are inferred strongly as the major drivers of transmission. Wild birds were inferred as the major source of viral dispersal even when heavily downsampled to be equal to domestic birds, and independent of whether or not turkeys were included in the analysis, indicating strong support for their role in H5N1 transmission. Second, regardless of sampling regime and presence/absence of turkey sequences, we infer that outbreaks in agricultural birds were driven by repeated, independent introductions from wild birds, with some onward transmission between domestic operations. While the exact number of inferred introductions vary across analyses (Table S12-13), we infer no fewer than 46, and as many as 113 independent introductions into domestic birds. When allowing sampling frequencies to approximate detections (the 1:3 ratio analysis), we resolve a higher number of introductions into domestic birds with shorter transmission chains, though lineages still persisted for an estimated 4-6 months. Analysis of turkey and non-turkey data indicate some degree of transmission between agricultural operations, and a potential role for turkeys in mediating transmission between wild birds and non-turkey domestic birds. Together, these results indicate that while both the 2014/2015 epizootic and the 2022 epizootic involved transmission between domestic premises, that transmission since 2022 has been fundamentally distinct. While the epizootic of 2014/2015 was started by a small number of introductions that rapidly propagated between commercial operations, the epizootic since 2022 has been driven by intensive and persistent transmission among wild birds, resulting in continuous incursions into domestic bird populations that have continuously sparked new outbreaks in agriculture. Wild birds thus played a critical role in agricultural outbreaks in North America from 2021-2023, marking a significant departure from past epizootics that may necessitate updates to biosecurity, surveillance, and outbreak control.

### Spillovers to backyard birds occur earlier and slightly more often than those to commercial birds

The 2014/2015 H5Nx epizootic in the United States was driven by extensive transmission in commercial poultry, prompting a series of biosecurity updates for commercial poultry farms^5,63^. However, not all domestic birds are raised in commercial settings. Rearing domesticated poultry in the home setting has become increasingly popular in the United States, with an estimated 12 million Americans owning “backyard birds” in 2022^64^. These birds have been heavily impacted during the ongoing epizootic, 579,005 backyard birds killed/euthanized, with some evidence for distinct transmission chains circulating in backyard birds vs. commercial poultry^10^. Because backyard birds generally experience less biosecurity than commercial birds and are more likely to be reared outdoors^34^, we hypothesized that spillovers into backyard birds may be more frequent than spillovers directly into commercial poultry.

To test this hypothesis, we used a subset of sequences sampled between January and May of 2022 that contained additional metadata specifying whether they were collected from commercial poultry or from backyard birds (n= 275 from commercial poultry, n=85 from backyard birds). We then built a tree that included an approximately equal number of sequences sampled from domestic and wild birds, but in which the domestic sequences were evenly split between commercial poultry and backyard birds (commercial birds = 85, backyard bird = 85, wild birds=193). As with the previous analysis, we infer wild birds as the primary source population and infer repeated introductions into commercial and backyard birds (Figure S24A). Unexpectedly though, backyard bird sequences appeared to cluster more basally than commercial poultry sequences, and sometimes fell directly ancestral to clusters of commercial poultry sequences (Figure S24A). While every introduction into backyard birds was inferred to descend from wild birds, 10 out of 26 introductions into commercial poultry were inferred to descend from backyard birds (Figure S25A). This pattern was reproducible across multiple replicate subsamples, indicating that it was independent of the exact subset of wild bird sequences included in the tree. Given the debated role of backyard birds in outbreaks in commercial poultry^65^, we further explored two hypotheses that could explain this pattern. The first is that backyard birds “mediated” transmission between wild birds and commercial birds, possibly due to their greater likelihood of outdoor rearing. Under this model, spillovers into backyard birds could be spread to commercial populations via shared personnel, clothing, or equipment, resulting in sequences from backyard birds preferentially nesting between wild and commercial bird sequences on the tree. Alternatively, backyard birds could have simply been infected earlier on average than commercial birds. If transmission in wild birds is persistent and high, and backyard birds have a higher risk of exposure due to lessened biosecurity and increased interactions with wildlife, then it could take less time for a successful spillover event to occur and be detected in these birds, resulting in clustering that is more basal in the tree.

To differentiate between these hypotheses, we performed a second titration analysis. We started with the phylogeny including equal numbers of sequences from commercial and backyard birds, allowing us to directly compare introduction patterns in these two groups. Then, we added in progressively more wild bird sequences into the tree until all available wild bird sequences were added into the tree. We added sequences in increments of 25% (where % refers to percentage of total available wild bird sequences in the time period), resulting in 3 additional analyses that included 50%, 75%, and 100% of all available wild bird sequences. For example, the final dataset included 942 sequences, comprising 85 commercial bird, 85 backyard bird, and 772 wild bird sequences. For each dataset, we inferred the number and timings of transmission events between wild birds, commercial birds, and backyard birds across the posterior set of trees. If backyard birds acted as mediators to outbreaks in commercial birds (hypothesis 1), then the relationship between backyard birds and commercial birds should remain unchanged as more wild bird sequences are added into the tree. Alternatively, if backyard birds and commercial birds were infected independently (hypothesis 2), then additional wild bird sequences should disrupt these clusters, and intersperse between commercial and backyard bird sequences, resulting in more independent introductions that occur earlier in backyard birds.

Throughout the experiment, wild bird sequences attached throughout the phylogeny, disrupting nearly every backyard bird-commercial bird cluster, and dissolving the signal of backyard bird to commercial bird transmission originally observed (Figure S24). The final tree that included all available wild bird sequences resulted in inference of ∼82 independent introductions from wild birds to domestic birds, with most clusters containing only commercial (39 clusters) or backyard bird (43 clusters) sequences (Figure 6A-B, Figure S24-25), suggesting that outbreaks in these groups were likely seeded independently. Indeed, of the initial 10 transmission events inferred from backyard birds to commercial birds, only 2 remained undisturbed in the final tree (Figure 6B, Figure S25). These two events represent outbreaks that occurred in the same state within 6 days of each other, so it is plausible that these outbreaks are directly linked. However, all other clusters were disrupted. As wild bird sequences were added into the tree, the number of inferred introductions into backyard birds and commercial birds diverged across the posterior trees for each titration (Figure S26), with backyard birds experiencing slightly more introductions (mean = 42 introductions, 95% HPD: (35, 49)) than commercial poultry (mean = 39 introductions, 95% HPD: (32, 44)) (Figure 6C). This is consistent with the rate of transmission inferred from wild birds into each domestic population with a slightly higher rate to backyard birds (48.963 markov jumps/year, 95% HPD: 40.163, 56.228, BF = 18,000, posterior probability = 0.99), then to commercial birds (46.48 markov jumps/year, 95% HPD: 40.163, 52.785, BF = 18,000, posterior probability = 0.99) (Table S14-S17). We infer a low transition rate from backyard birds to commercial birds (2.678 markov jumps/year, 95% HPD:1.148, 4.59, BF = 4.58, posterior probability = 0.7), and transitions from domestic birds back to wild birds were not statistically well-supported.

**Figure 6.**
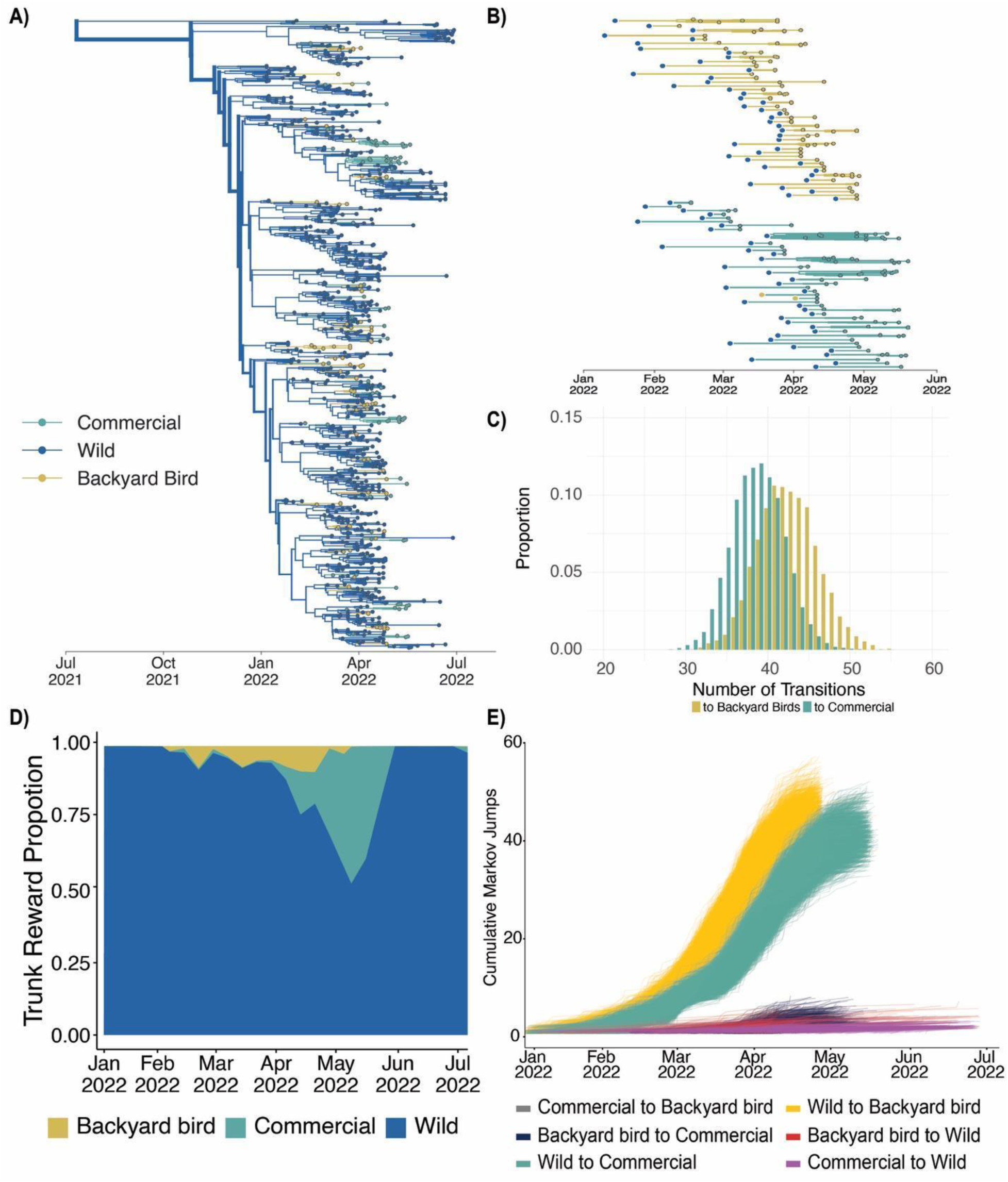
Backyard birds are infected by wild birds earlier than commercial birds. A) Phylogenetic reconstruction of sequences collected between Jan 2022 and May 2023 with all available wild bird sequences and equal proportions of commercial and backyard birds (n= 942) where taxa and branches are colored by host domesticity status. B) Exploded tree view of the phylogeny showing the branches of transmission in each domestic bird type following transmission from wild birds where subtrees represent the traversal of a tree from the root to the tip where the state is unchanged from the initial state (given by the large dot on left) to the tips represented by the smaller dots representing continuous chains of transmission within a given state. C) Proportion of trees from the posterior tree set with a given number of transitions from wild birds to backyard birds and commercial birds (100% available wild sequences). D) Markov rewards trunk proportion for domesticity status showing the waiting time for a given status across branches of the phylogeny over time. E) Cumulative Markov jumps from a given bird type to another over time where each line represents a single phylogeny from the posterior sample of trees.

To determine whether spillovers into backyard birds occurred earlier than those into commercial poultry, we estimated the number of transitions between hosts across the phylogeny (“Markov jumps”) and the amount of time that is spent in each host between transitions (“Markov rewards”) ^66,67^. We infer the highest mean duration in wild birds, representing 87.7% of Markov rewards. Backyard bird and commercial bird sequences showed lower, and similar reward time percentages of 5.3% and 7.0% respectively. The Markov Reward proportion is calculated as the proportion of the phylogeny at a given time being a given discrete state. By looking at the proportion a given state is over time across the phylogeny we can provide a proxy for how long transmission has occurred in each group between transition events. We find that early in the epizootic, transmission in backyard birds slightly preceded transmission in commercial poultry (Figure 6D, Figure S25). Enumeration of the cumulative number of transitions between hosts over time across the posterior set of trees (Markov jumps^66^), again revealed that backyard birds experienced slightly more jumps than commercial poultry (backyard birds = 43 introductions, 95% HPD: 36, 50; commercial birds = 39 introductions, 95% HPD: 32, 44), and that these introductions occurred earlier on average (Figure 6E). The lag time between the cumulative transitions for backyard birds and commercial birds was ∼9.6 days, indicating that transitions to backyard birds were inferred ∼9 days earlier than those to commercial birds. Comparison of detections and sequence availability in commercial birds vs backyard birds show no apparent skewing in availability of samples for each group in that time period, suggesting that this pattern is not simply due to an abundance of earlier cases in backyard birds at that time (Figure S27). While this pattern could also arise if cases in backyard birds are systematically reported earlier than commercial poultry, data on testing turnarounds and enrollment in the US indemnity payment register suggest that this is unlikely to be the case. Commercial and backyard bird farms have almost identical lag times between case reporting and confirmation (2.15 days for commercial birds and 2.4 days for backyard birds)^68^, and commercial poultry operations generally depopulate completely within 24 hours^69^, indicating testing and response in commercial operations that is efficient and slightly earlier than in backyard birds. Comparison of the proportion of farms that received indemnity payments (as a proxy for the percentage that submit to regular testing) shows that while 511 of 168,048 commercial operations (0.3%) reported cases, that only 656 out of ∼12 million backyard bird owners (0.0055%) were enrolled in the program^64,68,70^. While complete data on the distribution of backyard bird operations within North America are sparse, these data suggest that backyard birds are likely undersampled relative to commercial poultry, with somewhat lagged testing, making the early pattern we observe compelling. Future studies utilizing expanded datasets across future epizootic waves are necessary to confirm this pattern more broadly.

These data confirm that in the first 6 months of the epizootic, outbreaks in backyard bird and commercial bird populations were generally seeded independently, with limited evidence for transmission between them. We show phylogenetic evidence that spillovers into backyard birds may have occurred slightly more frequently and earlier on average than spillovers into commercial poultry, despite epidemiologic data that testing is slower and more limited in backyard birds. Should future studies support these findings more broadly, then backyard bird populations could potentially be investigated as early warning signals for upticks in transmission. Future work will be necessary to investigate the utility of this hypothesis.

## Discussion

The panzootic of HPAI H5N1 that began in 2021 has impacted wildlife health, agriculture, and human pandemic risk across the Americas. The North American epizootic has been distinguished by the high number of infections in wildlife not usually impacted by HPAI, its rapid dissemination across the Americas, and for its persistence despite aggressive culling of domestic birds. In this study, we used a dataset of 1,824 HA sequences paired with curated metadata to reconstruct how H5N1 viruses were introduced and spread. We show that H5N1 viruses were introduced ∼9 independent times into North America, with repeated incursions into the Pacific flyway that mostly persisted transiently. A single introduction into the Atlantic flyway spread across the continent within ∼4.8 months, transmitting from East to West by wild, migratory birds. Long-range dispersal and persistence were primarily driven by transmission in Anseriforme species, with infections in wild bird species like songbirds and owls primarily serving as dead-end hosts. Finally, we show through repeated subsampling experiments that unlike the epizootic of 2014/2015, the epizootic since 2021 was fundamentally distinct because transmission was driven by wild, migrating birds. Wild birds repeatedly re-seeded outbreaks in agriculture, sparking new introductions that persisted despite improvements in biosecurity and rapid culling responses. Outbreaks in commercial poultry and backyard birds were generally seeded independently, with introductions into backyard birds occurring ∼9 days earlier on average, though future work is necessary to confirm these patterns more broadly. Together, our results suggest a critical shift in the ecology of highly pathogenic avian influenza viruses in North America. As the primary source of transmission shifts from poultry to wild migratory birds, the ecology of clade 2.3.4.4b viruses in North America may now follow patterns unfolding globally, where evolution is increasingly governed by wild bird movement, ecology, and reassortment. Our results implicate surveillance in wild Anseriforme species as potentially fruitful targets for ongoing tracking and response, and highlight the necessity of continued surveillance in wild birds for accurate outbreak reconstruction. Finally, our results provide explanations for the rapid expansion across the continent and for why culling domestic birds may no longer be sufficient for preventing outbreaks in agriculture. Instead, layered interventions, like improved biosecurity, separation of wild and domestic birds, and domestic animal vaccination may now be necessary for reducing future spillovers to agriculture, and by extension, humans.

Our study develops multiple lines of evidence that collectively support wild birds as a critical emerging source of highly pathogenic avian influenza transmission in North America. By directly modeling transitions between host groups based on domestic/wild classification, taxonomic order, and migratory behavior, paired with geographic analyses showing strong dispersal across flyways, we show that wild birds were key drivers of the epizootic. These results imply that continuous surveillance in wild birds may now be key for viral tracking and outbreak reconstruction, with our data pointing to Anseriformes as good potential targets. Recent modeling of HPAI risk in Europe identified *Anatinae* and *Anserinae* (within the order Anseriformes) species prevalence as the most consistent predictors of HPAI detection across seasons^71^, in line with our findings. Our data suggest that non-canonical hosts acted as sinks for viral diversity, played limited roles in epizootic spread, and are unlikely to establish as long-term reservoirs, aligning with a model in which these species were opportunistically infected from more classical species. Test positivity data from fall 2022 showed test positivity in wild ducks of 81.8% in Newfoundland and Labrador, indicating incredibly widespread transmission in these species early on^72^. Thus, given limited resources for wildlife surveillance, our data suggest wild, migrating Anseriformes as a useful target for surveillance for risk assessment, vaccine strain selection, and tracking of viral evolution and spread.

Utilizing a large and geographically representative dataset, we estimate the number and timings of introductions in North America, and highlight the capacity of migratory birds to rapidly disseminate these viruses across the continent. Our analyses suggest that highly pathogenic H5N1 viruses may have been circulating in wild birds in North America as early as September – October of 2021, one to two months prior to the first reported detection. Following successful introduction into the Atlantic flyway, initial epizootic spread proceeded rapidly between adjacent migratory flyways, spreading from East to West within ∼4.8 months. This rapid speed of dissemination contrasts with other avian-transmitted viruses like West Nile Virus, which spread from the Northeast across the continent over the course of 4 years^46^. We speculate that the rapid geographic spread and strong signal of East to West diffusion could be explained by one or a combination of the following factors: high inherent transmissibility of clade 2.3.4.4b viruses in wild birds; rapid, long-range migration of wild bird species driving transmission; and exponential spread among an immunologically naïve population of wild birds in North America during early panzootic expansion^73,74^. Previous work has shown that within-flyway viral population structure is more observable within short time periods (< 5 years)^75^, while transmission assessed over longer time scales is more likely to span multiple flyways^76,77^. Continuous surveillance of wild species will be necessary to determine whether the rapid degree of transmission, and strong clustering by flyways observed early in the panzootic will lessen over time as immunity builds in avian hosts.

Though a single introduction into the Atlantic flyway accounted for most transmission in the epizootic, the Pacific flyway experienced more independent incursions (∼7). Compared to other studies, we infer more 5 previously undescribed incursions^78^ into the Pacific, suggesting frequent viral flow between Asia and the Pacific coast of North America that could be an important target for surveillance efforts. Despite this high degree of flow, these introductions mostly did not persist long-term. Limited transmission from the Pacific flyway could be explained by differential fitness of the lineages introduced into the Pacific vs. Atlantic flyways, ecological isolation of the Pacific flyway, or by differences in host distributions at the locations and times of these incursions. Early after introduction, the European lineage introduced into the Atlantic flyway reassorted with endemic, low-pathogenicity H5 viruses in North America, resulting in a virus that exhibited altered tissue tropism in mammals^79^. Whether this early reassortment event, or subsequent reassortments, produced viruses with enhanced fitness in wild birds is currently unknown, but could explain the success of the Atlantic flyway introduction. Alternatively, the failure of Pacific incursions to spread onward could be explained by ecological isolation of the Pacific flyway, potentially due to land features like the Rocky Mountains. Prior studies of low-pathogenicity avian influenza show a strong influence of geography on restricting virus dispersal across the North American continent, providing plausible support for this hypothesis^80–82^. Finally, Pacific flyway introductions could have persisted for less time due to a lack of suitable host species at the times and locations of the incursions, or simply by chance. Future work will be necessary to differentiate among these hypotheses.

In this study, we find that outbreaks in agriculture were seeded by repeated introductions from wild birds. This pattern held true regardless of sampling regime, and aligns with anecdotal observations that clade 2.3.4.4b viruses are increasingly being maintained by transmission among wild bird species more globally, including now in North America^17,83^. These findings contrast with genomic and epidemiologic investigation of the epizootic in 2014/2015, which implicated transmission between commercial poultry operations via personnel, equipment, and clothing as the major source of dissemination^5,84^, prompting updates to recommended biosecurity protocols^5,63^. Because the viruses circulating in 2014/2015 did not establish in local wild bird populations, that epizootic was well-controlled by culling domestic flocks, and after culling 50.5 million domestic birds, the epizootic died out. In contrast, at the time of writing, detections in wild and domestic birds in North America have continued despite culling over 160 million domestic birds. We show that despite some persistence among domestic bird populations, that introductions into domestic populations ultimately led to transmission chains that died out, and that they generally did not result in re-introduction into wild birds. These relatively short transmission chains likely reflect improved biosecurity plans developed since 2015^5^, and far more rapid and aggressive culling of domestic birds. Compared to 2014/2015, the median time from case detection to depopulation in this epizootic ranged from 4-51 hours^69^, vs. 6 days previously ^5^, indicating a far more rapid culling response. Despite these improvements in biosecurity, the number of poultry outbreaks, and corresponding human spillovers, has been staggering since 2021. Our results provide an explanation for these phenomena. Compared to past outbreaks, transmission in agricultural settings in North America since 2021 have been fundamentally distinct because these viruses circulated efficiently in wild birds, allowing for rapid dispersal and continuous outbreak re-seeding. While improvement in biosecurity may have reduced farm-to-farm spread, the establishment of continuously circulating lineages in wild birds likely made this epizootic far more challenging to prevent and control. While current US and Canadian policy classifies H5N1 as a foreign animal disease that prioritizes culling, these new dynamics challenge the effectiveness of culling as the primary strategy for control. Our data suggest that future prevention of agricultural outbreaks may now require layered interventions that seek to reduce interactions between wild and domestic birds, paired with aggressive biosecurity between farms^20,22^. As highly pathogenic H5N1 viruses continue to circulate in North American wild birds, investment in control methods that reduce successful transmission between wild birds and agricultural animals, including potentially vaccination, should be explored.

Using a small dataset from the first 6 months of the epizootic, we find phylogenetic evidence that spillovers into backyard birds may have occurred slightly earlier and more frequently than those into commercial farms. We also show through extensive subsampling experiments that outbreaks on commercial and backyard premises were generally seeded independently, with limited viral flow between them. Though commercial poultry operations generally operate with higher degrees of biosecurity than backyard flocks, there are some documented locations that could allow for interaction between commercial poultry and wildlife^34^. Retention ponds on commercial poultry farms are frequently visited by wild waterfowl^85^, while natural features such as water bodies and vegetation near residential coops and commercial production sites could also act as potential points of wild to domestic transmission. Though the role of backyard birds on avian influenza outbreaks on commercial farms is debated^65^, our data suggest that backyard birds could potentially be investigated as sentinel species for transmission in wild birds. In the United States, commercial poultry operations are more likely to be enrolled in voluntary testing programs and indemnity programs that guarantee access to funds following depopulation. As a result, commercial poultry are generally more well surveilled and have more rapid response times for testing and depopulation. Though recent data on backyard bird rearing in the US is limited, a large survey of backyard bird populations from 2004 showed that backyard bird flocks often contain multiple species, usually have outdoor access, and that 60-75% of backyard flocks experience regular contact with wild birds^34^. 38% of surveyed backyard flocks were on a property that contained a pond that attracted wild waterfowl, and 40% have a wild bird feeder on the property, providing clear, plausible links to water bird interaction. Multiple surveys have shown that biosecurity precautions tend to be much more limited in backyard bird populations, with 88% of backyard flocks using no precautions (shoe covers, footbaths, clothing changes) at all^34^. Analysis of seropositivity to avian influenza in backyard poultry in Maryland showed that exposure to waterfowl resulted in a ∼3x higher likelihood of harboring anti-influenza antibodies compared to birds not exposed to waterfowl^86^. Early infections during the 2014/2015 epizootic were recorded in backyard birds^87^, though the overall rate of backyard bird detections have been substantially higher in the current epizootic^88^, presumably due to the difference in the predominant species driving transmission (e.g., commercial poultry vs wild birds). Given backyard birds’ enhanced likelihood of interaction with wild birds, and our inference of earlier spillovers in those groups, expanded studies may be useful for investigating whether the patterns of earlier spillovers in these populations hold true more broadly. If so, then backyard bird populations could be investigated as early warning sentinels for increasing transmission in local wild bird populations. 93% of backyard flocks contain <100 birds, and most backyard bird owners report raising birds for enjoyment and eggs^34^. A subset of backyard bird owners may therefore present an opportunity for community engagement that could potentially help identify early detections. If engagement were successful, prompt reporting for illness and infections could be used to alert other nearby operations that highly pathogenic avian influenza is circulating locally, allowing for more advanced warning for heightened biosecurity.

Sampling bias is pervasive across viral outbreak datasets, and no modeling approach can completely overcome inherent biases in data acquisition. In this study, we employed multiple, overlapping analyses to control for sampling biases, providing a framework for performing phylodynamic analyses in the presence of uncertain sampling. Highly pathogenic avian influenza data is rife with sampling issues. In the US, avian influenza sampling of wild birds employs 3 distinct methods (hunter harvest, sick and dead, and live birds), each with unmeasured biases. Only wild Anseriformes are sampled live or hunter harvested, while all other host groups are sampled sick or dead, likely skewing detections towards birds with dedicated rescue services and those located near humans. In domestic birds, detection depends on producer identification of illness and reporting, and subsequent testing, which likely varies across production types, locations, and premises. While we accumulated as much data as possible to contextualize how cases are sampled and sequenced in North America, it is impossible to know the true case burden in either domestic or wild birds. Because of this, we opted to investigate multiple distinct subsampling approaches (equal and proportional), report results that are consistent between them, and to employ statistical tests for trait bias to most accurately estimate transmission dynamics. Equal sampling regimes rely inherently on the diversity in sampled sequences to make inferences about sources of viral diversity, while proportional regimes bring the data in line with model assumptions that sequence numbers are proportional to cases. For analyses of host orders and wild/domestic bird transitions (via titration tests), we employ both. Beginning with an equal sampling regime, we show that the genetic diversity of viruses circulating in Anseriformes and wild birds is sufficient to infer them as source populations, patterns which remained under proportional sampling regimes. In the titration tests, we show that while the central conclusions remained true, that the precise number of transitions between wild and domestic birds depends heavily on sampling numbers, providing a clear argument for continuous surveillance in wildlife, and a warning for over-confidence in estimating the particular numbers of transitions between groups. Finally, we quantified the level of imbalance between sampled traits across each discrete trait analysis, and find limited evidence that sampling biases drove our inferences (Table S17-18). Still, these methods have important caveats. An equal subsampling regime might over-represent a truly under-impacted group, while a proportional subsampling might over-represent a more well sampled group. Though use of replicates allowed us greater confidence in our results, all phylodynamic inferences are limited by sequence data availability, and results could change if future data become available. Additional sequence data could result in inference of more introductions into North America, or altered numbers of transitions between domestic and wild birds. Our analyses only employ HA sequences, a caveat that could result in slightly different numbers of inferred transitions if full genomes were used. Accurately distinguishing between hypotheses of epizootic spread (e.g., whether agricultural outbreaks are driven by introductions from wild birds or by from farm-to-farm spread) depends on adequate sequence data from wild birds, without which transmission inference is impossible. As H5N1 viruses continue to evolve and spread globally, investment in surveillance strategies that capture circulating diversity among wild birds will likely be critical for accurately tracking viral evolution, prioritizing vaccine strains, and contextualizing new emergence events, like the recent outbreaks in dairy cattle.

Taken together, our findings implicate a critical shift in highly pathogenic avian influenza ecology in North America, with wild birds playing the central role in transmission and dispersal in the 2021-2023 epizootic. Persistent and intensive transmission in wild birds provides an explanation for the rapid cross-continental spread, and continued agricultural outbreaks despite aggressive culling. Our results highlight the utility of wild bird surveillance for accurately distinguishing hypotheses of epizootic spread, and suggest continuous surveillance as critical for preventing and dissecting future outbreaks. Our data underscore that continued establishment of H5N1 in North American wildlife may necessitate a shift in risk management and mitigation, with interventions focused on reducing risk within the context of enzootic circulation in wild birds. At the time of writing, outbreaks in dairy cattle highlight the critical importance of modeling the ecological interactions within and between wild birds and domestic production. Future work to effectively model viral evolution and spread hinges critically on effective surveillance across wild and domestic species to capture key transmission pathways across large geographic scales. Ultimately, these data are essential for informing biosecurity, outbreak response, and vaccine strain selection.

## Materials and Methods

### Dataset collection and processing

#### Genomic data processing and initial phylogenetics

We downloaded all available nucleotide sequence data and associated meta-data for the Hemagglutinin protein of all HPAI clade 2.3.4.4b H5Nx viruses from the GISAID database on 2023-11-25^89^. For each subset of the data described for further phylodynamic modeling the following process was followed. We first aligned sequences using MAFFT v7.5.20, sequence alignments were visually inspected using Geneious and sequences causing significant gaps were removed and nucleotides before the start codon and after the stop codon were removed^90,91^. We de-duplicated identical sequences collected on the same day (retaining identical sequences that occurred on different days). We identified and removed temporal outliers for all genomic datasets by performing initial phylogenetic reconstruction in a maximum likelihood framework using IQtree v.1.6.12 and used the program TimeTree v 0.11.2 was used to remove temporal outliers and to assess the clockliness of the dataset prior to Bayesian phylogenetic reconstruction^92,93^. This resulted in a dataset of 1824 sequences that were used in further analyses (Figure S24).

#### AVONET database

We downloaded the AVONET database for avian ecology data and merged it to available host metadata from GISAID for each sequence^48^. We used the species if provided to match the species indicated in the AVONET database. If host metadata in GISAID was defined using common name for a bird, we determined the taxonomic species name and used that for further merging with the AVONET data (e.g. “Mallard” was replaced with *Anas platyrhynchos*) for the given region to match the species to its respective ecological data. Domesticity status (whether a sequence was isolated from a wild host or a domestic host) was determined using available metadata downloaded from GISAID using the ‘Note’ and ‘Domestic_Status’ fields in sequence associated metadata. Additionally, if a given sequence strain name (in the field ‘Isolate_Name’) indicated domestic status (e.g. A/domestic_duck/2022) these sequences were labeled as belonging to domestic hosts.

#### Detections by USDA

Data for detections of HPAI in North America were collected from USDA APHIS. Reports for mammals, wild birds, and domestic poultry were all downloaded (download date: 2023-11-25)^60^.

### Phylodynamic analysis

The following Bayesian phylogenetic reconstructions and analyses were performed using BEAST v.1.10.4^94^.

#### Empirical tree set estimation and coalescent analysis

We performed Bayesian phylogenetic reconstruction for each dataset prior to discrete trait diffusion modeling to estimate a posterior set of empirical trees. The following priors and settings were used for each subset of the sequence data. We used the HKY nucleotide substitution model with gamma-distributed rate variation among sites and lognormal relaxed molecular clock model^95,96^. The Bayesian SkyGrid coalescent was used with the number of grid points corresponding to the number of weeks between the earliest and latest collected sample (e.g for a dataset collected between 2021-11-04 and 2023-08-11 we would set 92 grid points)^40^. We initially ran four independent MCMC chains with a chain-length of 100 million states logging every 10000 states. We diagnosed the combined results of the independent runs diagnosed Tracer v1.7.2. to ensure adequate ESS (ESS > 200) and reasonable estimates for parameters^94^. If ESS was inadequate additional independent MCMC runs were run increasing chain length to 150 million states, sampling every 15000 states were performed. We combined the tree files from each independent MCMC run removing 10-30% burn-in and resampling to get a tree file with between 9000 and 10000 posterior trees using Logcombiner v1.10.4. A posterior sample of 500 trees was extracted and used as empirical tree sets in discrete trait diffusion modeling.

### Discrete trait diffusion analysis

#### Dataset subsampling and definition of discrete traits

We characterized the geographic introduction of HPAI into North America by randomly sampling 100 sequences from Europe and Asia for each year between 2021-2023 (total 300 non-North American) and all available North American sequences across the study period. Following removal of temporal outliers this resulted in a dataset of n= 1927 sequences annotated by continent of origin. The sequence data available from North America broken down by country is as follows: United States (1590), Canada (224), Honduras (2), Costa Rica (5), and Panama (1).

To characterize geographic transmission within North America, following introduction, we constructed a dataset of sequences subsampled based on migratory flyway. We used place of isolation data to match the US state or Canadian province the sequence was collected from with the respective U.S. Fish and Wildlife Service Migratory Bird Program Administrative Flyway^44^. We subsampled 250 sequences for each flyway (Atlantic, Mississippi, Central, and Pacific) to create a dataset of 1000 sequences collected between November 2021 and August 2023. In addition to USFWS flyways we defined 4 geographic regions going North to South based on latitude lines with the following delineations for each group. We divided North America into 4 regions segregated by latitude, with the northernmost group above the 49N parallel and the southernmost group below the 36N parallel. We then sampled 916 sequences eveningly across these categories and inferred transitions between these regions.

We classified sequences by host taxonomic order, inferring the host species using designations in the strain name and/or metadata to match species records in AVONET^48^. To ensure that each discrete trait had an adequate number of samples for the discrete trait analysis of host orders we combined orders in two instances based on taxonomic and behavioral similarity. The order Falconiformes (n=14), which represents falcons, was added to Accipitriformes (n=363), which includes other raptors such as eagles, hawks, and vultures. Pelecaniformes (n=34) which includes pelicans were grouped with Charadriiformes (n = 74, shorebirds and waders) due to their similar aquatic lifestyles and behaviors. Mammals were kept as a broad non-human classification as most samples were of the order carnivora (foxes, skunks, bobcats etc.), apart from samples of dolphins (Artiodactyla) and Virginia opossum (Didelphimorphia). The following orders were omitted due to low number of sequences: Rheaforimes (n=2), Casuariiformes (n=1), Apodiformes (n=2), Suliformes (n=7), Gaviiformes (n=1), Gruiformes (n=1), Podicipediformes (n=1).

We randomly subsampled 100 sequences for each host order between 2021-11-04 and 2023-08-11 resulting in a dataset of n=655 sequences where all isolates for host orders with less than 100 samples, Passeriformes (n = 57) and Strigiformes (n=99) (removing one temporal outlier), were used (Figure S25). We repeated this random subsampling three times resulting in three separate datasets. We additionally performed three sub-samples of sequences based on the proportion of detections in each host order group which were collected between 2021-11-04 and 2023-08-11. Three random proportional samples were taken each with the following number of sequences for each group: Accipitriformes =133, Anseriformes = 342, Passeriformes = 12, Nonhuman-mammal = 16, Galliformes = 83, Charadriiformes = 40, Strigiformes = 29 (total n=655 sequences).

We defined discrete traits for use in discrete trait diffusion modeling based on the available sequence metadata and merged AVONET data. In addition to taxonomic order, we defined migratory behavior. Birds were classified as sedentary (staying in each location and not showing any major migration behavior), Partially migratory (e.g. small proportion of population migrates long distances, or population undergoes short-distance migration, nomadic movements, distinct altitudinal migration, etc.), or Migratory (majority of population undertakes long-distance migration). We subsampled sequences based on migratory behavior including nonhuman-mammals and domestic birds to create a subsample of 500 sequences with equal sampling across behavior groups.

#### Discrete trait modeling framework

For each discrete trait dataset, we used an asymmetric continuous time Markov chain discrete trait diffusion model and implemented the Bayesian stochastic search variable selection (BSSVS) to determine the most parsimonious diffusion network^41^. We inferred the history of changes from a given trait to another across branches of the phylogeny, providing a rate of transitions from A to B/year for each pair of trait states. When reporting these results, we refer to state A as the source population/state and B as the sink population/state. We implemented the Bayesian Stochastic Search Variable Selection which allows us to determine which rates have the highest posterior support by using a stochastic binary operator which turns on and off rates to determine their contribution to the diffusion network. In addition to the discrete trait diffusion rate employed a Markov Jump analysis to observe the number of jumps between discrete states across the posterior set of trees and estimated the Markov Rewards to determine the waiting time for a given discrete trait state in the phylogeny^66,67^. We calculate the transition rate as a realization of the CTMC process by dividing the number of markov jumps by the tree height (branch length from the earliest tip to the root of the tree), and separately, by tree length (sum of all branch lengths). For each pairwise transition rate, we calculate the level of Bayes Factor (BF) support that the given rate has. The BF represents the support of a given rate. The BF is calculated as the ratio of the posterior odds of the given rate being non-zero divided by the equivalent prior odds which is set as a Poisson prior with a 50% prior probability on the minimal number of rates possible^41^. We use the support definitions by Kass and Rafferty to interpret the BF support where a BF > 3 indicates little support, a BF between 3 and 10 indicates substantial support, a BF between 10 and 100 indicates strong support, and a BF greater than 100 indicates very strong support^97^.

Empirical sets were used with the discrete traits defined for each sequence to perform discrete trait diffusion modeling. Each discrete trait model was implemented using three independent MCMC chains with a chain length of 10 million states, logging every 1000 states. Runs were combined using LogCombiner v.1.10.4. subsampling a posterior sample of 10,000 trees/states. The Bayes Factor support for transition rates were calculated using the program SPREAD3 ^98^. Maximum clade credibility trees were constructed using TreeAnnotator v1.10.4.

#### Domestic/Wild titration analysis

To study the impact of sampling of wild birds on the estimation of rates between domestic and wild birds we created five separate datasets with varying numbers of wild birds for sequences collected between 2021 and 2023. We randomly sampled 270 domestic sequences and 270 wild sequences as the initial 1:1 ratio dataset. We then made four more datasets increasing the number of wild sequences by a factor of 0.5 (adding 135 wild sequences) resulting in a final “titration” of 1:3 domestic to wild sequences (n=1080). We applied a two-state asymmetric CTMC discrete trait diffusion model where sequences were labeled as domestic or wild. All priors and model parameters selected are the same as those described in the empirical tree set description above. To study the impact of the inclusion of turkeys in the transmission between domestic and wild populations we annotated all unannotated sequences collected from turkeys as domestic. We then created three datasets starting with 525 domestic and 525 wild bird sequences, adding 263 sequences to successive titrations resulting in 1:1,1:1.5, and 1:2 (domestic:wild) sequence datasets with a final titration size of 1,575 sequences. We again applied a two-state asymmetric CTMC discrete trait diffusion model where sequences were labeled as domestic or wild with all priors and model parameters selected are the same as those described in the empirical tree set description above. To determine whether the proportion of turkeys to other domestic birds would impact the results of the previously described titration analysis we built a dataset with where the domestic bird group had equal numbers of turkey and domestic (non-turkey) sequences. This dataset included 173 turkey, 173 domestic bird, and 692 wild bird sequences totaling 1038 sequences. We applied an asymmetric CTMC discrete trait diffusion model using a BSSVS for a three-trait model with the following states: wild birds, domestic birds (not turkey), and turkey. We performed three independent runs of this analysis using the models and parameters described in the empirical tree analysis section above. All titration replicates were performed using an MCMC chain length of 100 million states sampling every 10,000 states.

#### Commercial, backyard, wild bird titration analysis

Metadata and annotated sequences were made available describing sequences as being from backyard birds for sequences collected in early 2022 which distinguished them from commercial poultry (previously all sequences being determined domestic)^10^. We used this metadata to create a dataset with equally sampled backyard birds and commercial birds (n= 85 for each bird type) and then added all available wild birds (n=722) in 25% increments creating four separate datasets for sequences collected between Jan 2022 and June 2022. This resulted in a final dataset of n= 942 sequences. We performed discrete trait diffusion modeling using an asymmetric CTMC diffusion model described in the previous section for sequences labeled as backyard bird, commercial bird, and wild bird.

#### Extraction of phylogenetic metrics

We calculated the transitions between states across branches of phylogenies estimated from ancestral state reconstructions using the Baltic python package^28^. To calculate the persistence of a given discrete trait we used the program PACT v0.9.5. which calculates the persistence of a trait by traversing the phylogenetic tree backwards and measuring the amount of time a tip takes to leave its sampled state^99^.

### Assessment of Sampling Bias

#### BaTs analysis

To determine if the discrete traits analyzed correlated with shared ancestry in the phylogeny, we employed tip trait association tests implemented in the Bayesian Tip-association Significance (BaTs) program v1.0^45^. This program assesses the phylogenetic structure of discrete traits across viral lineages using three metrics: the association index (AI), parsimony score (PS), and (maximum monophyletic clade size (MC). The AI measures the imbalance of internal nodes of a phylogeny for a given set of traits. The PS calculates the number of state changes in the phylogeny. The MC measures the maximum number of tips belonging to a monophyletic clade for each discrete trait of interest. These metrics are calculated for the phylogeny as tips are randomly swapped to create a null distribution to compare against. Taken together these metrics quantify the degree of clustering within the phylogeny with Lower AI and PS values indicating stronger phylogenetic structure, suggesting that closely related taxa tend to share the same trait, whereas higher values indicate weaker structure and more frequent transitions between trait states. Statistical significance was assessed by comparing observed values against a null distribution generated through randomization, with p-values reported for each test. All discrete trait groupings showed evidence for clustering by trait, supporting the use of trait modeling across the tree.

#### Tip shuffle analysis

To assess the sensitivity of each of our discrete trait reconstructions to differences in sampling between groups, we implemented a modified version of a tip swap analysis^55^. As originally developed, a tip swap analysis attempts to assess the impact of trait sampling on discrete trait measurements. An operator is implemented within the MCMC chain that randomly picks pairs of tips and swaps their trait values, thus generating a posterior set of trees among which pairs of trait assignments have been randomly swapped. The probability of each state at the root is then computed, and compared to the inferred root state probabilities in the empirical data. Because the root state probabilities in randomized datasets should primarily reflect the frequency of each trait in the analysis, empirical results that differ substantially from this null distribution are interpreted as evidence that the sequence data is informing the analysis beyond what is expected based on trait frequencies alone. Thus, traits for which the root state probability differs considerably from the root state probability in the null data are frequently interpreted as being informed by the data, rather than sampling bias. While this approach has been shown to perform well on small phylogenies^50,77^, the strategy of swapping single pairs of tips poses challenges for larger trees. In our flyways dataset which includes ∼1000 tips, we found that even with extremely high operator values (4000), the traditional tip-swap analysis resulted in a posterior set of trees in which the majority of tips (93.3%) remained assigned to their true state at least 50% of the time, resulting in a null dataset that was only partially randomized. We believe this is due to the high number of tips in our analysis, resulting in only an extremely small fraction of tips randomized at any given step in the MCMC chain. To overcome this limitation, we instead performed a randomized tip shuffle analysis. Using the empirical set of trees inferred for each discrete trait analysis, we generated 100 null datasets in which we shuffled the trait assignments randomly across the tips. In this approach, we preserve the phylogenetic tree topology and the ratio of samples from each group, but shuffle their assignments at the tips. For each discrete trait analysis, we generated 100 distinct shuffled versions of the empirical trees, reran the analysis, and summarized the resulting posterior distribution by inferring a maximum clade credibility tree. We then computed the root state probabilities for each trait for each mcc tree, and computed the mean root state probability across all 100 replicates. This computed mean is reported in Table S19, in the column labelled “Mean root state probability across 100 datasets with randomly shuffled tip states.**”** Finally, we compared these null values to the root state probabilities calculated for each group in the empirical data (reported as “Root stat probability in Empirical data” in Table S19). As expected, the root state probabilities inferred in the shuffled datasets are proportional to the number of sequences included for each group. For the analyses using an equal sampling regime (Migration, Flyway, Host Orders Equal, and initial titration tests), this leads to approximately equal expected root state probabilities across groups. In contrast, the root state probabilities in the empirical data generally differ significantly from expectation, suggesting that the phylogenetic results are informed by the genetic data rather than from sampling alone.

The results of BaTS analyses and tip shuffling analyses for each discrete trait in this study can be found in the supplemental material (Figure S30-S39), Table S18-19).

## Data and code availability

All analytical scripts, metadata annotations, and BEAST XMLs used in this analysis can be found at the following GitHub repository: https://github.com/moncla-lab/North-American-HPAI

All data that was used in this analysis were sourced from public databases. Acknowledgement table for GISAID isolates used in this analysis can be found in Table S20.

Several of the analyses presented have also been publicly made available using a maximum likelihood framework through the Nextstrain pipeline and a narrative of this work can be found in the following link: https://nextstrain.org/community/narratives/moncla-lab/nextstrain-narrative-hpai-north-america@main/HPAI-in-North-America

## Acknowledgements

We would like to thank Mia Kim Torchetti for her helpful feedback and discussion about our results. This work was supported by NIH R00-AI147029-05 and by funding from the Centers of Excellence for Influenza Research and Response (CEIRR), funded by NIH 75N93021C00015. LHM is a Pew Biomedical Scholar and is supported by NIH R00-AI147029-05. LD is supported by NIH 75N93021C00015.

## Citations

1. Puryear, W. B. & Runstadler, J. A. High-pathogenicity avian influenza in wildlife: a changing disease dynamic that is expanding in wild birds and having an increasing impact on a growing number of mammals. J. Am. Vet. Med. Assoc. 1–9 (2024) doi:10.2460/javma.24.01.0053.

2. Lycett, S. J., et al. Genesis and spread of multiple reassortants during the 2016/2017 H5 avian influenza epidemic in Eurasia. Proc. Natl. Acad. Sci. 117, 20814–20825 (2020).

3. Smith, G. J. D., et al. Characterization of Avian Influenza Viruses A (H5N1) from Wild Birds, Hong Kong, 2004–2008. Emerg. Infect. Dis. 15, 402–407 (2009).

4. Alexander, D. J. Summary of Avian Influenza Activity in Europe, Asia, Africa, and Australasia, 2002–2006. Avian Dis. 51, 161–166 (2007).

5. Final Report for the 2014–2015 Outbreak of Highly Pathogenic Avian Influenza (HPAI) in the United States. https://www.aphis.usda.gov/media/document/2086/file (2016).

6. Caliendo, V., et al. Transatlantic spread of highly pathogenic avian influenza H5N1 by wild birds from Europe to North America in 2021. Sci. Rep. 12, 11729 (2022).

7. Gass, J. D., et al. Ecogeographic Drivers of the Spatial Spread of Highly Pathogenic Avian Influenza Outbreaks in Europe and the United States, 2016–Early 2022. Int. J. Environ. Res. Public. Health 20, 6030 (2023).

8. Erdelyan, C. N. G., et al. Multiple transatlantic incursions of highly pathogenic avian influenza clade 2.3.4.4b A(H5N5) virus into North America and spillover to mammals. Cell Rep. 43, 114479 (2024).

9. Klaassen, M. & Wille, M. The plight and role of wild birds in the current bird flu panzootic. Nat. Ecol. Evol. 7, 1541–1542 (2023).

10. Youk, S., et al. H5N1 highly pathogenic avian influenza clade 2.3.4.4b in wild and domestic birds: Introductions into the United States and reassortments, December 2021–April 2022. Virology 587, 109860 (2023).

11. Spackman, E., Pantin-Jackwood, M. J., Lee, S. A. & Prosser, D. The pathogenesis of a 2022 North American highly pathogenic clade 2.3.4.4b H5N1 avian influenza virus in mallards (*Anas platyrhynchos*). Avian Pathol. 52, 219–228 (2023).

12. Knief, U., et al. Highly pathogenic avian influenza causes mass mortality in Sandwich Tern *Thalasseus sandvicensis* breeding colonies across north-western Europe. Bird Conserv. Int. 34, e6 (2024).

13. European Food Safety Authority et al. Avian influenza overview December 2022 – March 2023. EFSA J. 21, (2023).

14. Elsmo, E., et al. Pathology of natural infection with highly pathogenic avian influenza virus (H5N1) clade 2.3.4.4b in wild terrestrial mammals in the United States in 2022. Preprint at 10.1101/2023.03.10.532068 (2023).

15. Nguyen, T.-Q. et al. Emergence and interstate spread of highly pathogenic avian influenza A(H5N1) in dairy cattle. Preprint at 10.1101/2024.05.01.591751 (2024).

16. Puryear, W. et al. Outbreak of Highly Pathogenic Avian Influenza H5N1 in New England Seals. Preprint at 10.1101/2022.07.29.501155 (2022).

17. Xie, R., et al. The episodic resurgence of highly pathogenic avian influenza H5 virus. Nature 622, 810–817 (2023).

18. Wille, M. Ecology and Evolution of Avian Influenza A Viruses in Wild Birds. in Genetics and Evolution of Infectious Diseases 863–898 (Elsevier, 2024). doi:10.1016/B978-0-443-28818-0.00005-7.

19. Kaplan, B. S. & Webby, R. J. The avian and mammalian host range of highly pathogenic avian H5N1 influenza. Virus Res. 178, 3–11 (2013).

20. Hill, N. J., et al. Ecological divergence of wild birds drives avian influenza spillover and global spread. PLOS Pathog. 18, e1010062 (2022).

21. Sonnberg, S., Webby, R. J. & Webster, R. G. Natural history of highly pathogenic avian influenza H5N1. Virus Res. 178, 63–77 (2013).

22. Verhagen, J. H., Fouchier, R. A. M. & Lewis, N. Highly Pathogenic Avian Influenza Viruses at the Wild–Domestic Bird Interface in Europe: Future Directions for Research and Surveillance. Viruses 13, 212 (2021).

23. Alkie, T. N., et al. A threat from both sides: Multiple introductions of genetically distinct H5 HPAI viruses into Canada via both East Asia-Australasia/Pacific and Atlantic flyways. Virus Evol. 8, veac077 (2022).

24. HIGHLY PATHOGENIC AVIAN INFLUENZA RESPONSE PLAN THE RED BOOK. https://www.aphis.usda.gov/sites/default/files/hpai_response_plan.pdf.

25. Policy on the Importation of Terrestrial Foreign or Emerging Animal Disease Agents into Canada by External Facilities. https://inspection.canada.ca/en/animal-health/requirements-handling-pathogens/pathogen-imports/policy-importation?utm_source=chatgpt.com.

26. Volz, E. M., Koelle, K. & Bedford, T. Viral Phylodynamics. PLoS Comput. Biol. 9, e1002947 (2013).

27. Frost, S. D. W. & Volz, E. M. Viral phylodynamics and the search for an ‘effective number of infections’. Philos. Trans. R. Soc. B Biol. Sci. 365, 1879–1890 (2010).

28. Dudas, G., Carvalho, L. M., Rambaut, A. & Bedford, T. MERS-CoV spillover at the camel-human interface. eLife 7, e31257 (2018).

29. De Maio, N., Wu, C.-H., O’Reilly, K. M. & Wilson, D. New Routes to Phylogeography: A Bayesian Structured Coalescent Approximation. PLOS Genet. 11, e1005421 (2015).

30. Overview of How Canada Prevents, Prepares and Responds to Bird Flu Outbreaks. https://inspection.canada.ca/en/animal-health/terrestrial-animals/diseases/reportable/avian-influenza/prevention-preparedness-and-response.

31. Bevins, S. N., et al. Large-Scale Avian Influenza Surveillance in Wild Birds throughout the United States. PLoS ONE 9, e104360 (2014).

32. Shriner, S. A., et al. Surveillance for highly pathogenic H5 avian influenza virus in synanthropic wildlife associated with poultry farms during an acute outbreak. Sci. Rep. 6, 36237 (2016).

33. Bokma, B. H., Hall, C., Siegfried, L. M. & Todd Weaver, J. Surveillance for Avian Influenza in the United States. Ann. N. Y. Acad. Sci. 1081, 163–168 (2006).

34. Poultry 2004 Part I: Reference of Health and Management of Backyard / Small Production Flocks in the United States, 2004. https://www.aphis.usda.gov/sites/default/files/poultry04_dr_parti.pdf.

35. Garber, L., Hill, G., Rodriguez, J., Gregory, G. & Voelker, L. Non-commercial poultry industries: Surveys of backyard and gamefowl breeder flocks in the United States. Prev. Vet. Med. 80, 120–128 (2007).

36. 2022 OIE - Terrestrial Animal Health Code. https://www.woah.org/fileadmin/Home/eng/Health_standards/tahc/current/glossaire.pdf (2022).

37. McKellar, A. E. Phenological Synchrony and Bird Migration: Changing Climate and Seasonal Resources in North America Phenological Synchrony and Bird Migration: Changing Climate and Seasonal Resources in North America edited by Eric M. Wood and Jherime L. Kellermann. 2015. CRC Press, Boca Raton, Florida, USA. xiv + 228 pages, 8 color and 53 black-and-white illustrations. $116.96 (hardcover). ISBN 978-1-4822-4030-6. *The Auk* 133, 113–114 (2016).

38. Marra, P. P., Francis, C. M., Mulvihill, R. S. & Moore, F. R. The influence of climate on the timing and rate of spring bird migration. Oecologia 142, 307–315 (2005).

39. Kent, C. M., et al. Spatiotemporal changes in influenza A virus prevalence among wild waterfowl inhabiting the continental United States throughout the annual cycle. Sci. Rep. 12, 13083 (2022).

40. Gill, M. S., et al. Improving Bayesian Population Dynamics Inference: A Coalescent-Based Model for Multiple Loci. Mol. Biol. Evol. 30, 713–724 (2013).

41. Lemey, P., Rambaut, A., Drummond, A. J. & Suchard, M. A. Bayesian Phylogeography Finds Its Roots. PLoS Comput. Biol. 5, e1000520 (2009).

42. Ramey, A. M., et al. Molecular detection and characterization of highly pathogenic H5N1 clade 2.3.4.4b avian influenza viruses among hunter-harvested wild birds provides evidence for three independent introductions into Alaska. Virology 589, 109938 (2024).

43. Krammer, F., Hermann, E. & Rasmussen, A. L. Highly pathogenic avian influenza H5N1: history, current situation, and outlook. J. Virol. 99, e02209–24 (2025).

44. Migratory Bird Program Administrative Flyways | U.S. Fish & Wildlife Service. https://www.fws.gov/partner/migratory-bird-program-administrative-flyways (2023).

45. Parker, J., Rambaut, A. & Pybus, O. G. Correlating viral phenotypes with phylogeny: Accounting for phylogenetic uncertainty. Infect. Genet. Evol. 8, 239–246 (2008).

46. Kramer, L. D., Ciota, A. T. & Kilpatrick, A. M. Introduction, Spread, and Establishment of West Nile Virus in the Americas. J. Med. Entomol. 56, 1448–1455 (2019).

47. Gergely, K.J., Boykin, K.G., McKerrow, A.J., Rubino, M.J., Tarr, N.M., and Williams, S.G. Gap Analysis Project (GAP) Terrestrial Vertebrate Species Richness Maps for the Conterminous U.S. 10.3133/sir20195034. (2019).

48. Tobias, J. A., et al. AVONET: morphological, ecological and geographical data for all birds. Ecol. Lett. 25, 581–597 (2022).

49. Arnal, A., et al. Laridae: A neglected reservoir that could play a major role in avian influenza virus epidemiological dynamics. Crit. Rev. Microbiol. 41, 508–519 (2015).

50. Lee, D.-H., et al. Transmission Dynamics of Highly Pathogenic Avian Influenza Virus A(H5Nx) Clade 2.3.4.4, North America, 2014–2015. Emerg. Infect. Dis. 24, 1840–1848 (2018).

51. Munster, V. J. et al. Spatial, Temporal, and Species Variation in Prevalence of Influenza A Viruses in Wild Migratory Birds. PLoS Pathog. 3, e61 (2007).

52. Kwon, J., et al. Domestic ducks play a major role in the maintenance and spread of H5N8 highly pathogenic avian influenza viruses in South Korea. Transbound. Emerg. Dis. 67, 844–851 (2020).

53. Dhingra, M. S., et al. Global mapping of highly pathogenic avian influenza H5N1 and H5Nx clade 2.3.4.4 viruses with spatial cross-validation. eLife 5, e19571 (2016).

54. Hong, S. L., Lemey, P., Suchard, M. A. & Baele, G. Bayesian Phylogeographic Analysis Incorporating Predictors and Individual Travel Histories in BEAST. Curr. Protoc. 1, e98 (2021).

55. Edwards, C. J., et al. Ancient Hybridization and an Irish Origin for the Modern Polar Bear Matriline. Curr. Biol. 21, 1251–1258 (2011).

56. Nemeth, N. M., et al. Bald eagle mortality and nest failure due to clade 2.3.4.4 highly pathogenic H5N1 influenza a virus. Sci. Rep. 13, 191 (2023).

57. Bahl, J., et al. Ecosystem Interactions Underlie the Spread of Avian Influenza A Viruses with Pandemic Potential. PLOS Pathog. 12, e1005620 (2016).

58. Shearn-Bochsler, V. I., Knowles, S. & Ip, H. Lethal Infection of Wild Raptors with Highly Pathogenic Avian Influenza H5N8 and H5N2 Viruses in the USA, 2014–15. J. Wildl. Dis. 55, 164 (2019).

59. Redig, P. T. & Goyal, S. M. Serologic Evidence of Exposure of Raptors to Influenza A Virus. Avian Dis. 56, 411–413 (2012).

60. 2022–2024 Detections of Highly Pathogenic Avian Influenza. https://www.aphis.usda.gov/livestock-poultry-disease/avian/avian-influenza/hpai-detections.

61. Farahat, R. A., Khan, S. H., Rabaan, A. A. & Al-Tawfiq, J. A. The resurgence of Avian influenza and human infection: A brief outlook. New Microbes New Infect. 53, 101122 (2023).

62. Patyk, K. A., et al. Investigation of risk factors for introduction of highly pathogenic avian influenza H5N1 infection among commercial turkey operations in the United States, 2022: a case-control study. Front. Vet. Sci. 10, 1229071 (2023).

63. APHIS FOREIGN ANIMAL DISEASE FRAMEWORK RESPONSE STRATEGIES. https://www.aphis.usda.gov/sites/default/files/fadprep_manual_2.pdf.

64. APPA NATIONAL PET OWNERS SURVEY 2021-2022.

65. Smith, G. & Dunipace, S. How backyard poultry flocks influence the effort required to curtail avian influenza epidemics in commercial poultry flocks. Epidemics 3, 71–75 (2011).

66. Minin, V. N. & Suchard, M. A. Counting labeled transitions in continuous-time Markov models of evolution. J. Math. Biol. 56, 391–412 (2007).

67. Minin, V. N. & Suchard, M. A. Fast, accurate and simulation-free stochastic mapping. Philos. Trans. R. Soc. B Biol. Sci. 363, 3985–3995 (2008).

68. FOIA Report 24-04961-F_000001. 24-04961-F_000001 https://www.aphis.usda.gov/sites/default/files/avian-influenza-incidents-and-depopulation-methods-feb-2022-to-june-2024.pdf.

69. 2022–2023 HIGHLY PATHOGENIC AVIAN INFLUENZA OUTBREAK Summary of Depopulation Methods and the Impact on Lateral Spread. https://www.aphis.usda.gov/sites/default/files/hpai-2022-2023-summary-depop-analysis.pdf.

70. Census of Agriculture. https://www.nass.usda.gov/Publications/AgCensus/2022/#full_report (2022).

71. Hayes, S. et al. Ecology and environment predict spatially stratified risk of highly pathogenic avian influenza in wild birds across Europe. Preprint at 10.1101/2024.07.17.603912 (2024).

72. Wight, J., et al. Avian influenza virus circulation and immunity in a wild urban duck population prior to and during a highly pathogenic H5N1 outbreak. Vet. Res. 55, 154 (2024).

73. Graziosi, G., Lupini, C., Catelli, E. & Carnaccini, S. Highly Pathogenic Avian Influenza (HPAI) H5 Clade 2.3.4.4b Virus Infection in Birds and Mammals. Animals 14, 1372 (2024).

74. Stallknecht, D. E., et al. Highly Pathogenic H5N1 Influenza A Virus (IAV) in Blue-Winged Teal in the Mississippi Flyway Is Following the Historic Seasonal Pattern of Low-Pathogenicity IAV in Ducks. Pathogens 13, 1017 (2024).

75. Bahl, J., et al. Influenza A Virus Migration and Persistence in North American Wild Birds. PLoS Pathog. 9, e1003570 (2013).

76. Fourment, M., Darling, A. E. & Holmes, E. C. The impact of migratory flyways on the spread of avian influenza virus in North America. BMC Evol. Biol. 17, 118 (2017).

77. Prosser, D. J., et al. Maintenance and dissemination of avian-origin influenza A virus within the northern Atlantic Flyway of North America. PLOS Pathog. 18, e1010605 (2022).

78. Ahlstrom, C. A., et al. Genomic characterization of highly pathogenic H5 avian influenza viruses from Alaska during 2022 provides evidence for genotype-specific trends of spatiotemporal and interspecies dissemination. Emerg. Microbes Infect. 13, 2406291 (2024).

79. Kandeil, A., et al. Rapid evolution of A(H5N1) influenza viruses after intercontinental spread to North America. Nat. Commun. 14, 3082 (2023).

80. Lam, T. T., et al. Migratory flyway and geographical distance are barriers to the gene flow of influenza virus among North American birds. Ecol. Lett. 15, 24–33 (2012).

81. Krauss, S., et al. Influenza in Migratory Birds and Evidence of Limited Intercontinental Virus Exchange. PLoS Pathog. 3, e167 (2007).

82. Girard, Y. A., Runstadler, J. A., Aldehoff, F. & Boyce, W. Genetic structure of Pacific Flyway avian influenza viruses is shaped by geographic location, host species, and sampling period. Virus Genes 44, 415–428 (2012).

83. Yang, Q., et al. Synchrony of Bird Migration with Global Dispersal of Avian Influenza Reveals Exposed Bird Orders. Nat. Commun. 15, 1126 (2024).

84. Hicks, J. T., et al. Agricultural and geographic factors shaped the North American 2015 highly pathogenic avian influenza H5N2 outbreak. PLOS Pathog. 16, e1007857 (2020).

85. Sullivan, J. D., McDonough, A. M., Lescure, L. M. & Prosser, D. J. Identifying an Understudied Interface: Preliminary Evaluation of the Use of Retention Ponds on Commercial Poultry Farms by Wild Waterfowl. Transbound. Emerg. Dis. 2024, 1–9 (2024).

86. Madsen, J. M., Zimmermann, N. G., Timmons, J. & Tablante, N. L. Avian Influenza Seroprevalence and Biosecurity Risk Factors in Maryland Backyard Poultry: A Cross-Sectional Study. PLoS ONE 8, e56851 (2013).

87. Bertran, K., et al. Pathobiology of Clade 2.3.4.4 H5Nx High-Pathogenicity Avian Influenza Virus Infections in Minor Gallinaceous Poultry Supports Early Backyard Flock Introductions in the Western United States in 2014-2015. J. Virol. 91, e00960–17 (2017).

88. Epidemiologic and Other Analyses of HPAI-Affected Poultry Flocks. https://www.aphis.usda.gov/sites/default/files/epi-analyses-avian-flu-poultry-2nd-interim-rpt.pdf.

89. Shu, Y. & McCauley, J. GISAID: Global initiative on sharing all influenza data – from vision to reality. Eurosurveillance 22, (2017).

90. Katoh, K. & Standley, D. M. MAFFT Multiple Sequence Alignment Software Version 7: Improvements in Performance and Usability. Mol. Biol. Evol. 30, 772–780 (2013).

91. Kearse, M., et al. Geneious Basic: An integrated and extendable desktop software platform for the organization and analysis of sequence data. Bioinformatics 28, 1647–1649 (2012).

92. Minh, B. Q., et al. IQ-TREE 2: New Models and Efficient Methods for Phylogenetic Inference in the Genomic Era. Mol. Biol. Evol. 37, 1530–1534 (2020).

93. Sagulenko, P., Puller, V. & Neher, R. A. TreeTime: Maximum-likelihood phylodynamic analysis. Virus Evol. 4, (2018).

94. Suchard, M. A., et al. Bayesian phylogenetic and phylodynamic data integration using BEAST 1.10. Virus Evol. 4, (2018).

95. Shapiro, B., Rambaut, A. & Drummond, A. J. Choosing Appropriate Substitution Models for the Phylogenetic Analysis of Protein-Coding Sequences. Mol. Biol. Evol. 23, 7–9 (2006).

96. Drummond, A. J., Ho, S. Y. W., Phillips, M. J. & Rambaut, A. Relaxed Phylogenetics and Dating with Confidence. PLoS Biol. 4, e88 (2006).

97. Kass, R. E. & Raftery, A. E. Bayes Factors. J. Am. Stat. Assoc. 90, 773–795 (1995).

98. Bielejec, F., et al. SpreaD3: Interactive Visualization of Spatiotemporal History and Trait Evolutionary Processes. Mol. Biol. Evol. 33, 2167–2169 (2016).

99. Bedford, T., et al. Global circulation patterns of seasonal influenza viruses vary with antigenic drift. Nature 523, 217–220 (2015).

