## Supplemental material for "Intensive transmission in wild, migratory birds drove rapid geographic dissemination and repeated spillovers of H5N1 into agriculture in North America"

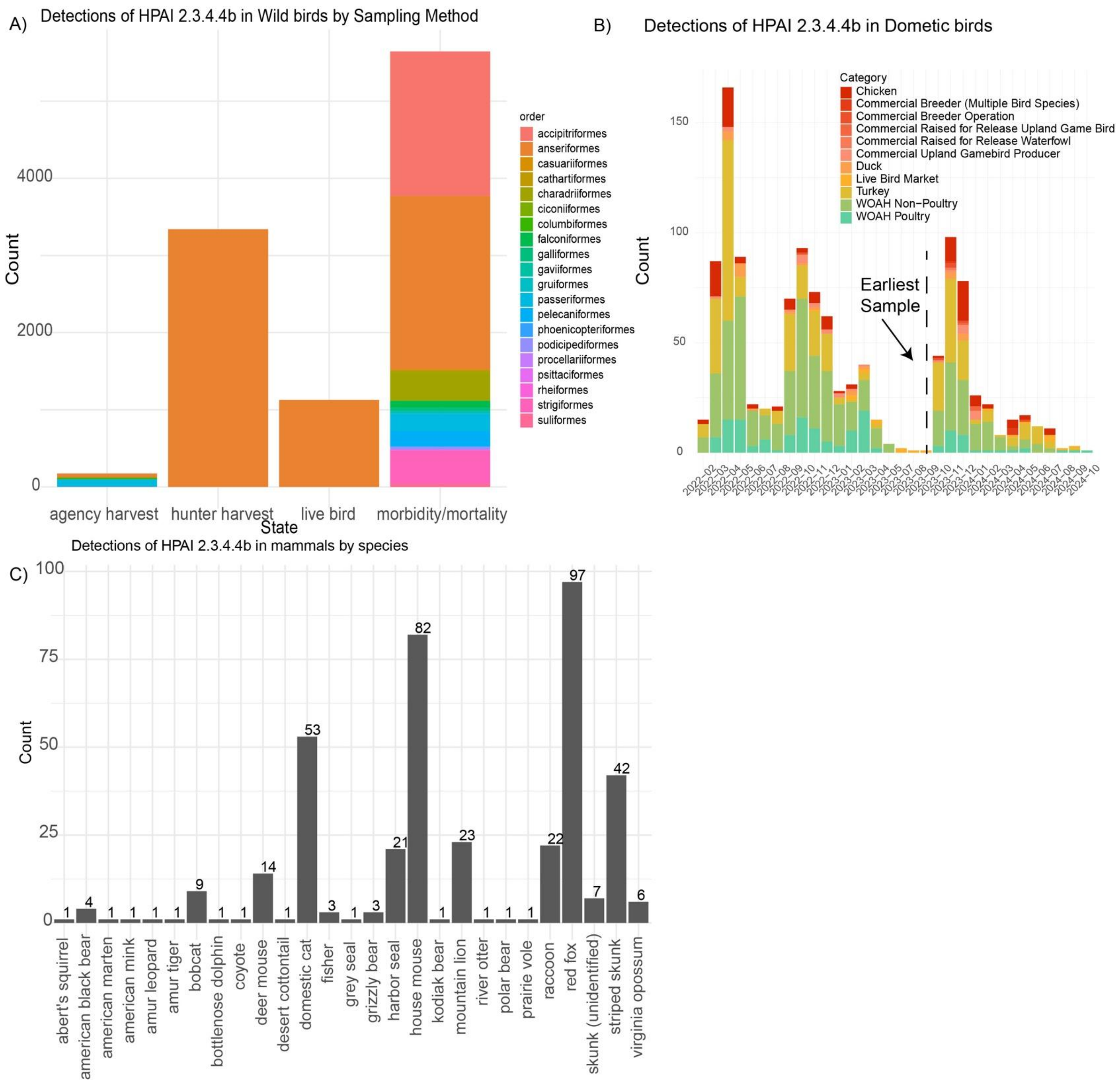


**Figure S1.** A) Number of detections of HPAI in North American wild birds by collection method. Morbidity and mortality refer to sick or dead birds. B) Domestic bird detections of HPAI in North America on a monthly basis colored by production type. C) Number of detections in mammals by species.

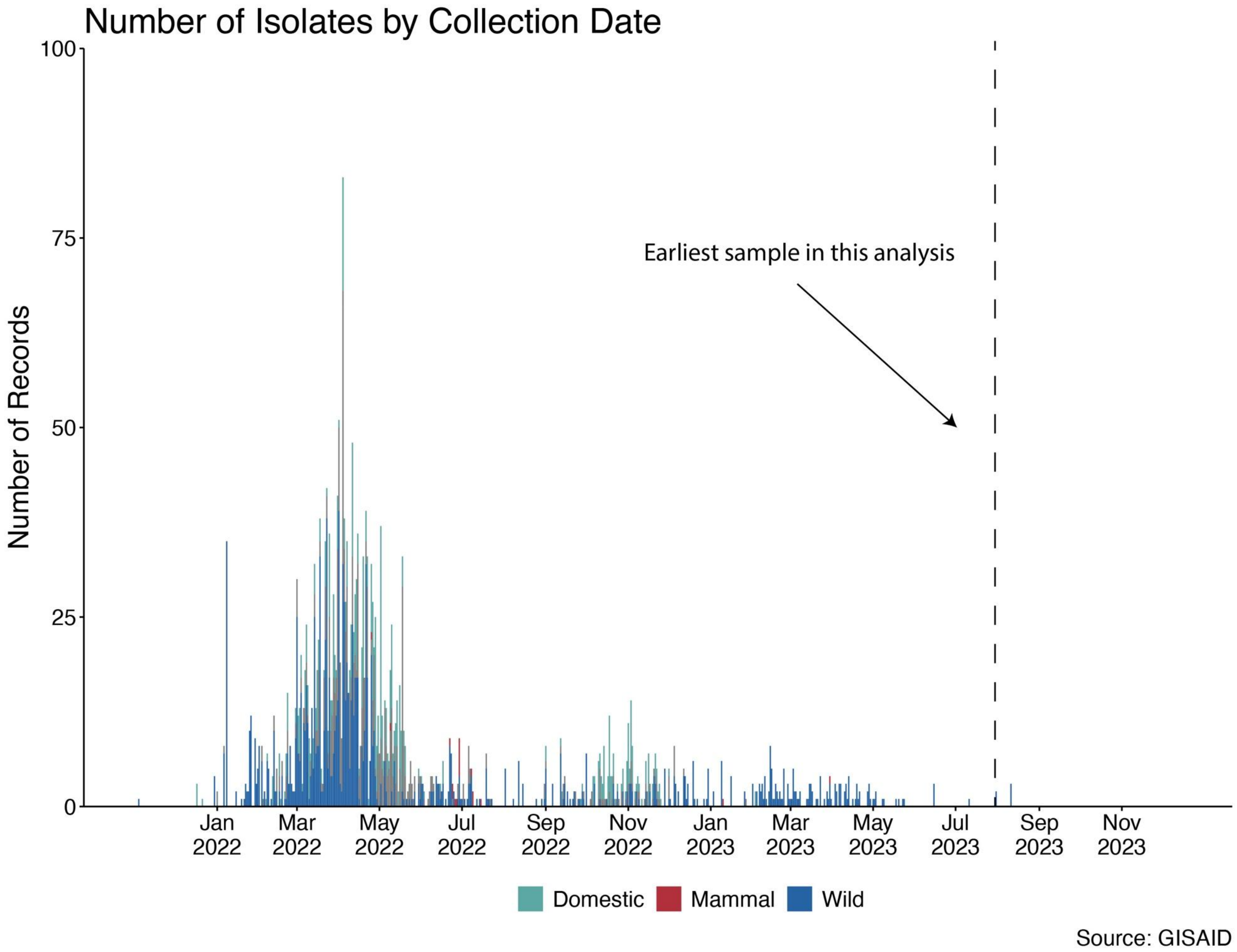


**Figure S2**. Number of HA isolates for HPAI in North America by collection date available in GISAID.

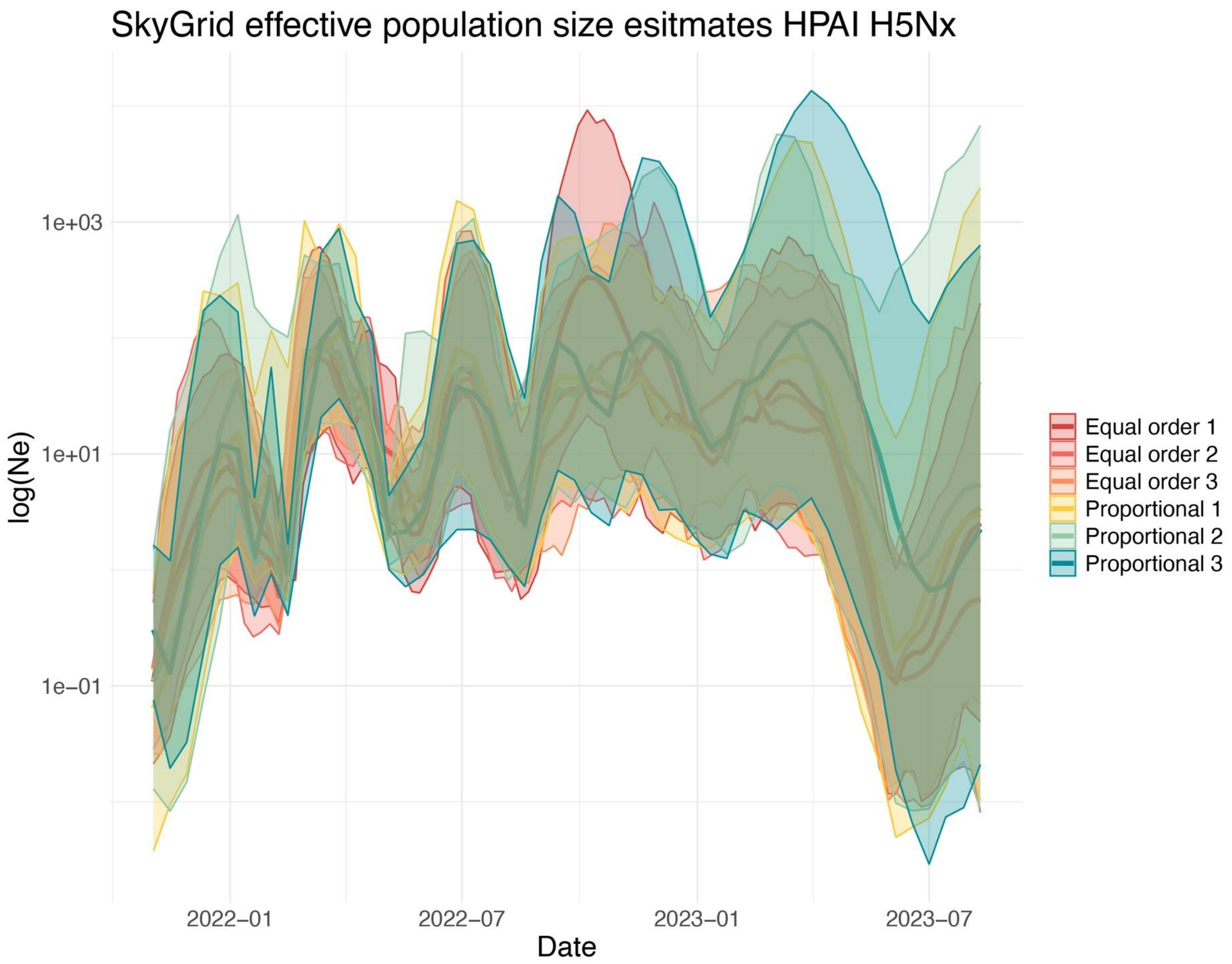


**Figure S3**. A) SkyGrid Coalescent effective population size reconstructions for six different subsamples of HPAI in North America between 2021 and 2023. Three subsamples with equal proportions of host orders and three subsamples with number of sequences proportional to detections in those hosts.

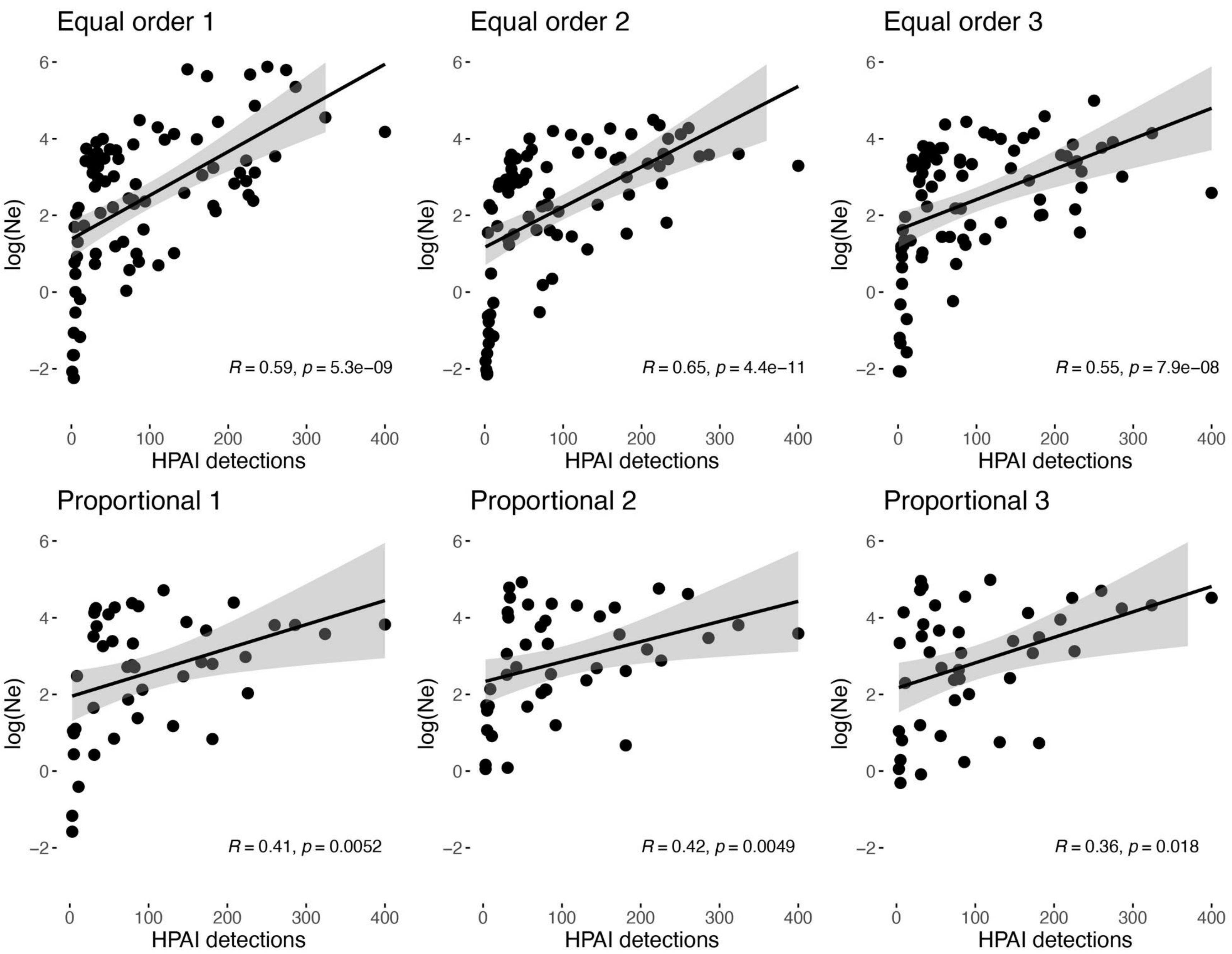


**Figure S4.** Spearman correlation plot of effective population size estimates vs detections at corresponding timepoints.

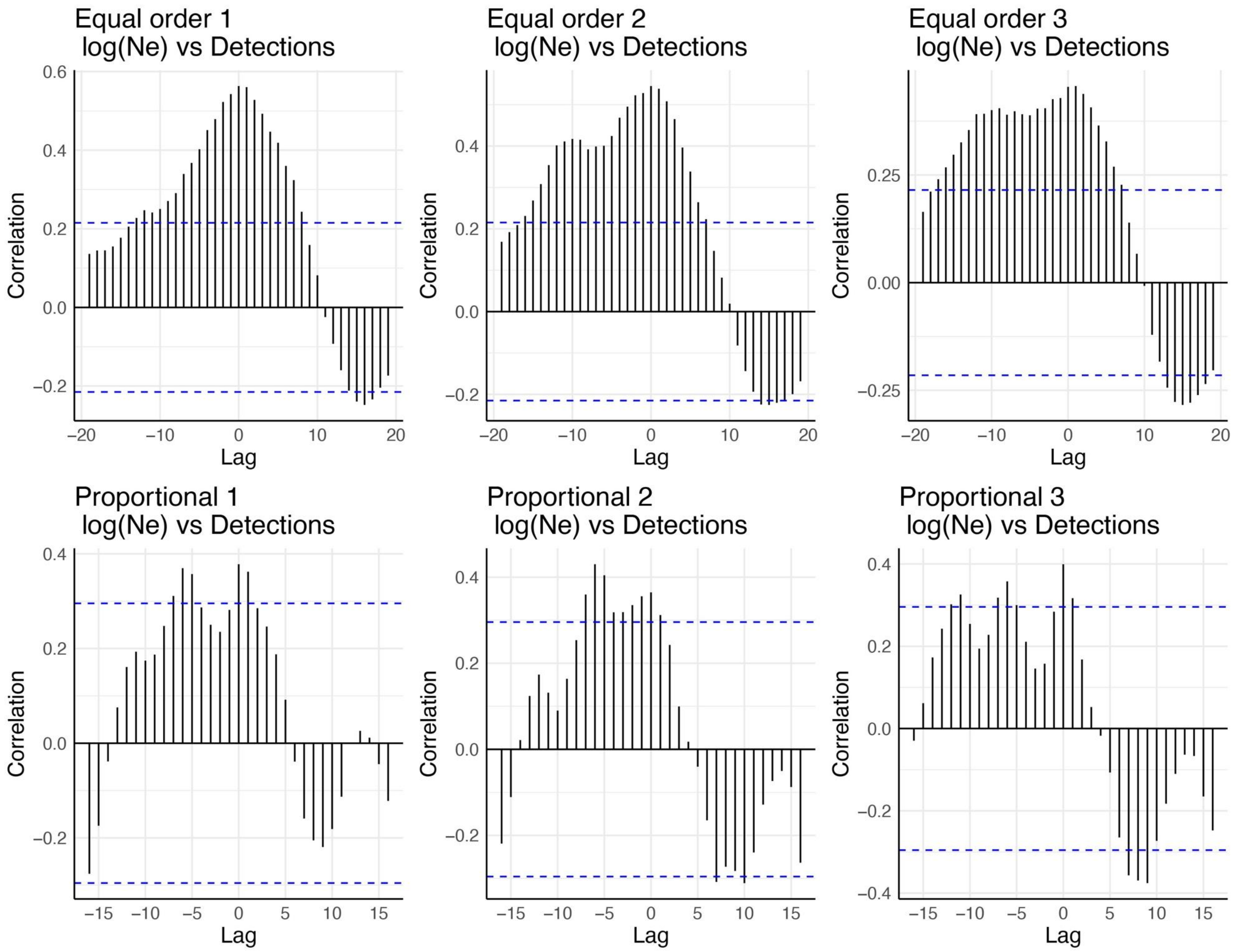


**Figure S5.** Cross correlation plot of effective population size estimates and detections of HPAI. The x-axis represents the time lag in weeks and the y-axis represents the correlation. Dotted lines represent the significance thresholds.

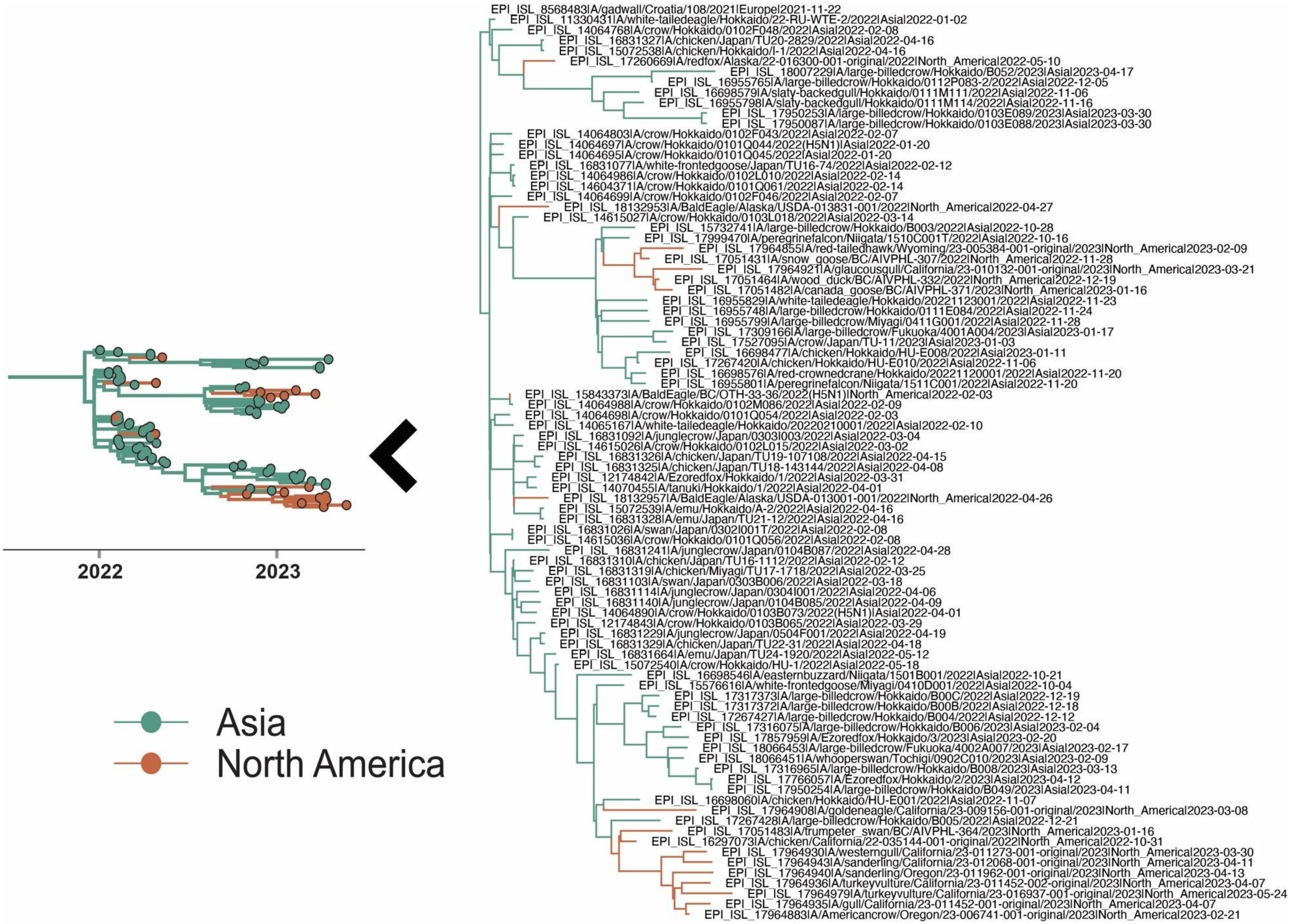


**Figure S6.** Clade associated with Asian introductions of HPAI into North America with taxa labeled.

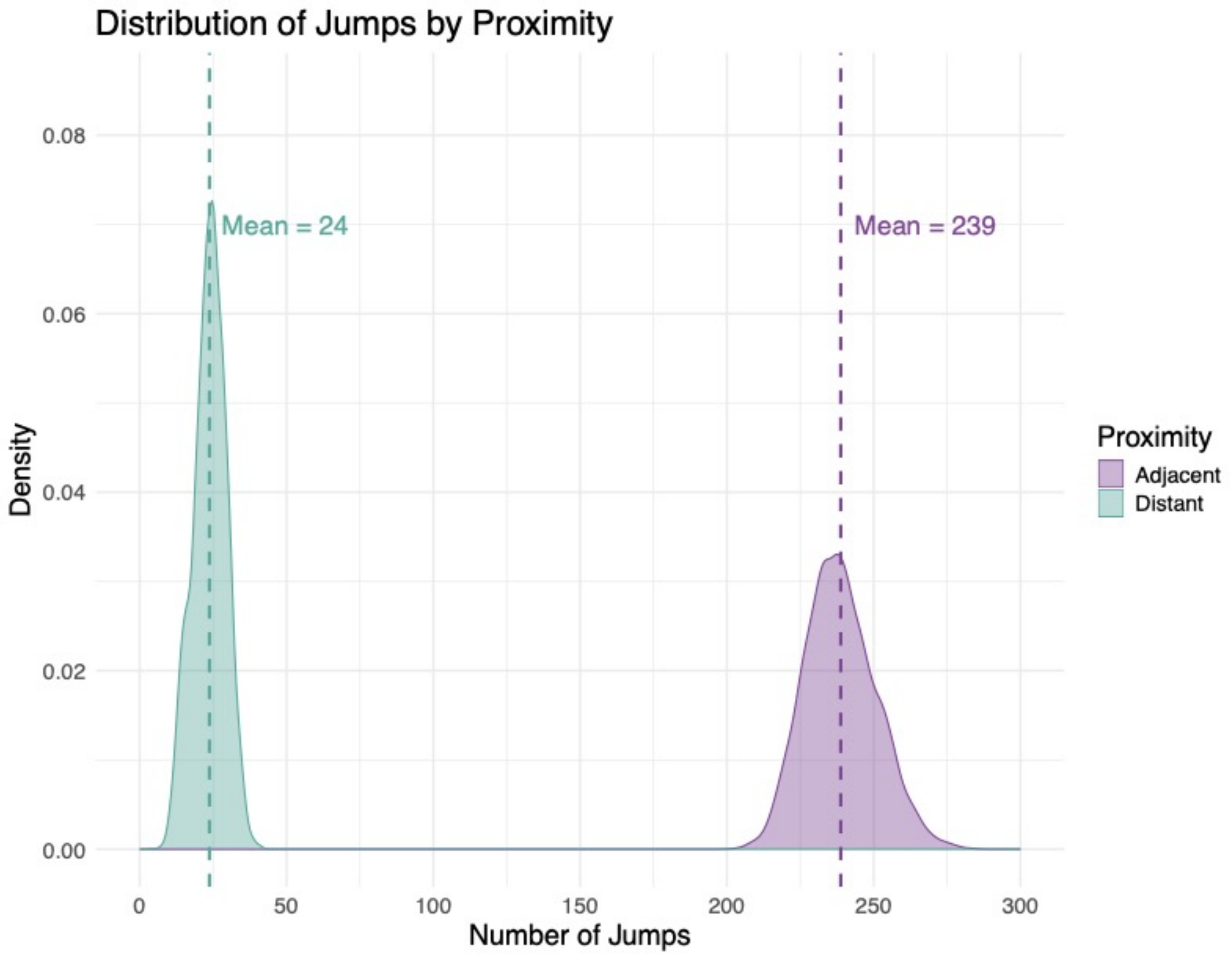


**Figure S7.** Mean number of Markov jumps across the posterior distribution of trees between adjacent (directly next to each other) and distant flyways.

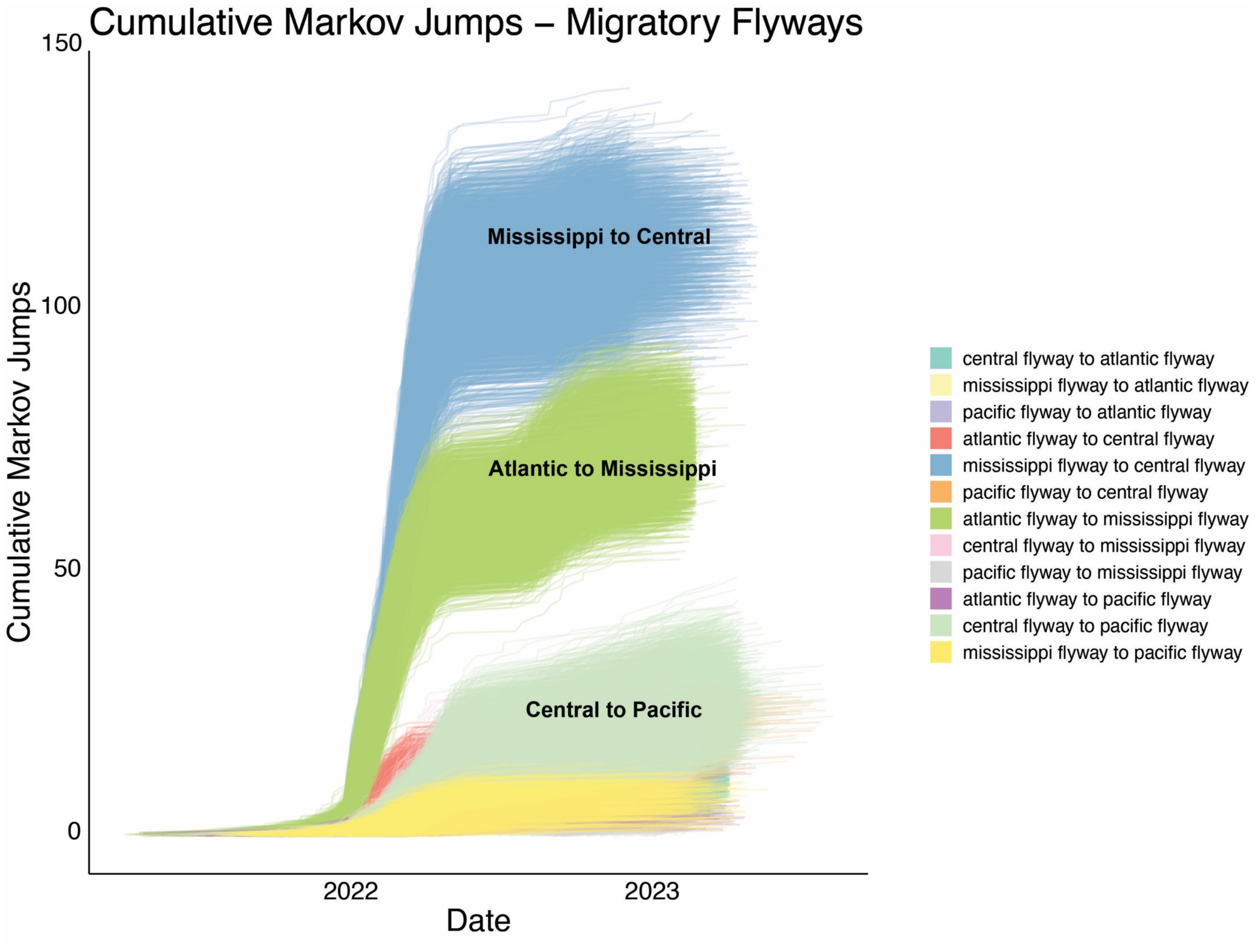


**Figure S8.** Cumulative Markov jumps over time between flyways.

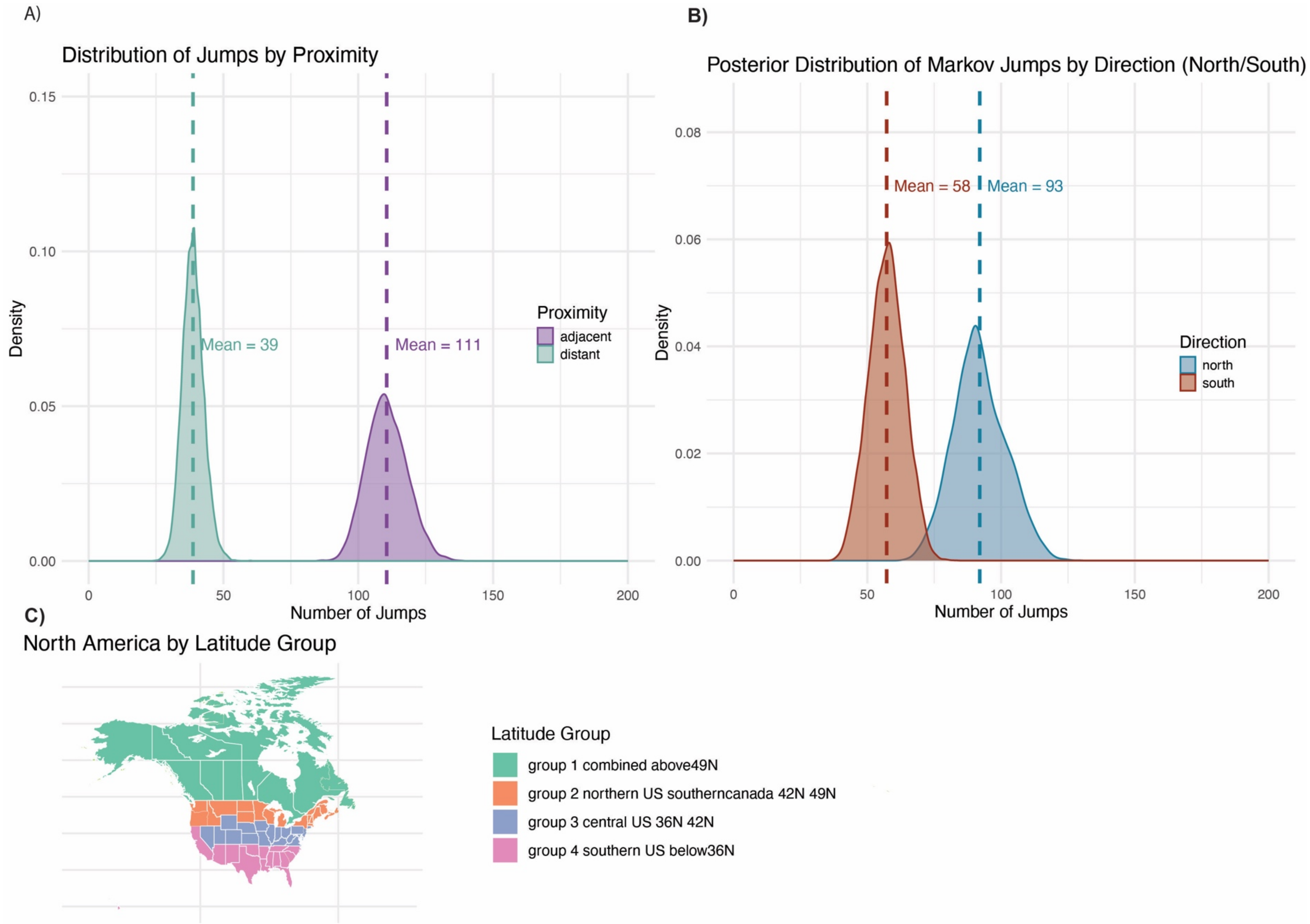


**Figure S9.** A) Mean number of Markov jumps across the posterior distribution of trees between adjacent (directly next to each other) and distant geographic groups based on latitude. B) Mean number of Markov jumps across the posterior distribution of trees based on direction of jump (North or South between geographic groups based on latitude). C) Map of North America with states colored by latitude groups.

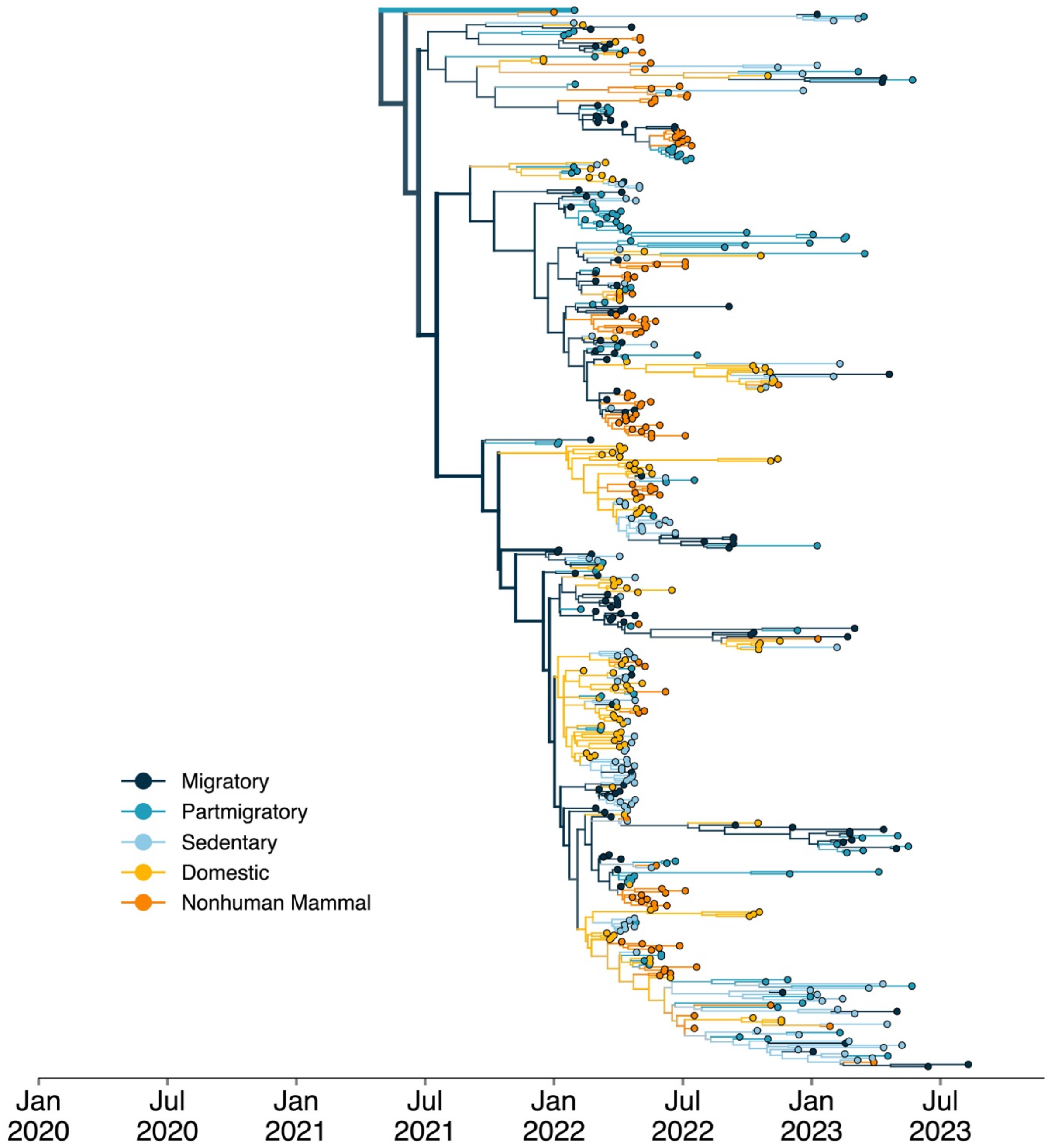


**Figure S10.** Phylogenetic reconstruction of n=1000 sequences colored by Migratory behavior of host (Determined using AVONET). Color corresponds to the migratory behavior of the host inferred for the branches and for the host of the tip.

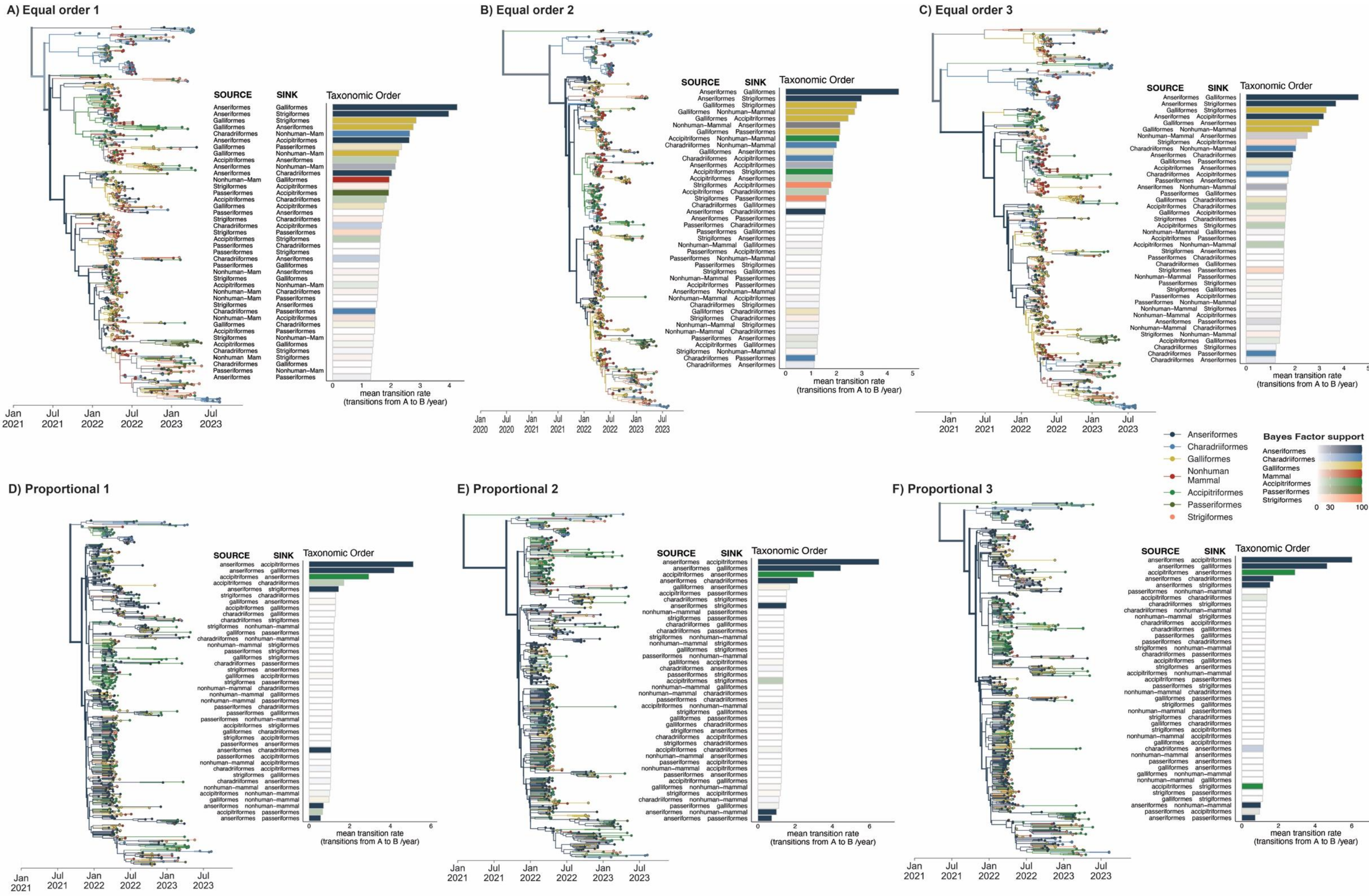


**Figure S11:** MCC trees for each subsampled dataset for analysis of host order transmission where colors correspond to host Order. with corresponding bar plot showing discrete trait rates. A-C) are equal number of order datasets and D-F) are case proportional datasets.

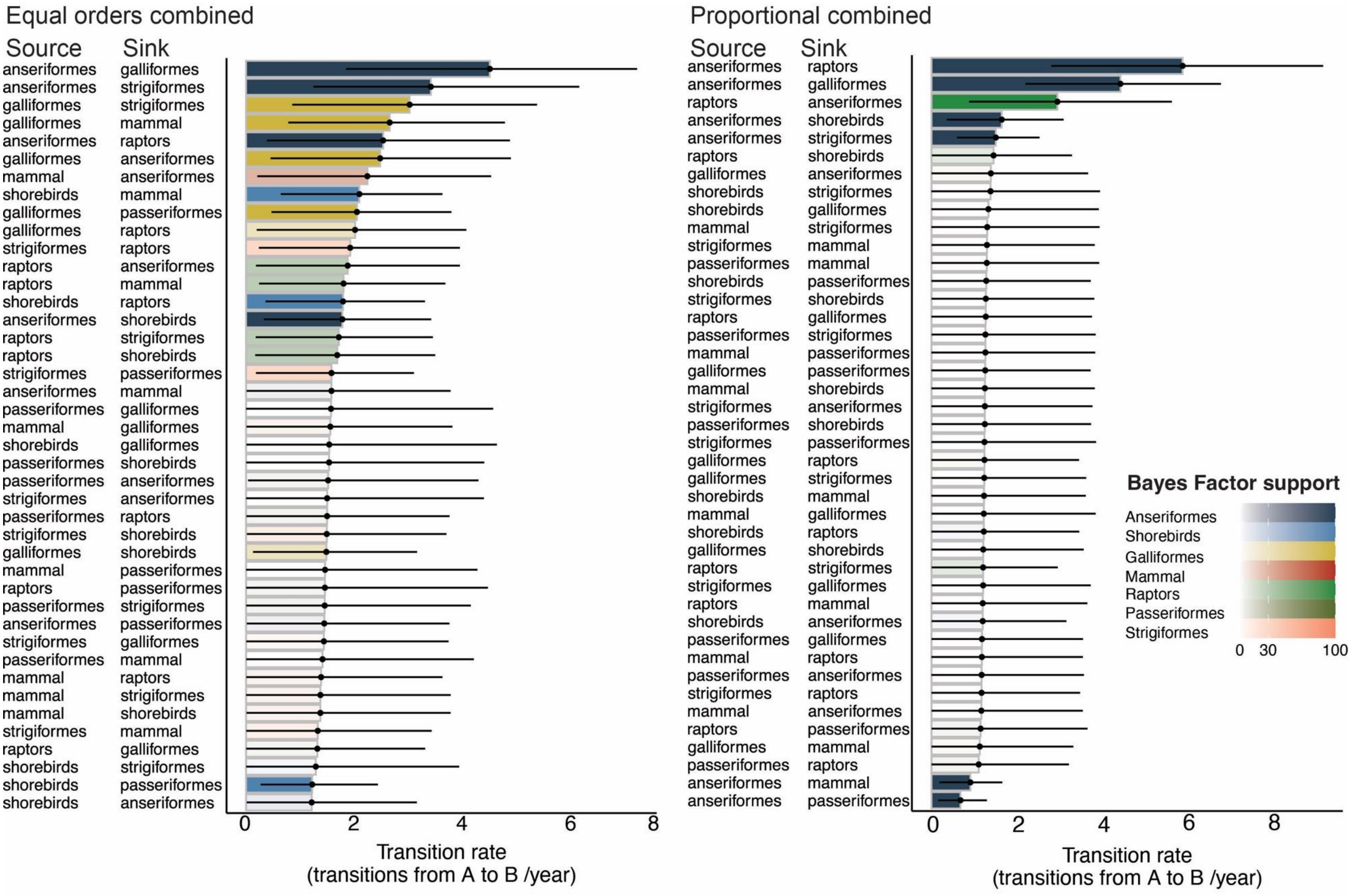


**Figure S12.** Transition rates (dot represents mean transition and bars represent 95% HPD) from combined results of the BSSVS of combined results for equal orders schema (Left) and proportional scheme (left) where color of the bar corresponds to the source population and the opacity corresponds to the bayes factor support (where white corresponds to BF < 3 and full color corresponds to BF > 100).

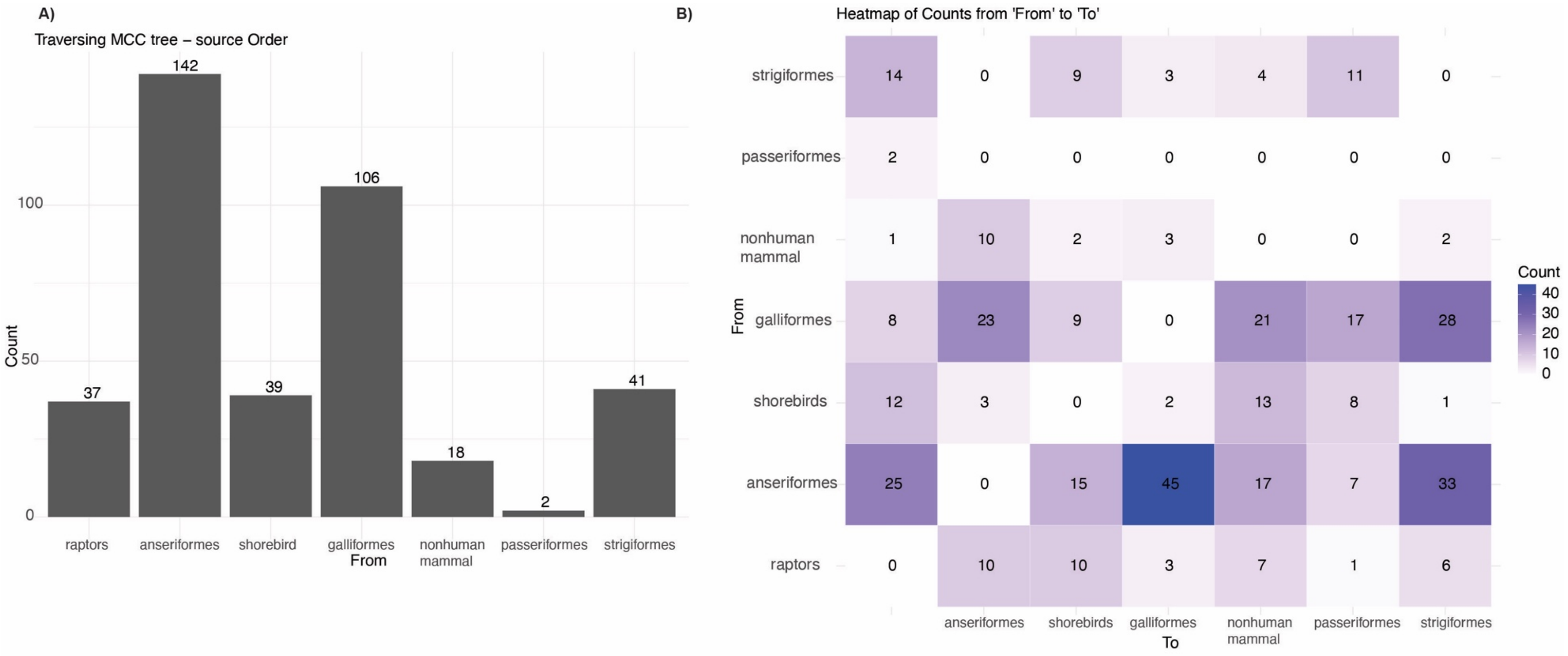


**Figure S13**. Results from traversing MCC tree and enumerating number of transitions between host orders. A) Number of times a given host order transition to another host order. B) Number of transitions between each host order pair. Transitions calculated using the BALTIC python package.

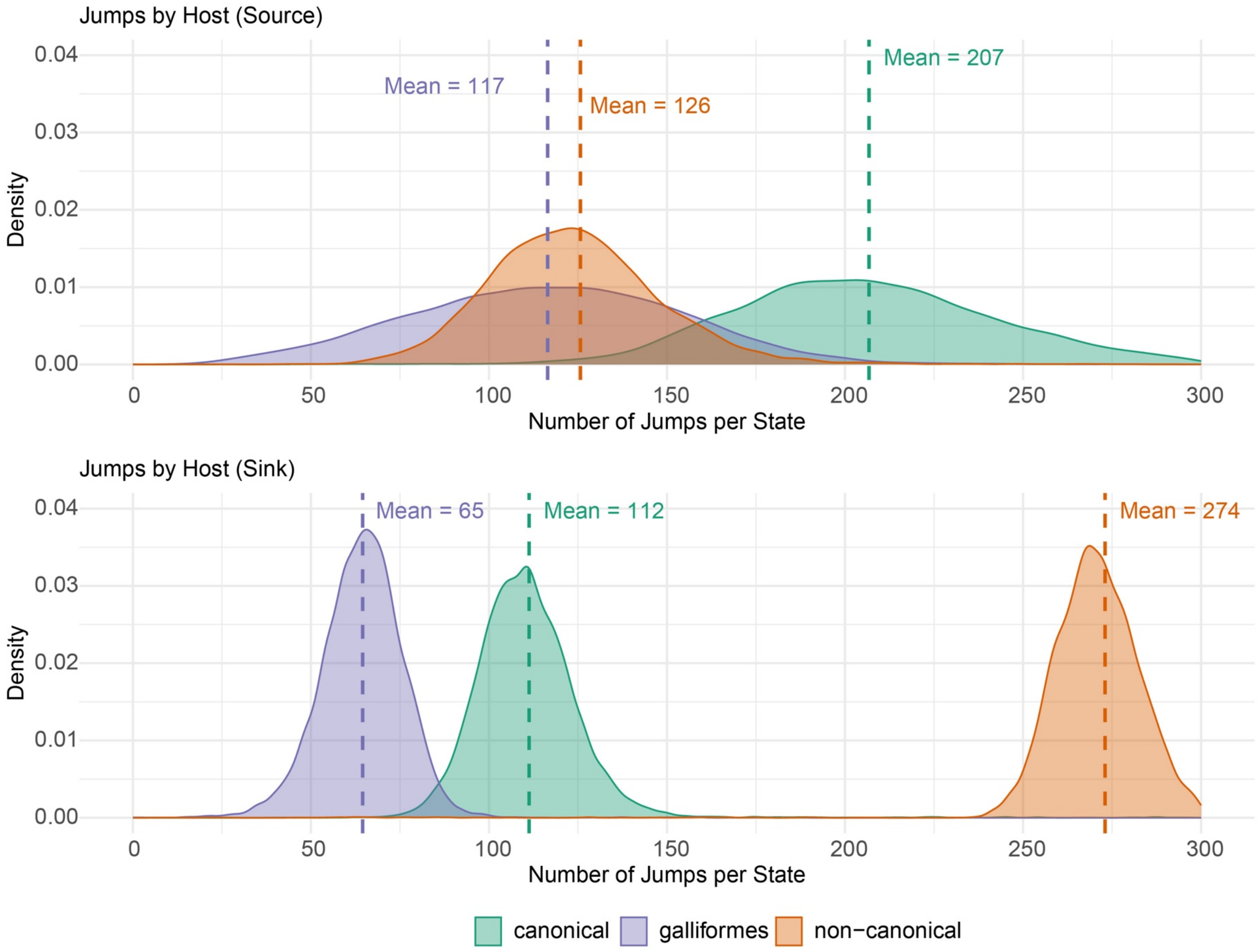


**Figure S14.** Top panel shows density plots for mean number of Markov jumps across the posterior distribution of trees for jumps where the host acts as source. Bottom panel shows density plots for mean number of Markov jumps across the posterior distribution of trees for jumps where the host acts as sink.

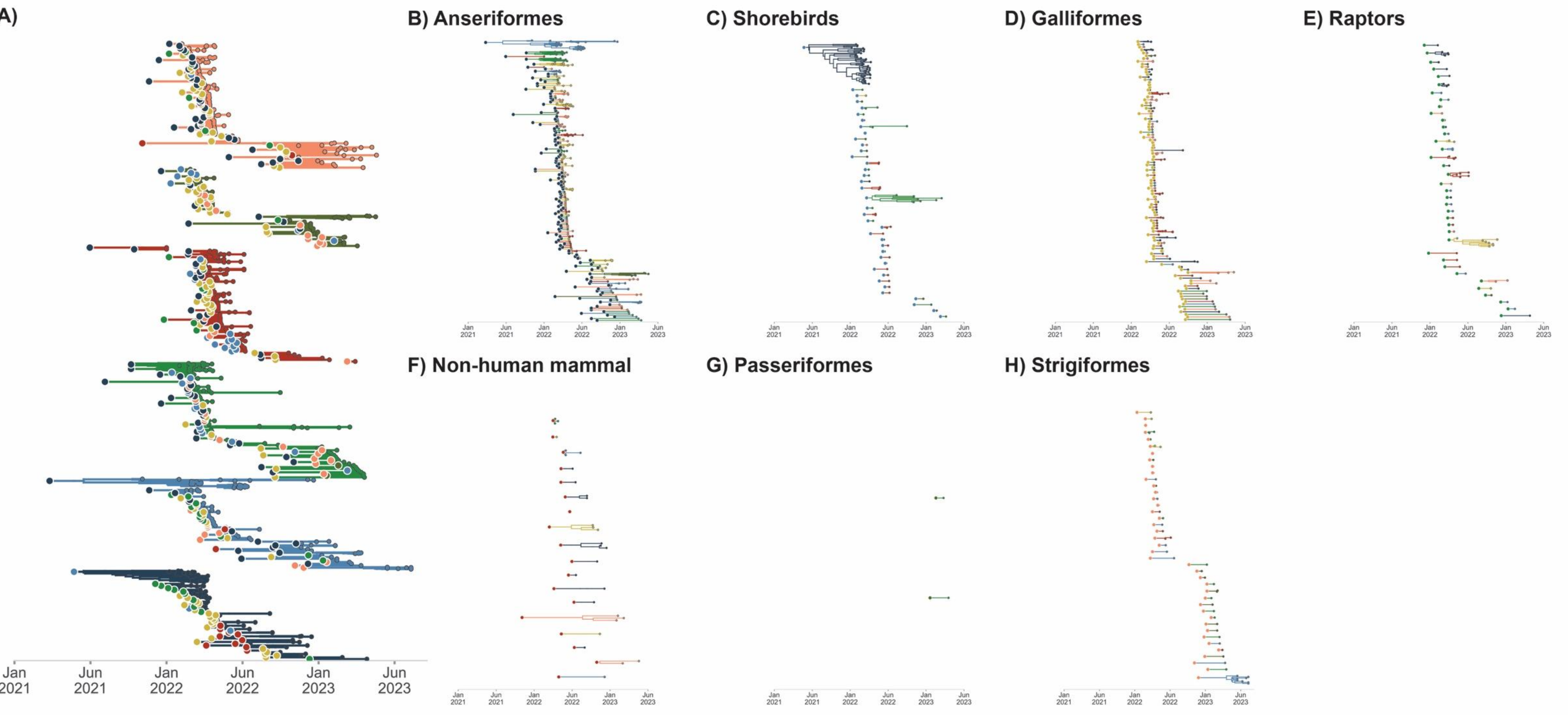


**Figure S15**: A) Exploded tree view of the MCC tree of equal order phylogenetic reconstruction where subtrees represent the traversal of a tree from the root to the tip where the state is unchanged from the initial state (given by the large dot on left) to the tips represented by the smaller dots representing continuous chains of transmission within a given state. Colors correspond to the state of a given tip, branch node. B-H represent the exploded trees faceted by taxonomic host at the origin: B) Anseriformes, C) Shorebirds , D) Galliformes, E) Raptors, F) Non-human mammal, G) Passeriformes, H) Strigiformes.

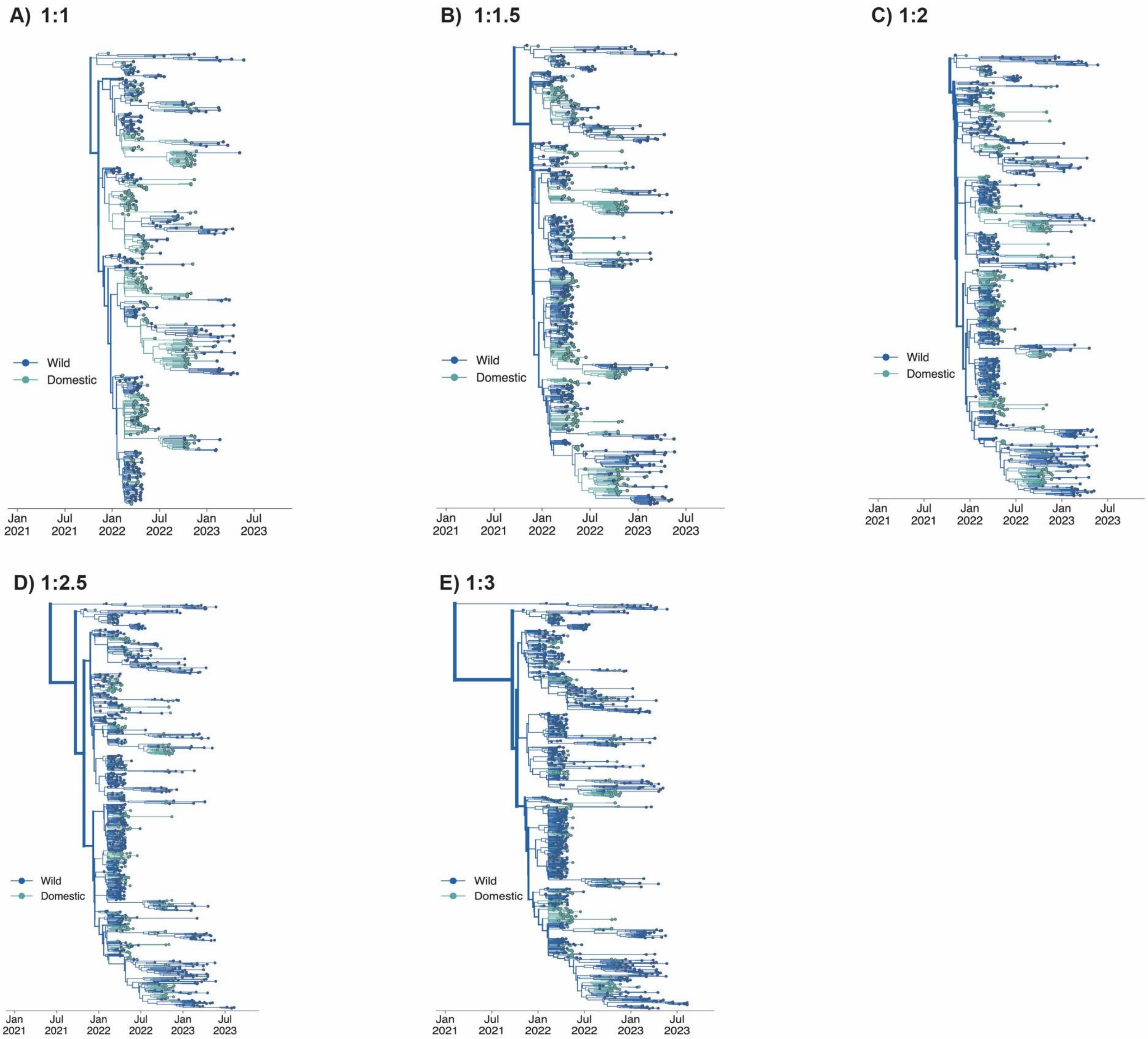


**Figure S16:** Two-state rarefaction analysis (domestic:wild MCC trees A-E) show the ratios of domestic to wild in increasing order of the number of wild bird sequences. Tips and branches are colored by the state (wild or domestic) of the sample and inferred state respectively.

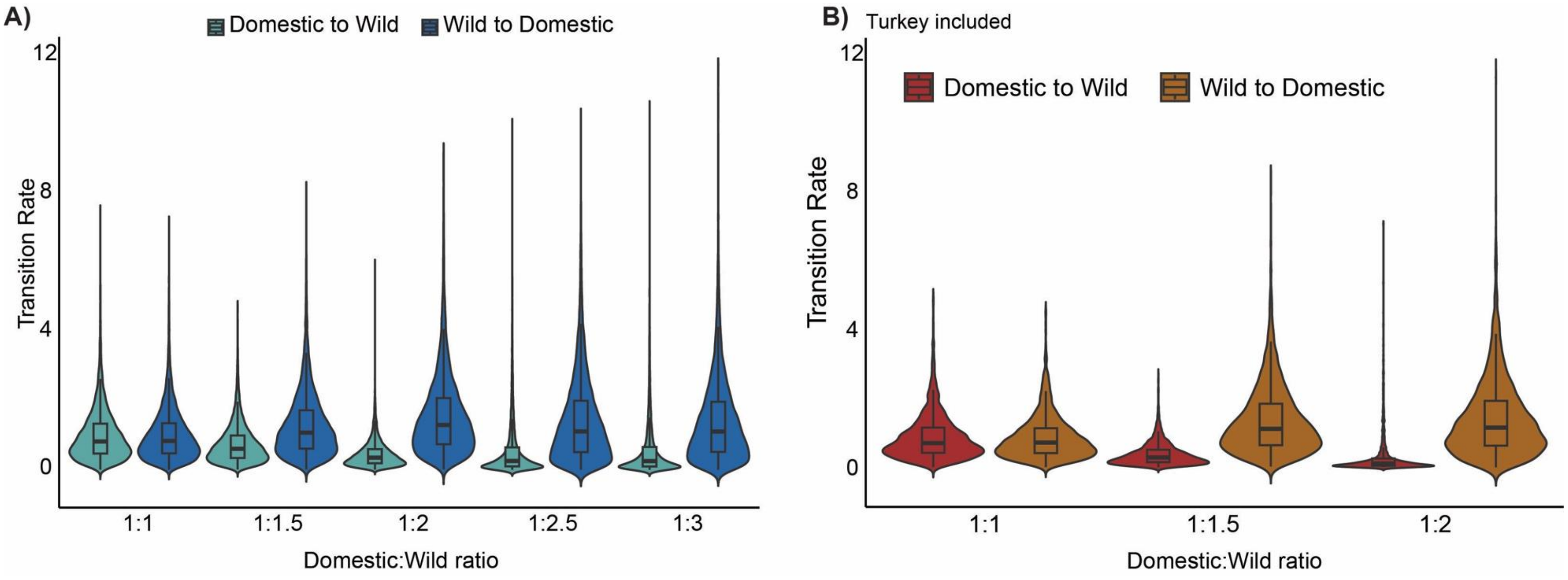


**Figure S17.** Violin plots of discrete trait transition rates from Domestic to Wild and Wild to Domestic for A) two state rarefaction and B) two state rarefaction including turkey sequences. All transition rates had a posterior probability of 0.99.

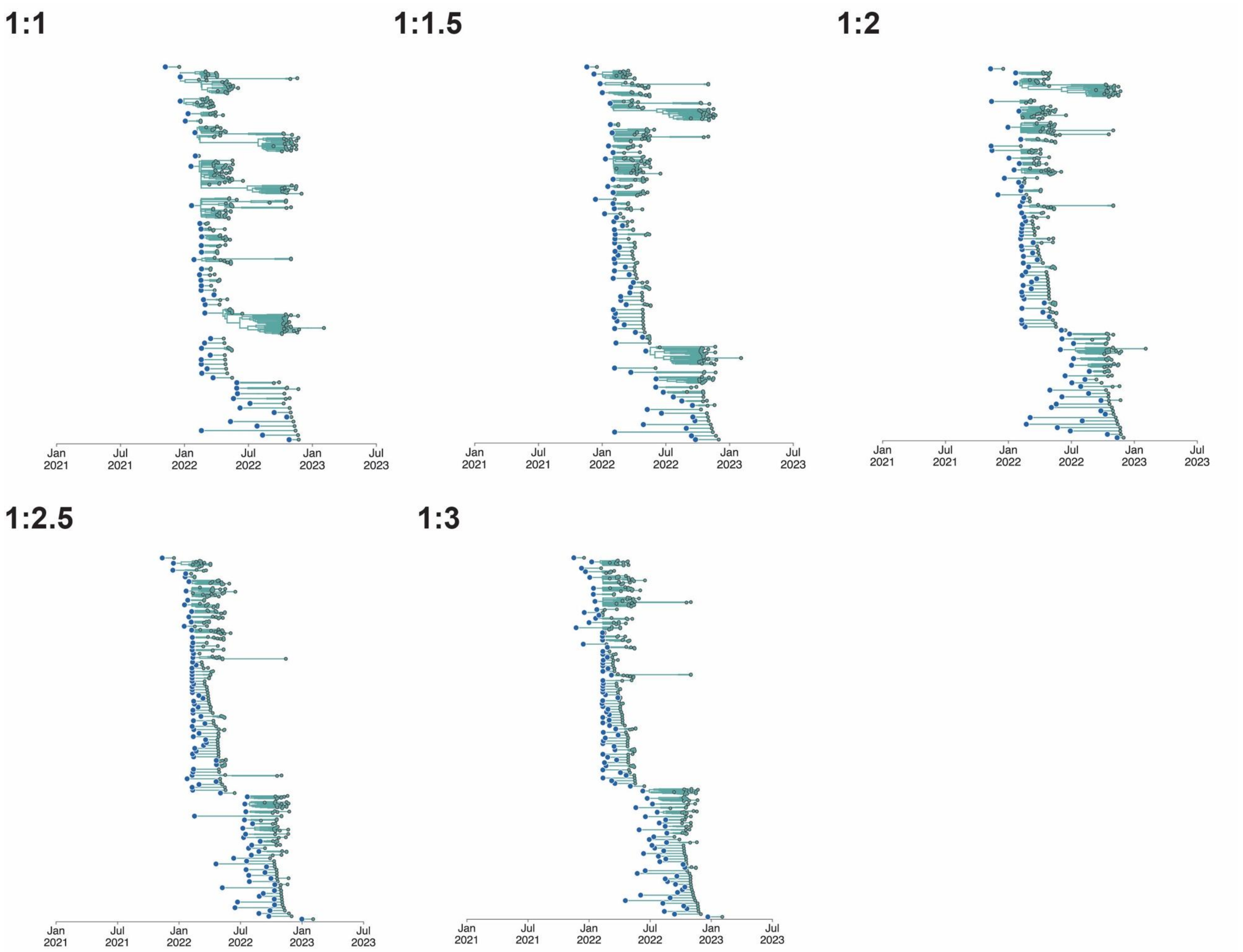


**Figure S18**: A) Exploded tree view of the MCC tree of two state rarefaction for wild to domestic transitions. Colors correspond to the state of a given tip, branch node.

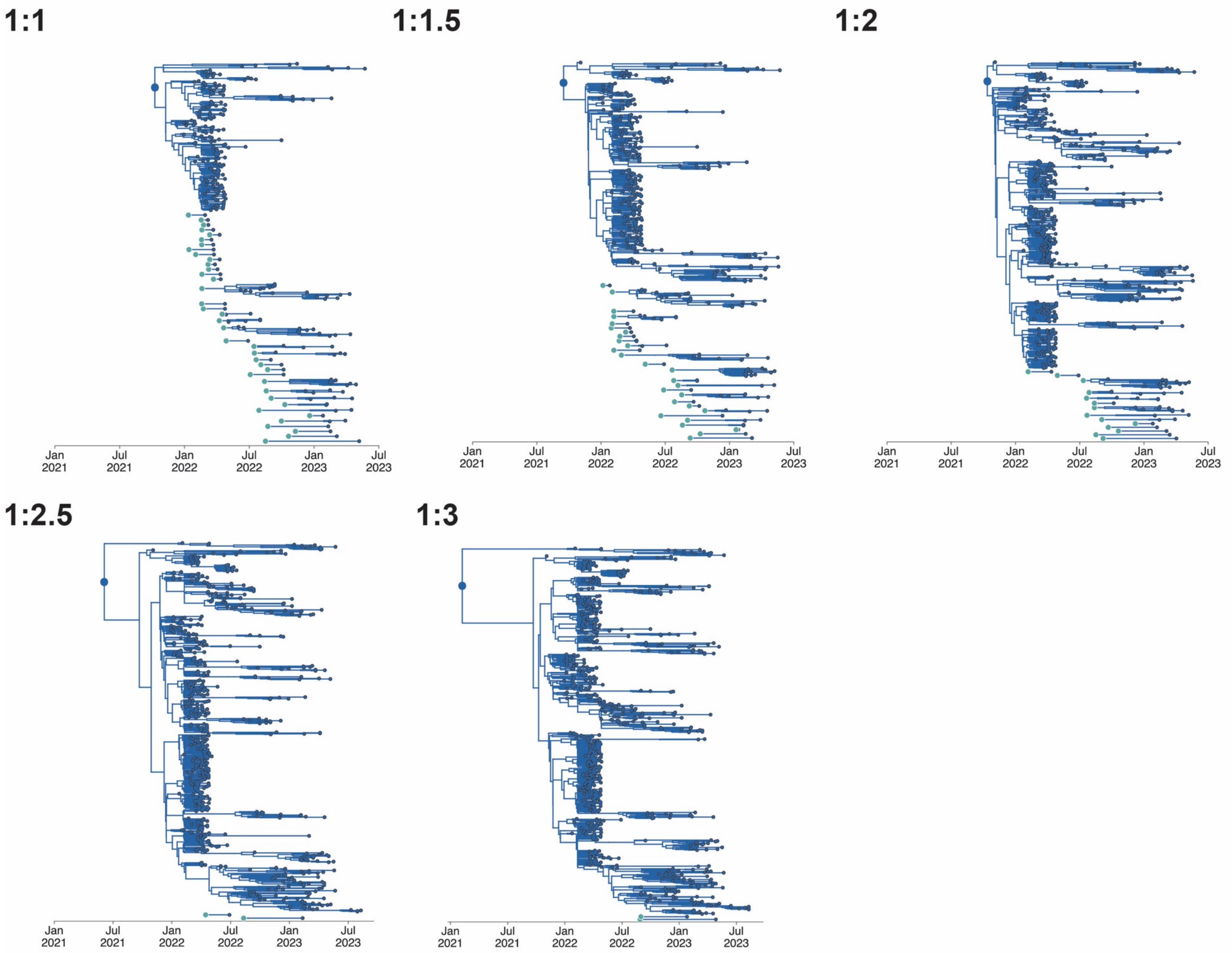


**Figure S19**: A) Exploded tree view of the MCC tree of two state rarefactions for domestic to wild transitions. Colors correspond to the state of a given tip, branch node.

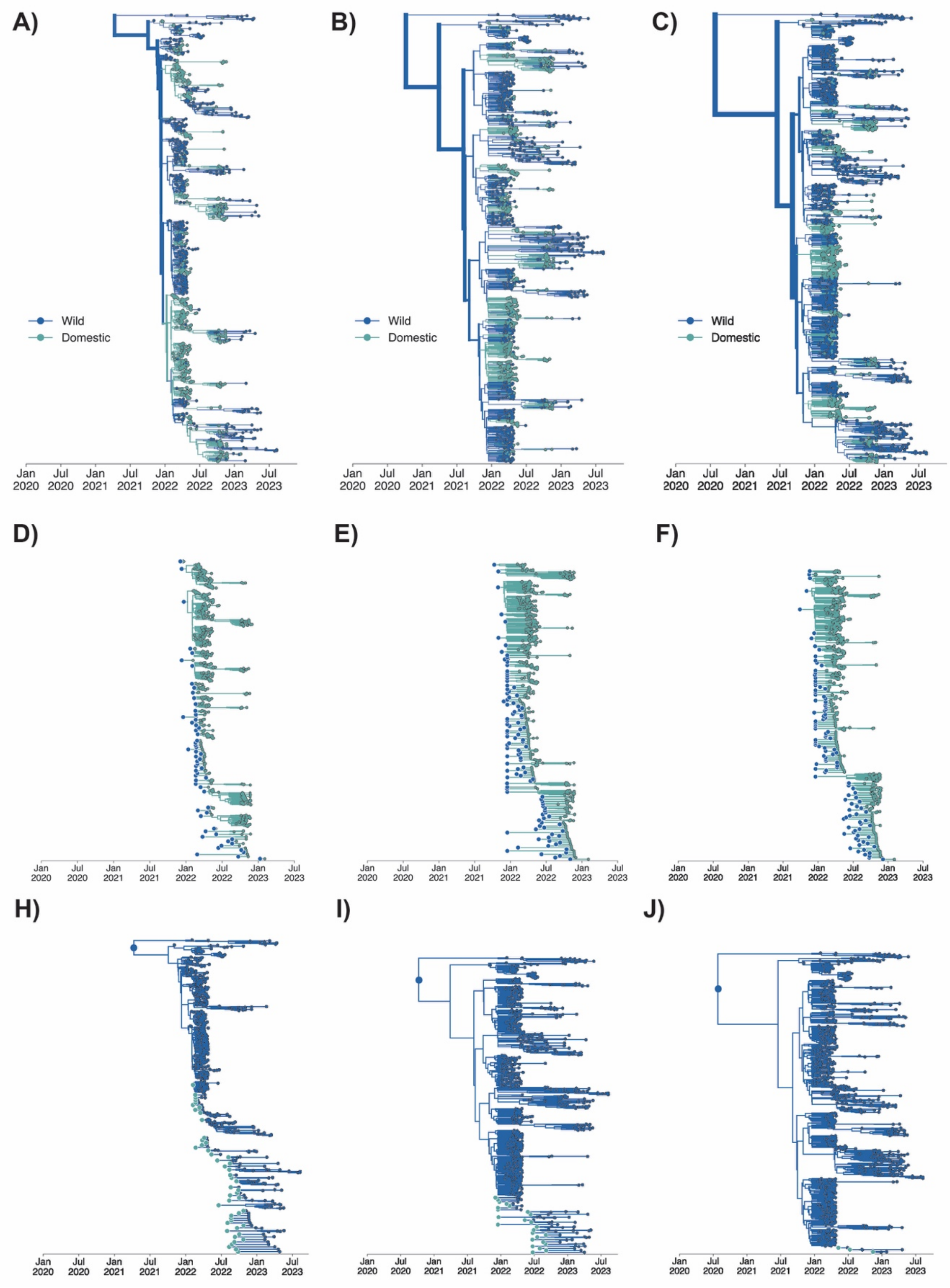


**Figure S20**: A-C) MCC trees of the ratios of domestic to wild including turkey sequences in increasing order of the number of wild bird sequences. Tips and branches are colored by the state (wild or domestic) of the sample and inferred state respectively. D-F) Exploded tree views of the MCC trees for each titration of two state rarefactions including turkey for wild to domestic transitions. G-J) Exploded tree views of the MCC trees for each titration of two state rarefactions including turkey for domestic to wild transitions. Colors correspond to the state of a given tip, branch node.

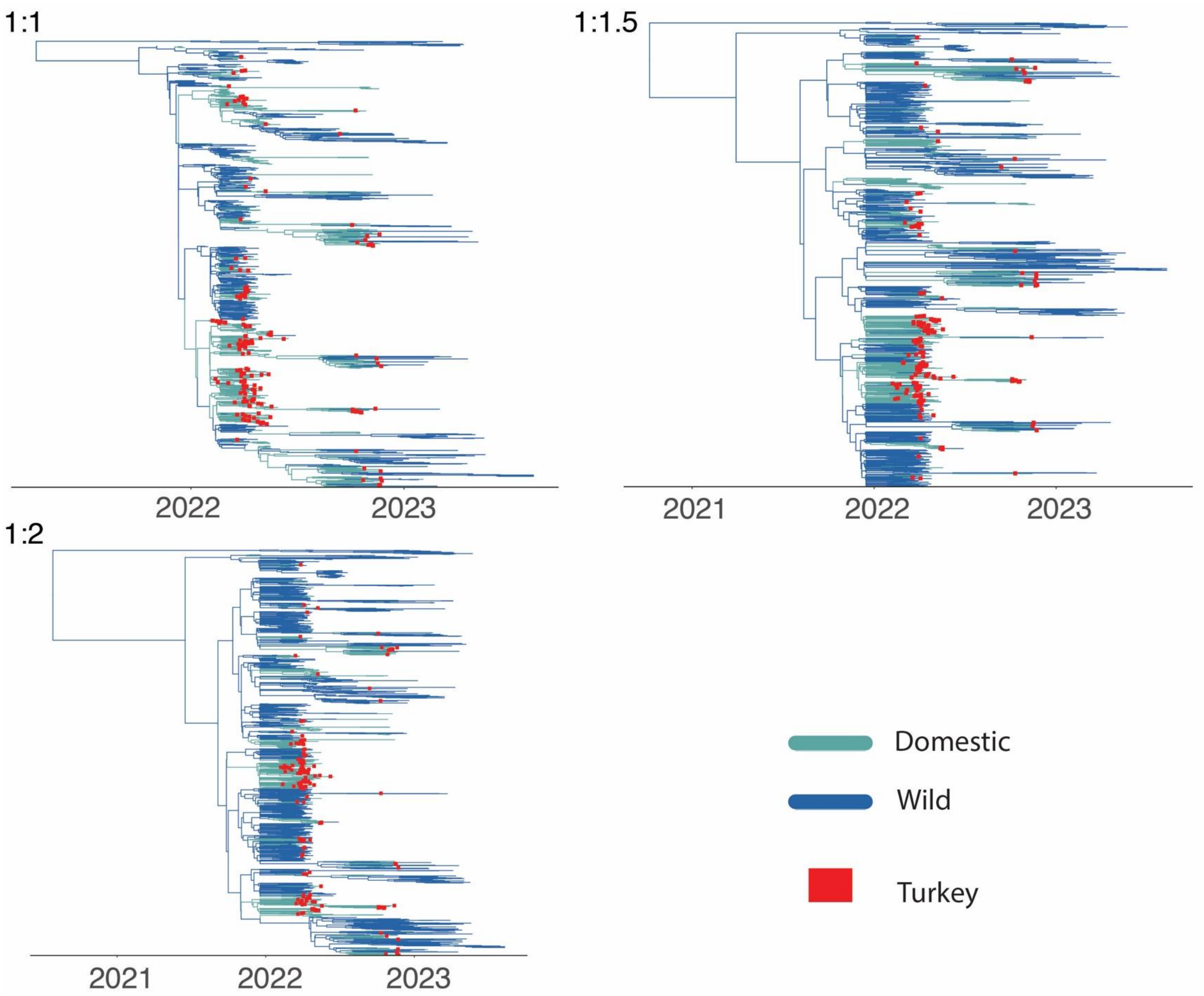


**Figure S21**: MCC trees of the ratios of domestic to wild including turkey sequences in increasing order of the number of wild bird sequences. Tips and branches are colored by the state (wild or domestic) of the sample and inferred state respectively with turkey sequences annotated with red boxes.

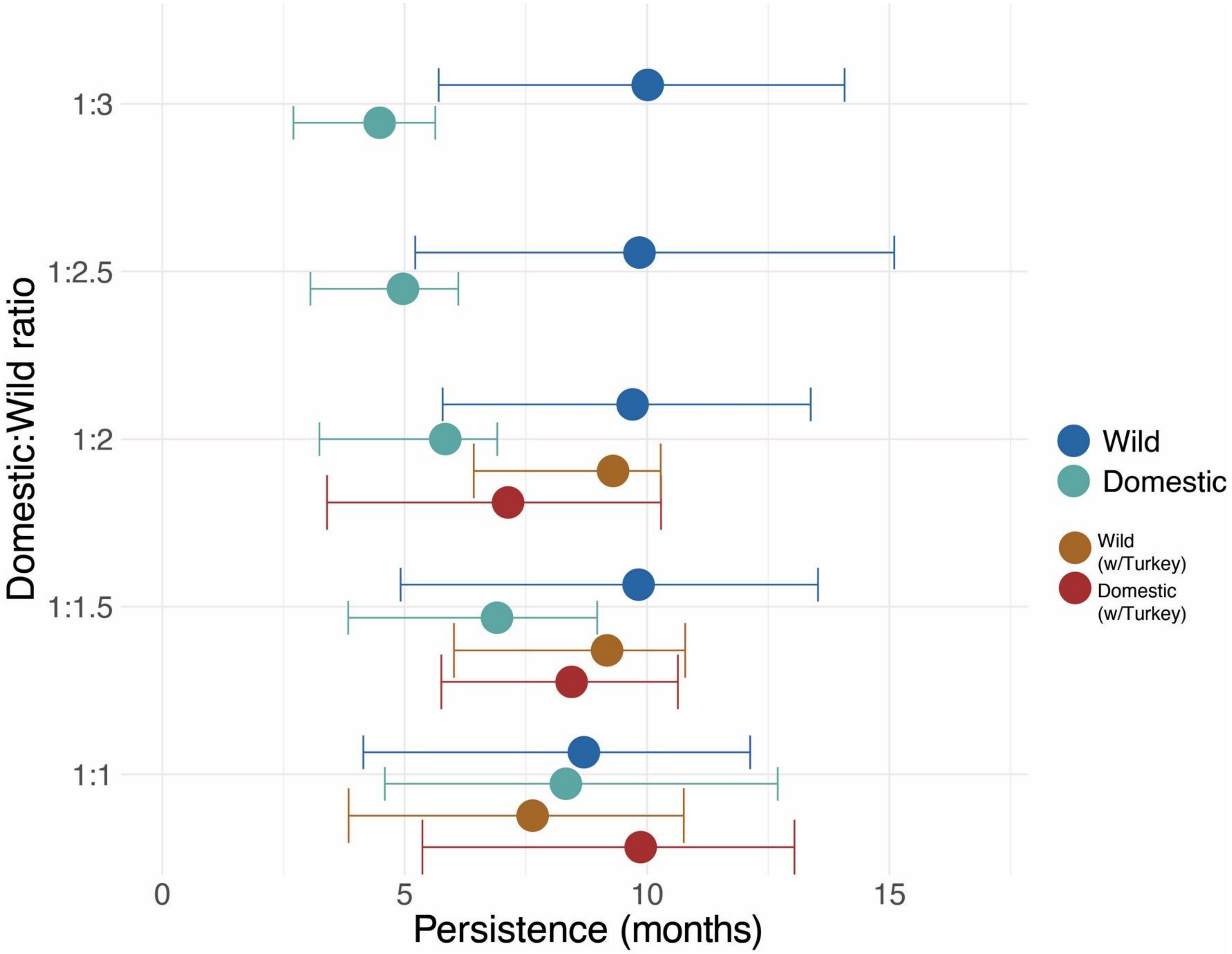


**Figure S22.** Combined results of PACT analysis for each titration in the 2-state rarefaction and 2-state rarefaction including turkey. Colors correspond to analysis.

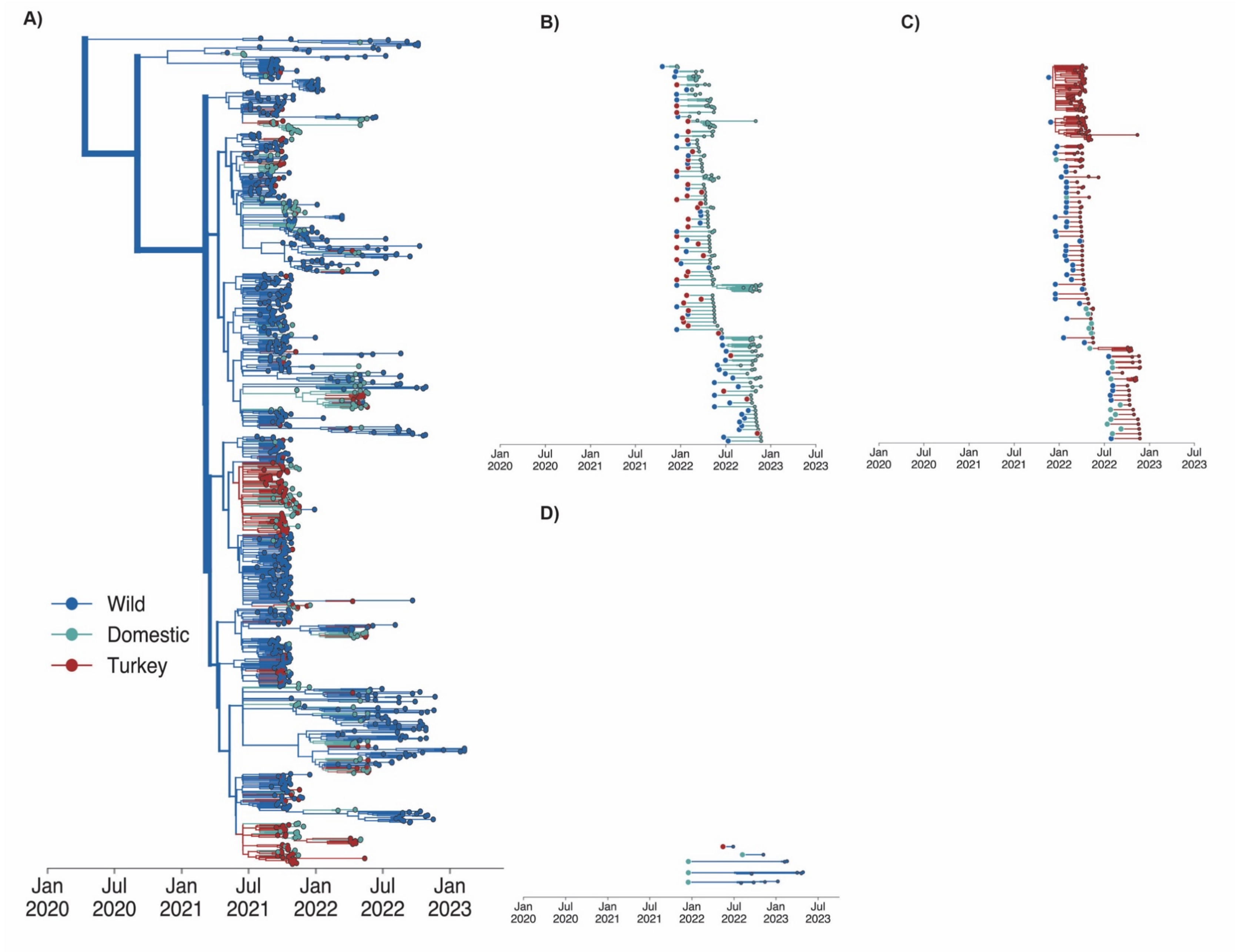


**Figure S23.** A) MCC tree of three state analysis of 1:2 (domestic:wild) sequence dataset with equal numbers of turkey and domestic bird sequences. B) Exploded tree view of the three state 1:2 MCC tree for transitions into domestic. C) Exploded tree view of the three state 1:2 MCC tree for transitions into turkey. D) Exploded tree view of the three state 1:2 MCC tree for transitions into wild.

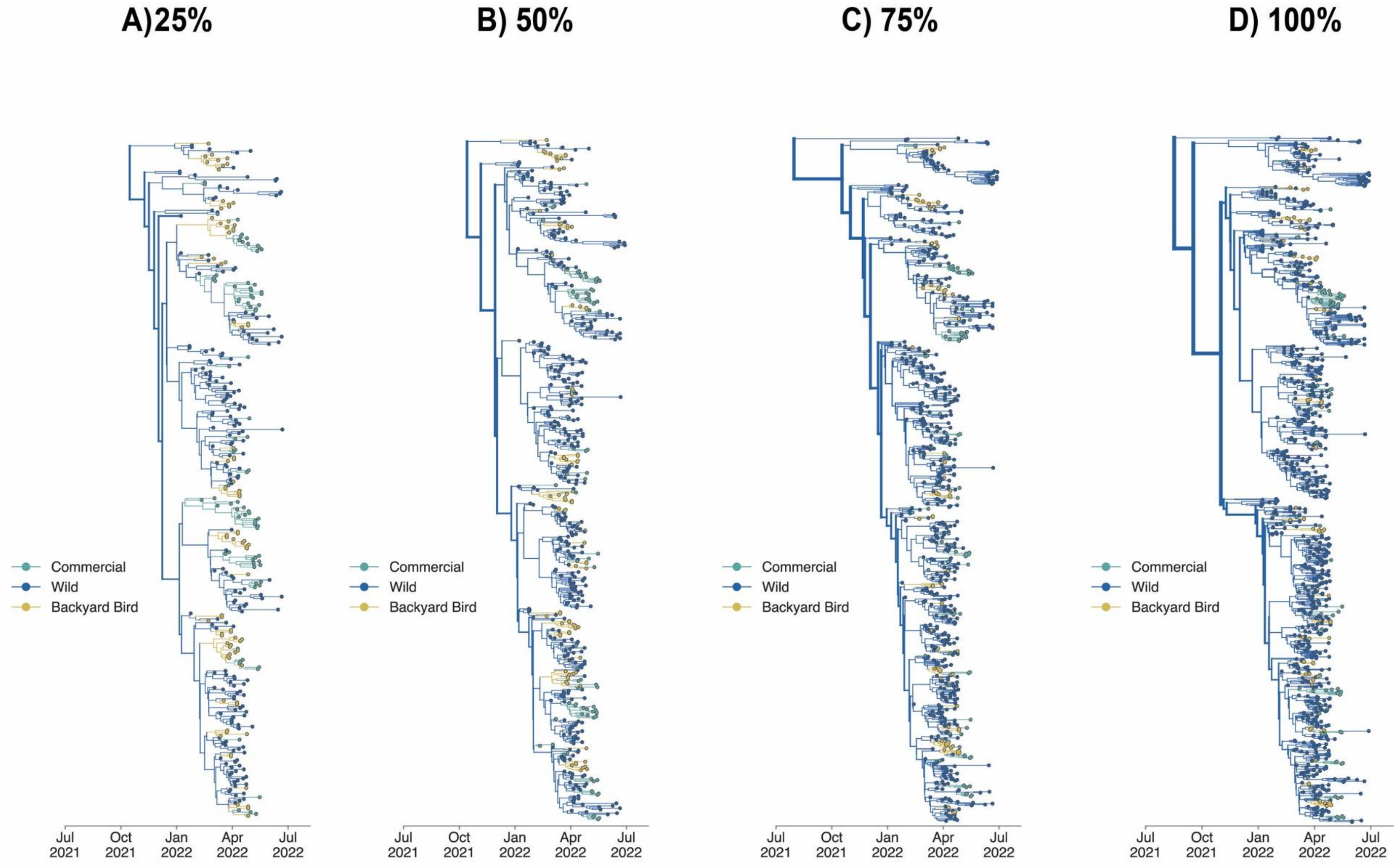


**Figure S24**: MCC trees for each tree in the three-state titration analysis. Percentage refers to the percentage of all available wild sequences used in the given analysis. Colors correspond to the host domesticity status (wild, backyard bird, commercial bird).

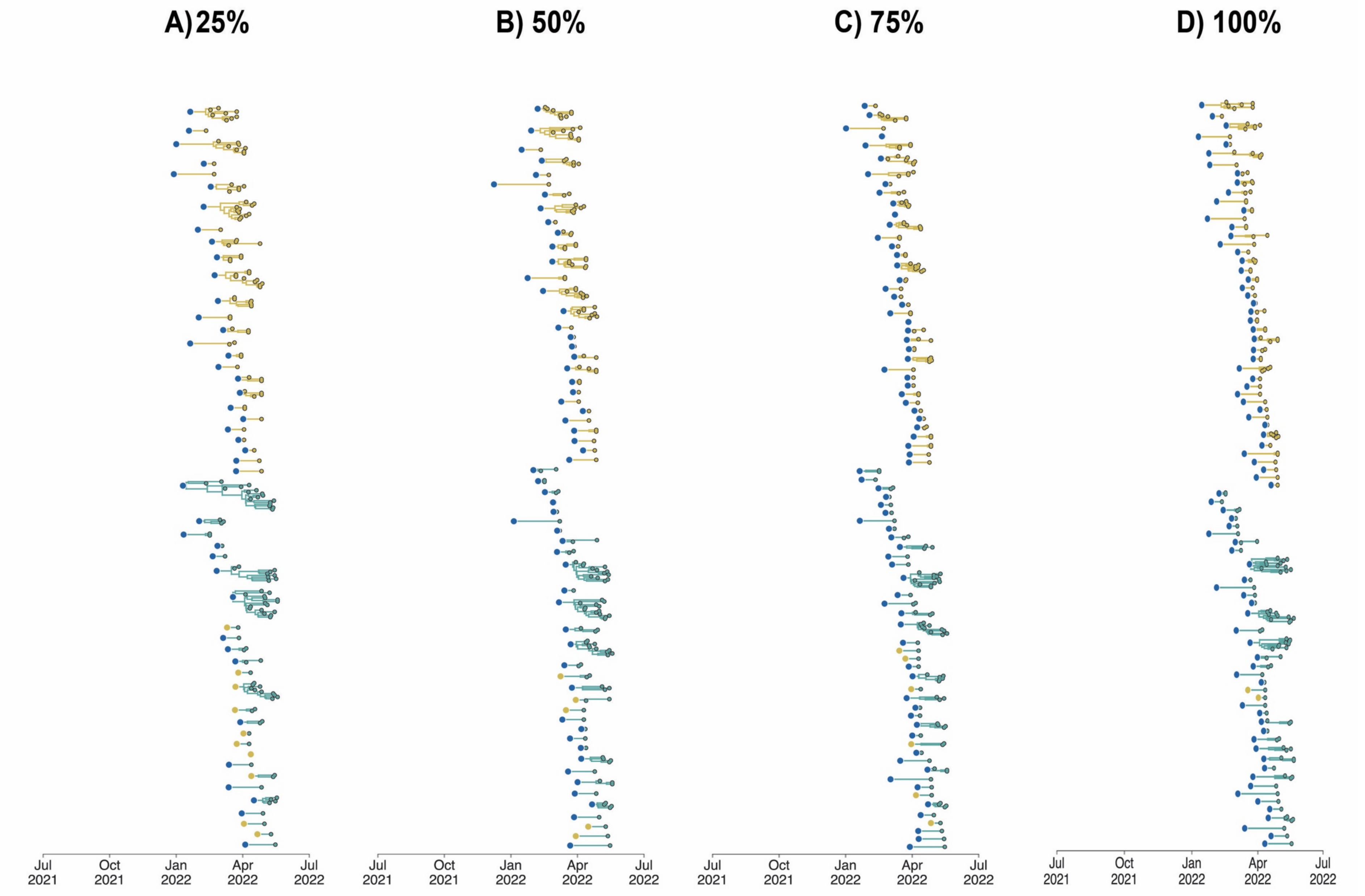


**Figure S25:** Exploded tree view of MCC trees for each tree in the three-state titration analysis. Percentage refers to the percentage of all available wild sequences used in the given analysis. Subtrees represent the traversal of a tree from the root to the tip where the state is unchanged from the initial state (given by the large dot on left) to the tips represented by the smaller dots representing continuous chains of transmission within a given state. Colors correspond to the state of a given tip, branch node where blue = wild bird, green = commercial bird, yellow = backyard bird.

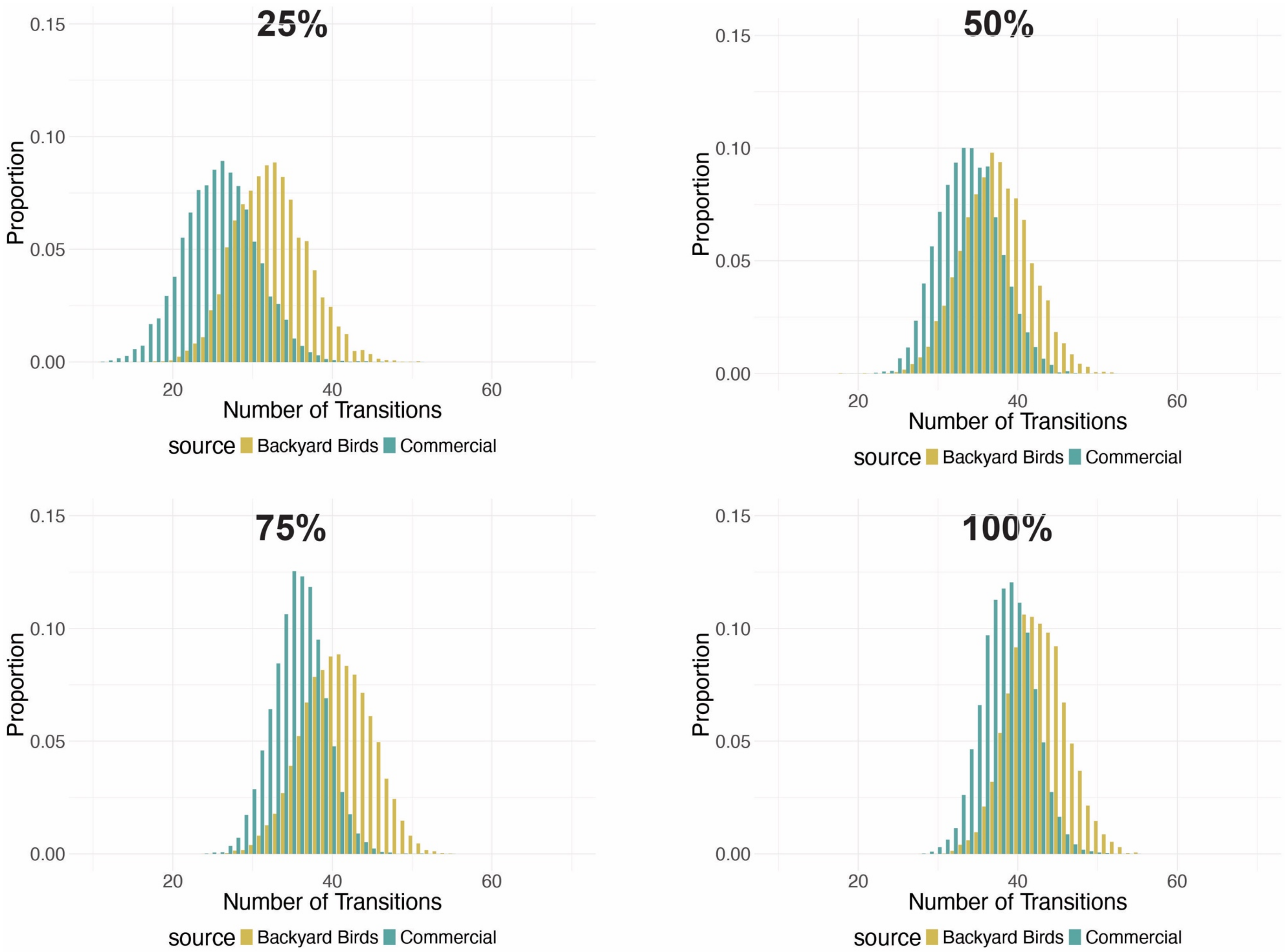


**Figure S26.** Proportion of transitions from Wild to Backyard birds and commercial birds across the posterior trees for each titration. Ratio represents the ratio (backyard:commercial:wild bird) of sequences in the given analysis.

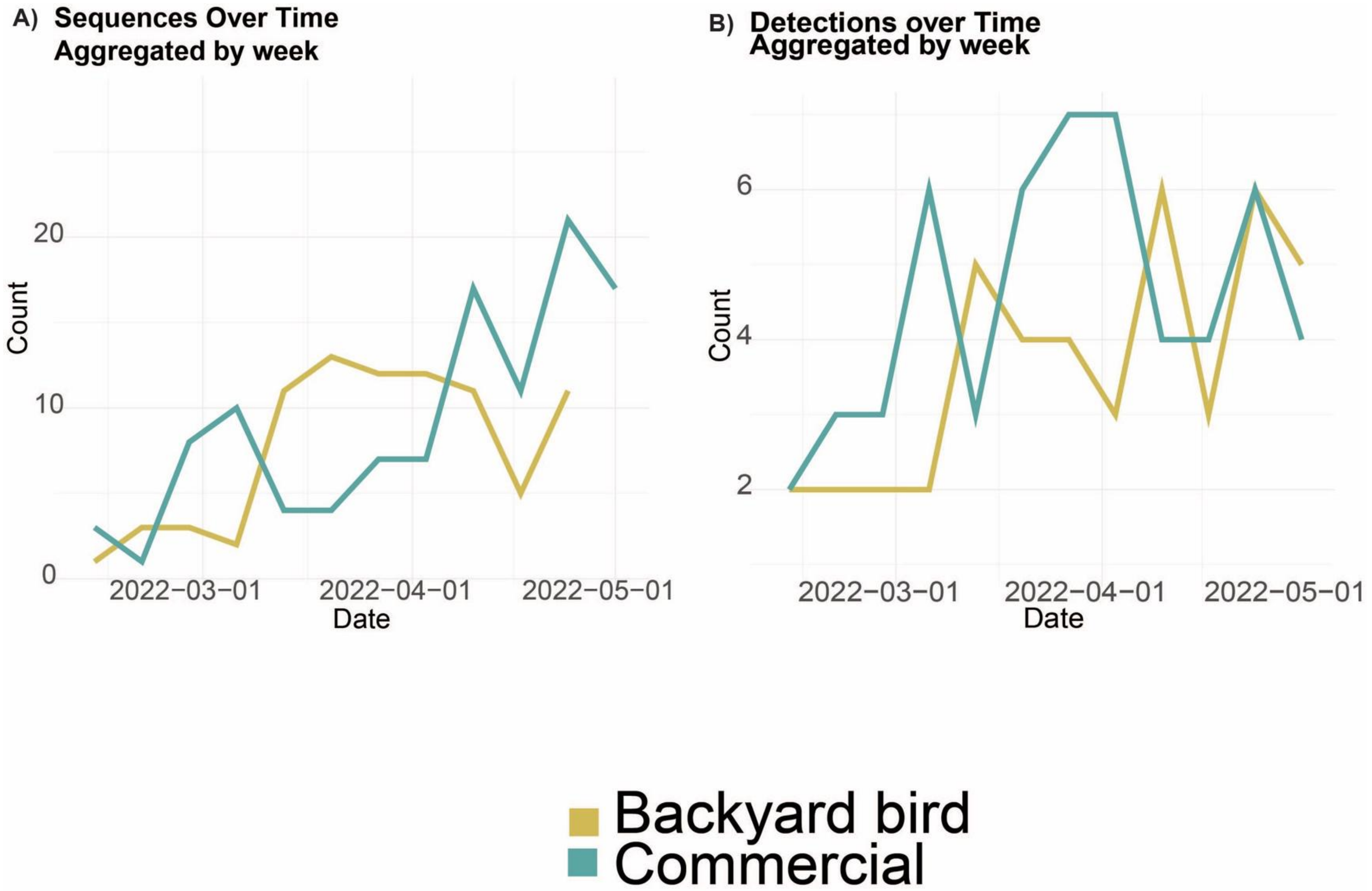


**Figure S27.** A) Number of sequences over time aggregated by week from Backyard birds (yellow) and commercial birds (Green) between 2022-02-12 and 2022-05-01. B) Number of detections (farms/premises) over time aggregated by week from WOAH Non-Poultry (Backyard birds) and WOAH Poultry (commercial birds) between 2022-02-12 and 2022-05-01. C) Number of times that a given record production type for a detection transitioned to a different production type for the next record in chronological order for all available detections.

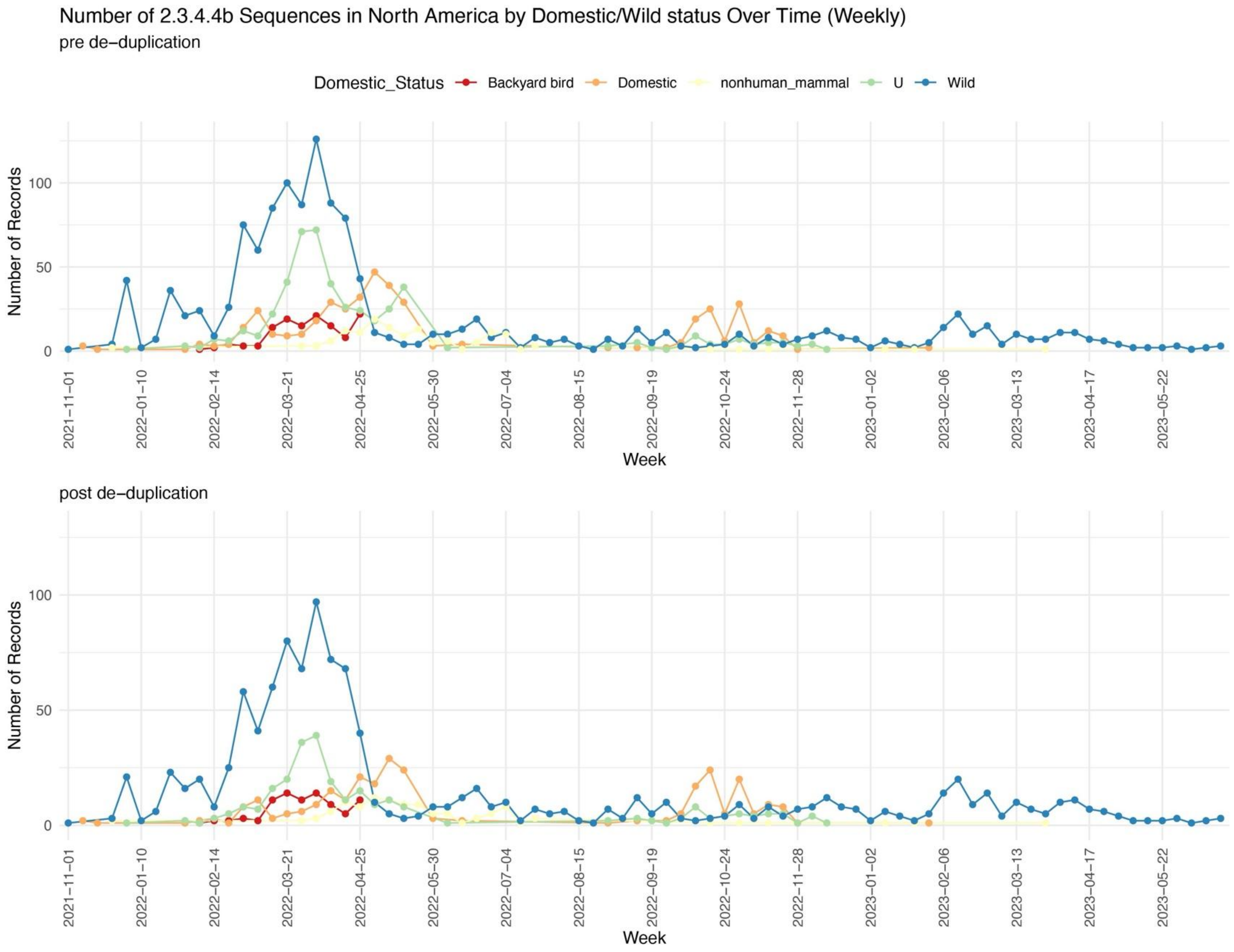


**Figure S28.** Number of sequences by week by domestic/wild status determined by available metadata before and after deduplication of identical sequences occurring on the same day. Post de-duplication total number of sequences 1818.

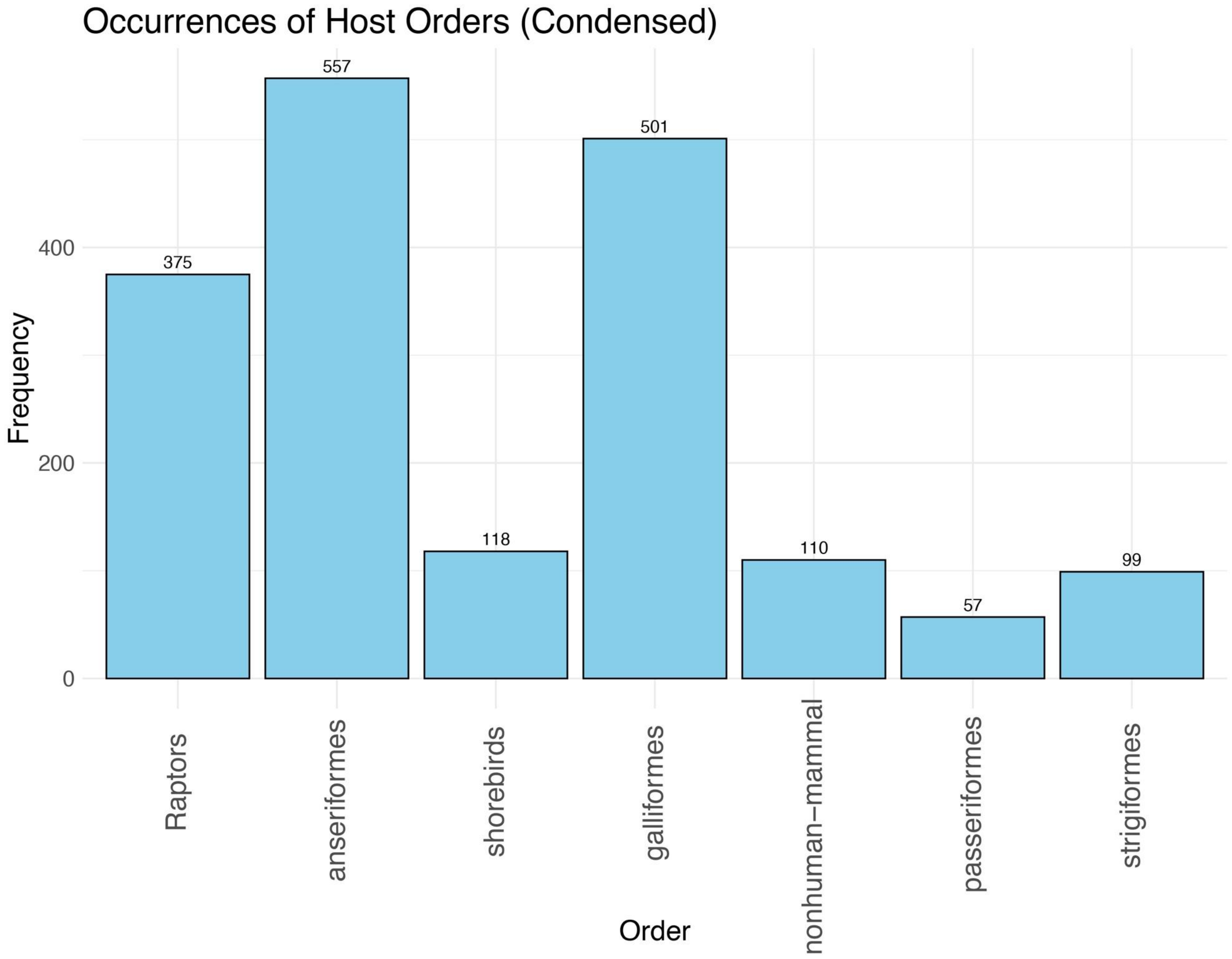


**Figure S29** Number of sequences for each host taxonomic order for available sequence data.

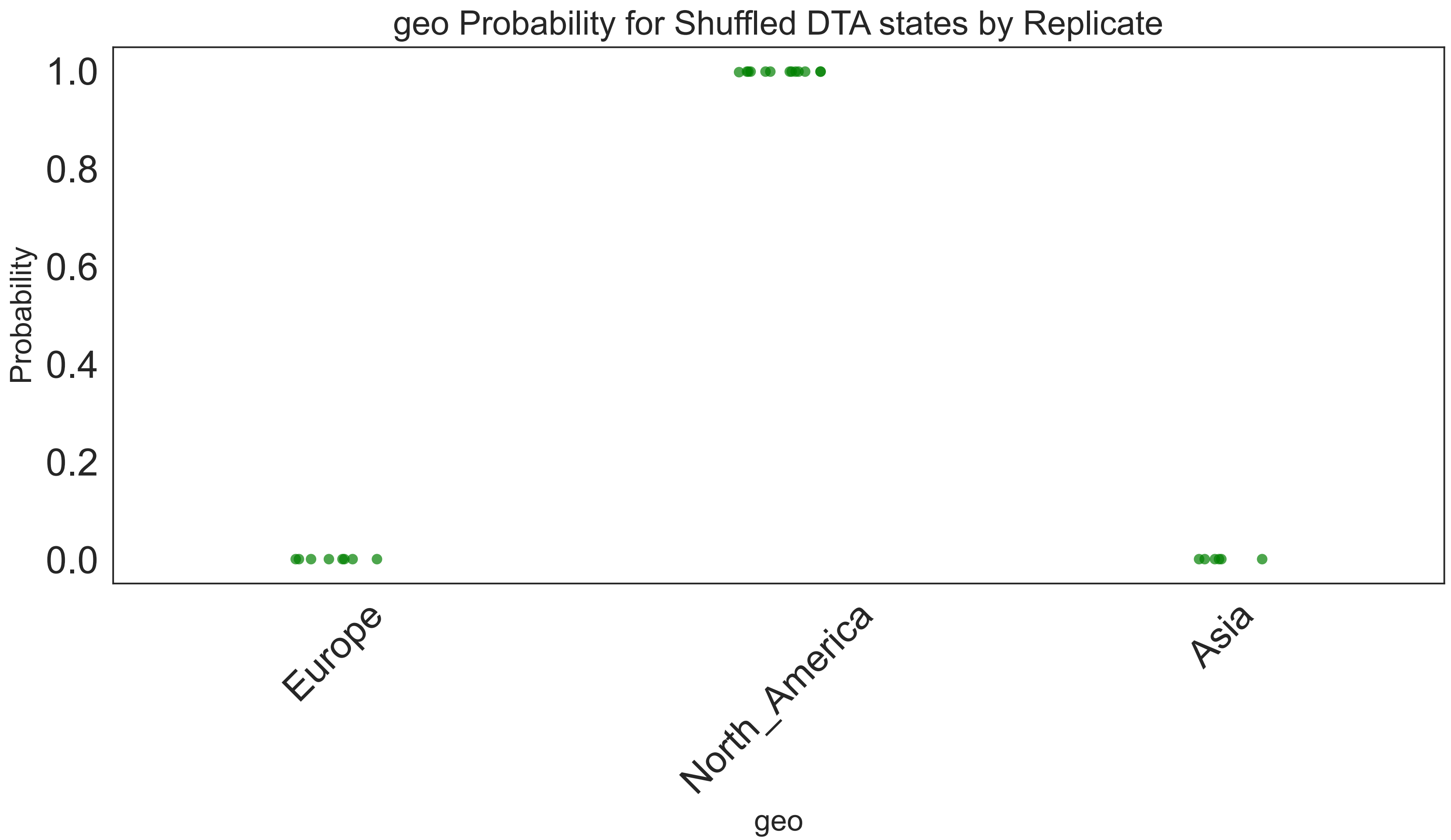


**Figure S30** Root state probabilities for 100 replicates of tip shuffle test for Global introduction dataset.

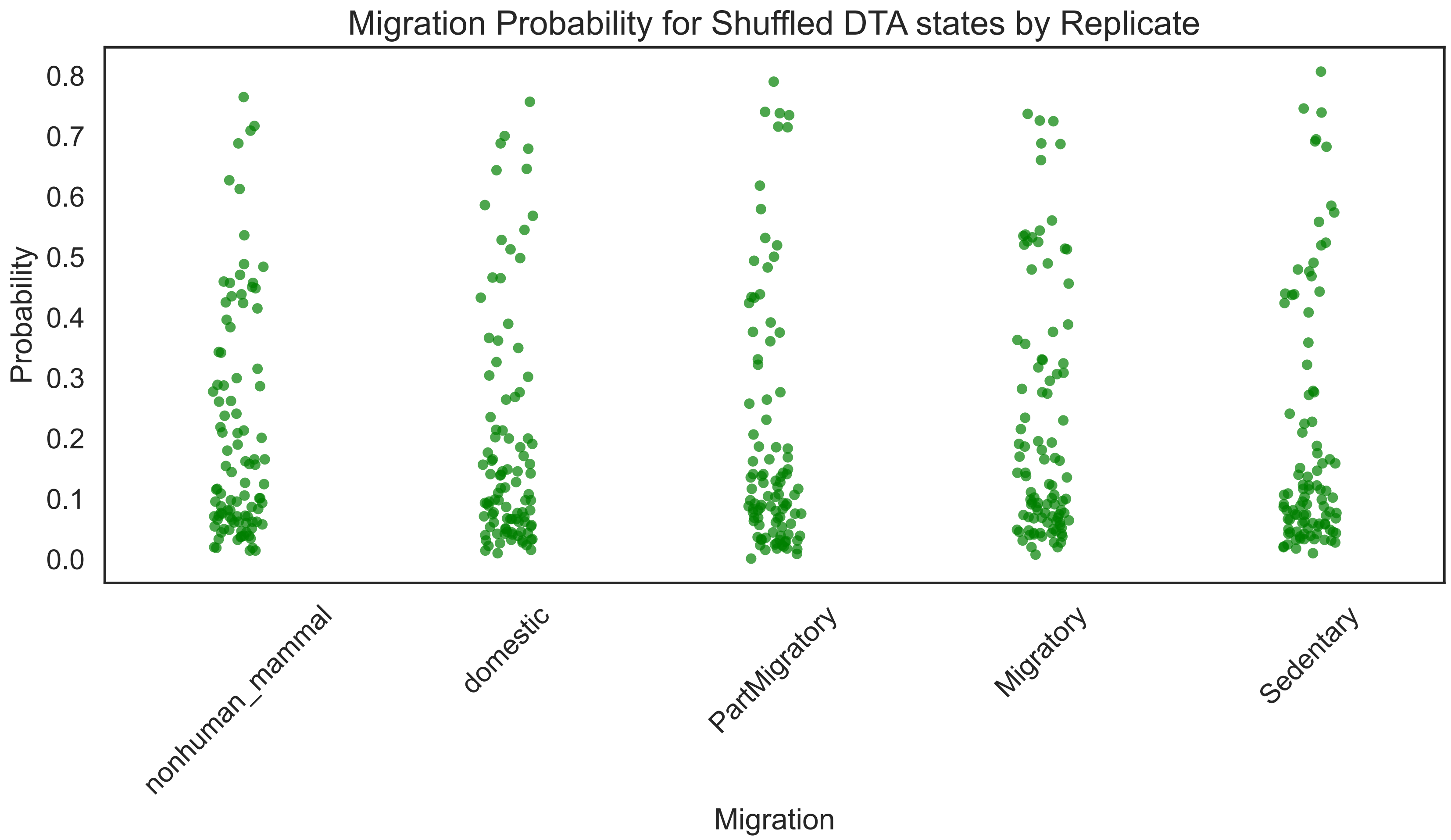


**Figure S31** Root state probabilities for 100 replicates of tip shuffle test for Migratory behavior dataset.

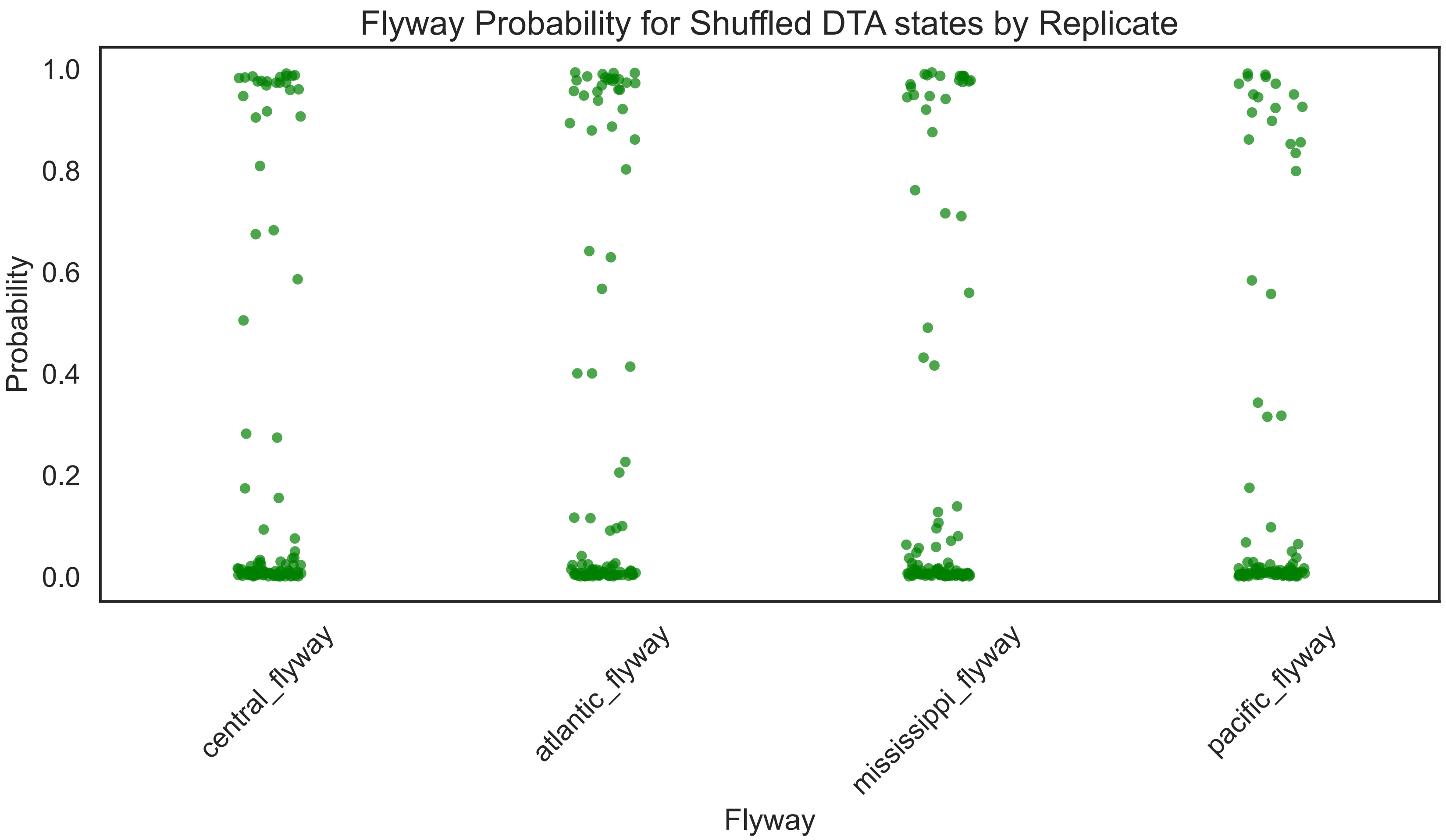


**Figure S32** Root state probabilities for 100 replicates of tip shuffle test for flyway dataset.

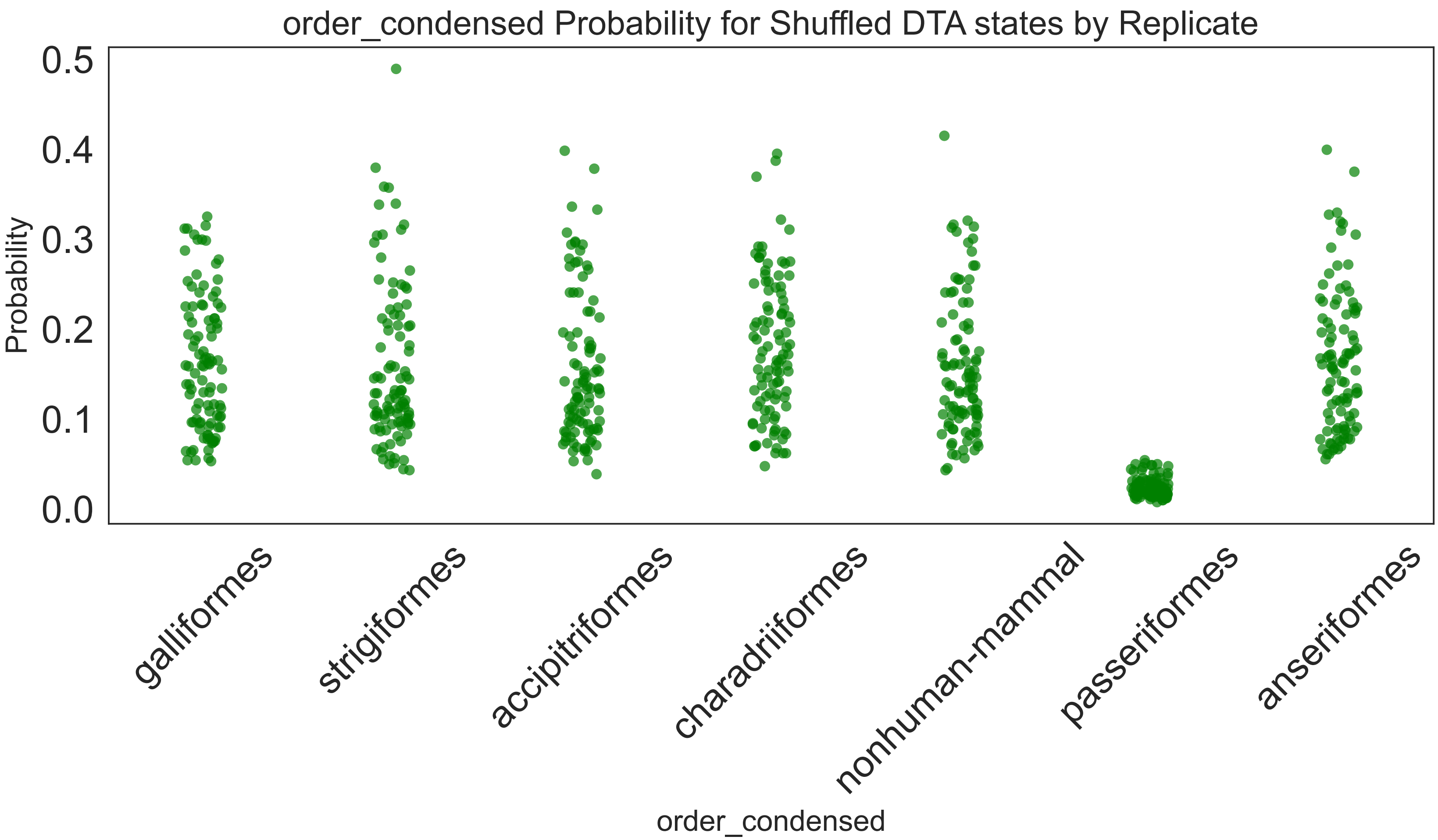


**Figure S33** Root state probabilities for 100 replicates of tip shuffle test for order combined

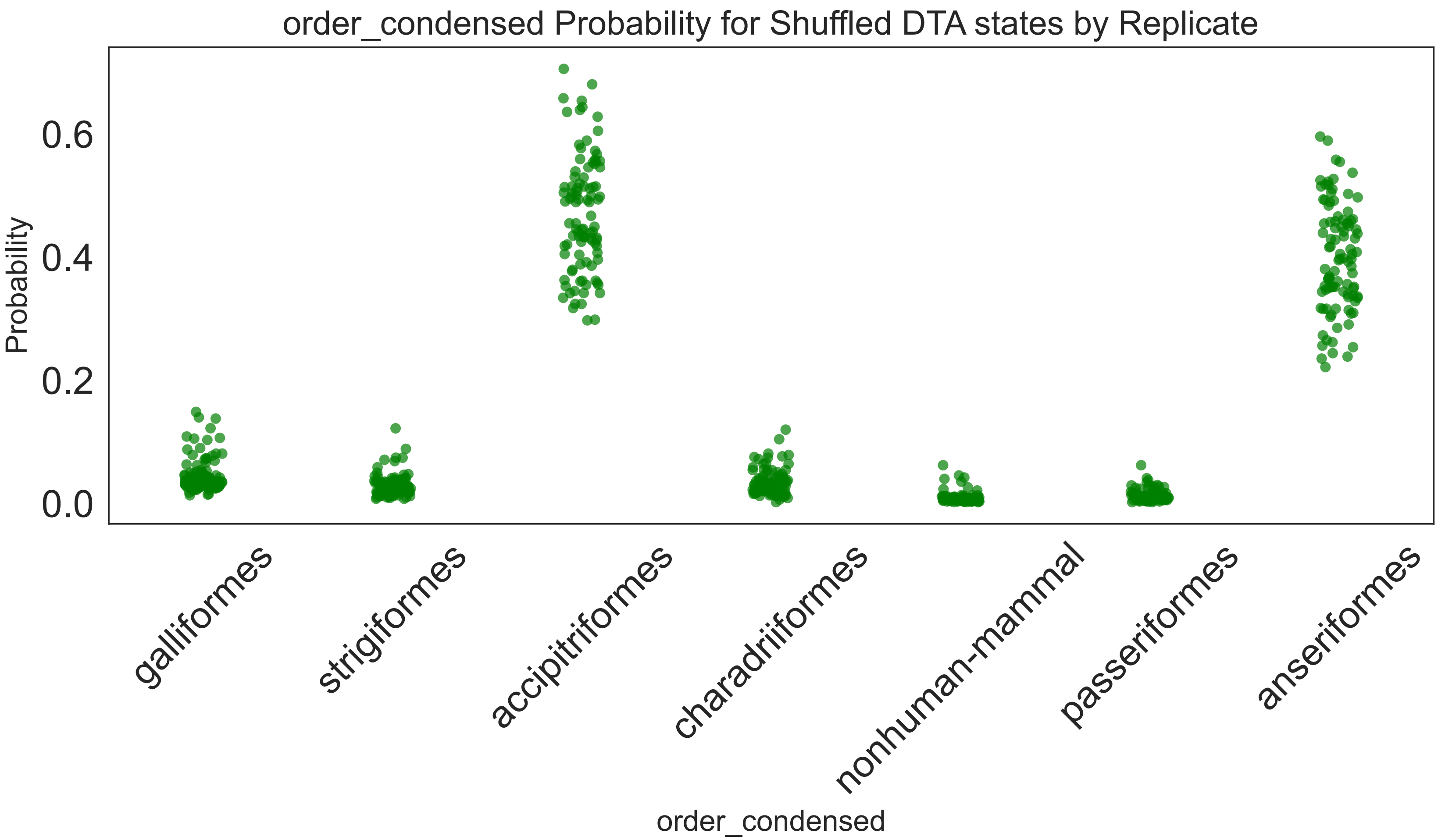


**Figure S34** Root state probabilities for 100 replicates of tip shuffle test for order prop combined dataset.

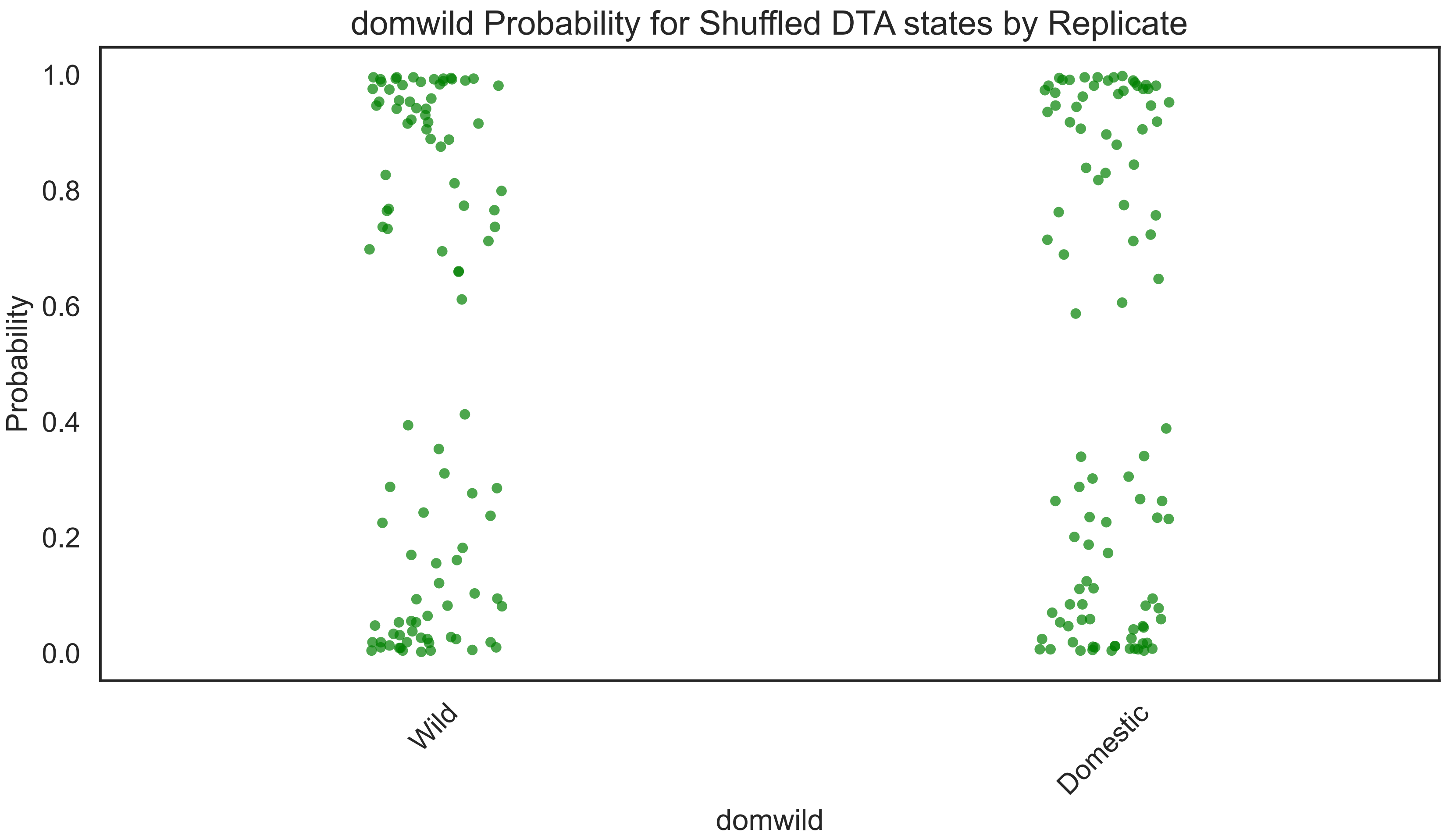


**Figure S35** Root state probabilities for 100 replicates of tip shuffle test for domestic wild (1:1) dataset.

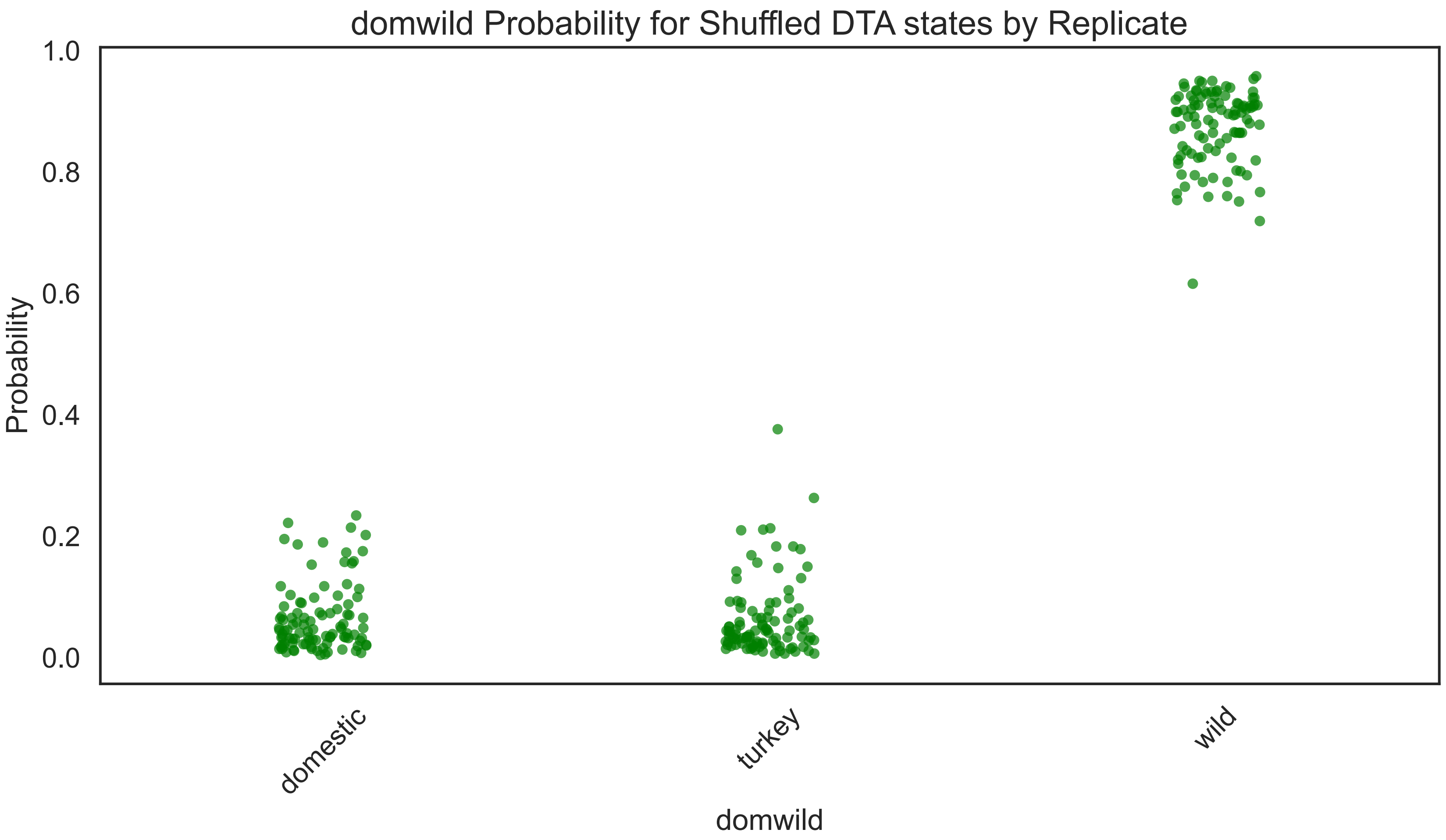


**Figure S36** Root state probabilities for 100 replicates of tip shuffle test for domestic wild turkey (1:1:1) dataset.

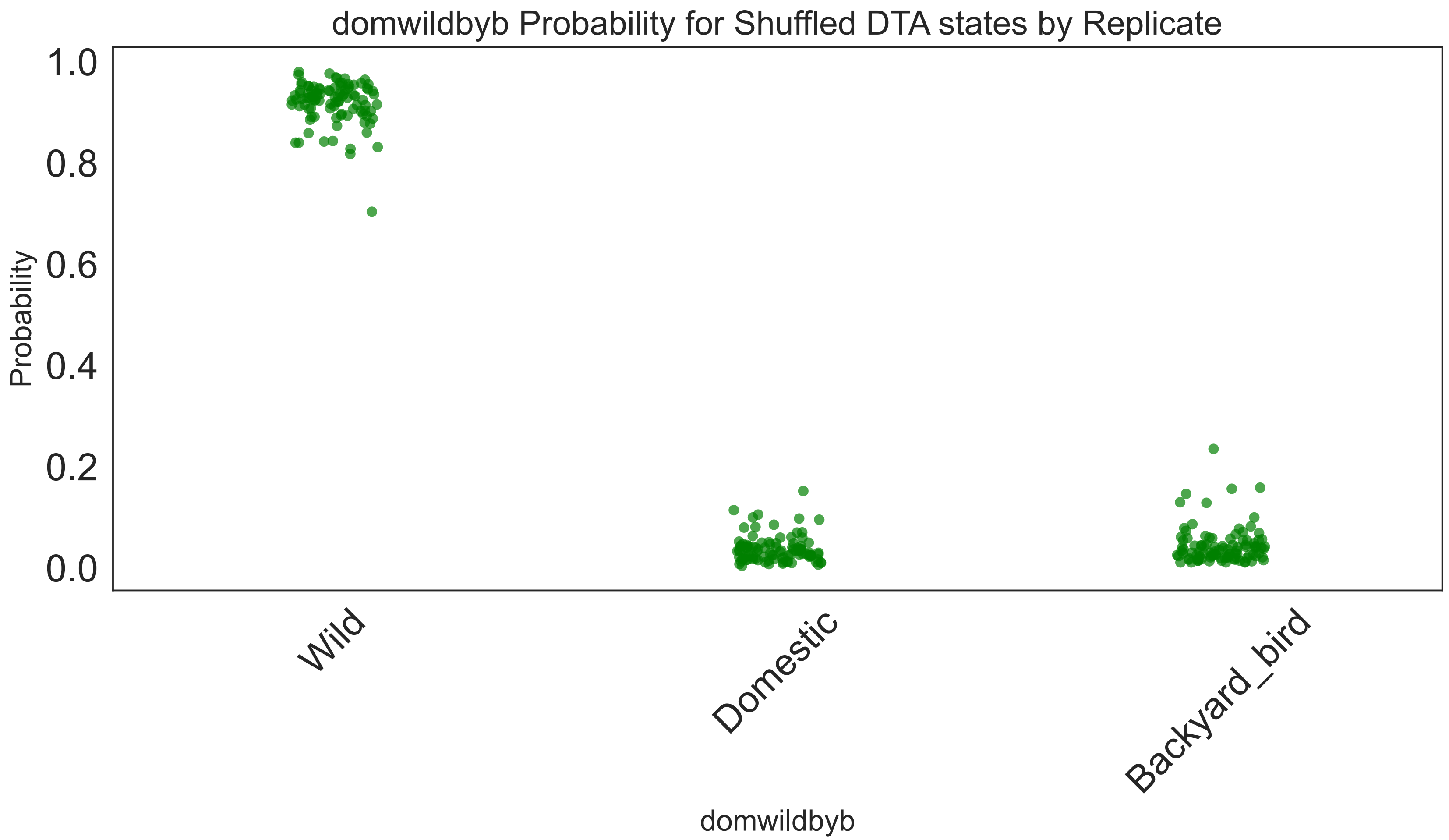


**Figure S37** Root state probabilities for 100 replicates of tip shuffle test for domestic wild backyard bird dataset (1:1 domestic:backyard bird + 25% all wild).

### Supplementary Tables

| From | To | mean | median | l95hpd | h95hpd | BAYES FACTOR | POSTERIOR PROBABILITY | mean Markov Jumps | Markov Jumps 95% lhpd | Markov Jumps 95% hhpd | mean Markov Jump / tree height | Markov jumps / tree hieght lower 95%hpd | Markov jumps / tree hieght upper 95%hpd | Markov Jump / tree length | Markov Jump / tree length lower 95% hpd | Markov Jump / tree length upper 95% hpd |
|---|---|---|---|---|---|---|---|---|---|---|---|---|---|---|---|---|
| **Atlantic** | **Mississippi** | **0.742** | **4.080** | **27000.000** | **1.000** | **2.305** | **2.186** | **70.232** | **58.000** | **81.000** | **37.340** | **30.837** | **43.065** | **0.345** | **0.285** | **0.398** |
| Atlantic | Pacific | 0.004 | 2.349 | 9.672 | 0.763 | 0.505 | 0.223 | 3.163 | 1.000 | 5.000 | 1.681 | 0.532 | 2.658 | 0.016 | 0.005 | 0.025 |
| **Central** | **Pacific** | **0.641** | **3.480** | **27000.000** | **1.000** | **1.935** | **1.823** | **24.690** | **15.000** | **34.000** | **13.127** | **7.975** | **18.077** | **0.121** | **0.074** | **0.167** |
| **Central** | **Mississippi** | **0.210** | **1.746** | **27000.000** | **1.000** | **0.899** | **0.818** | **16.657** | **10.000** | **23.000** | **8.856** | **5.317** | **12.228** | **0.082** | **0.049** | **0.113** |
| **Mississippi** | **Central** | **1.670** | **7.873** | **27000.000** | **1.000** | **4.687** | **4.491** | **105.895** | **90.000** | **121.000** | **56.301** | **47.850** | **64.332** | **0.520** | **0.442** | **0.595** |
| **Mississippi** | **Atlantic** | **0.111** | **1.529** | **95.912** | **0.970** | **0.724** | **0.643** | **2.689** | **1.000** | **7.000** | **1.430** | **0.532** | **3.722** | **0.013** | **0.005** | **0.034** |
| Pacific | Central | 0.139 | 1.418 | 27000.000 | 1.000 | 0.698 | 0.627 | 21.134 | 15.000 | 25.000 | 11.236 | 7.975 | 13.292 | 0.104 | 0.074 | 0.123 |

**Table S1:** Results of BSSVS discrete trait analysis for USFWS flyways. Table is in order of highest mean transition rate to lowest with rates having BF > 100 bolded. Rates with at least a BF > 3 are shown.

| From | To | mean | median | l95hpd | h95hpd | BAYES_FACTOR | POSTERIOR PROBABILITY |
|---|---|---|---|---|---|---|---|
| **group1_combined_above49N** | **group2_northernus_southerncanada_42N_49N** | **0.72298544** | **0.66041858** | **0.14856531** | **1.39344548** | **29700** | **1** |
| **group1_combined_above49N** | **group4_southernus_below36N** | **0.86767129** | **0.80234959** | **0.20925934** | **1.6251426** | **29700** | **1** |
| **group3_centralus_36N_42N** | **group2_northernus_southerncanada_42N_49N** | **1.82410787** | **1.69980647** | **0.48516246** | **3.46284291** | **29700** | **1** |
| **group4_southernus_below36N** | **group2_northernus_southerncanada_42N_49N** | **0.82898677** | **0.76352712** | **0.19578435** | **1.64072213** | **29700** | **1** |
| **group4_southernus_below36N** | **group3_centralus_36N_42N** | **1.92837598** | **1.82155582** | **0.63907759** | **3.45480903** | **29700** | **1** |
| **group2_northernus_southerncanada_42N_49N** | **group3_centralus_36N_42N** | **1.31353336** | **1.20124055** | **0.26230058** | **2.59867265** | **9898** | **0.999697** |
| **group2_northernus_southerncanada_42N_49N** | **group1_combined_above49N** | **1.05557566** | **0.97455431** | **0.23064274** | **2.03878799** | **799.783784** | **0.996263** |
| group2_northernus_southerncanada_42N_49N | group4_southernus_below36N | 0.60581523 | 0.53262685 | 0.04471526 | 1.32288619 | 86.7371601 | 0.96656903 |
| group3_centralus_36N_42N | group1_combined_above49N | 0.52887832 | 0.3500056 | 0.00020955 | 1.69816282 | 7.69992795 | 0.71962428 |
| group3_centralus_36N_42N | group4_southernus_below36N | 0.76006982 | 0.4825803 | 2.47E-05 | 2.51017546 | 2.44809244 | 0.44934855 |
| group1_combined_above49N | group3_centralus_36N_42N | 0.65898606 | 0.34038264 | 2.89E-05 | 2.39739077 | 2.28899573 | 0.43278457 |
| group4_southernus_below36N | group1_combined_above49N | 0.85125837 | 0.51546685 | 4.59E-05 | 2.79417744 | 0.59513435 | 0.16553883 |

**Table S2**: Results of BSSVS discrete trait analysis for geographic group based on latitude. Table is in order of highest mean transition rate to lowest with rates having BF > 100 bolded. Rates with at least a BF > 3 are shown.

| From | To | BAYES_FACTOR | POSTERIOR PROBABILITY | mean | median | l95hpd | h95hpd |
|---|---|---|---|---|---|---|---|
| **Migration** | **PartMigration** | **36000** | **1** | **2.00874182** | **1.90919107** | **0.73749557** | **3.48482027** |
| **Migration** | **Sedentary** | **181.587629** | **0.97844684** | **1.04729841** | **0.95439458** | **0.11917316** | **2.10127971** |
| **Migration** | **Domestic** | **36000** | **1** | **1.33497394** | **1.25320356** | **0.40376184** | **2.38054574** |
| **Migration** | **Nonhumanmammal** | **400.539326** | **0.99011221** | **1.19631674** | **1.12017969** | **0.24933207** | **2.33312444** |
| PartMigration | Sedentary | 97.9943343 | 0.96078214 | 0.87098186 | 0.77876747 | 0.07788287 | 1.84752126 |
| PartMigration | Domestic | 0.52141153 | 0.11532052 | 0.91600762 | 0.58774557 | 0.00061945 | 2.89768449 |
| PartMigration | Nonhumanmammal | 1.13316225 | 0.22075325 | 0.8826551 | 0.55567327 | 0.00047205 | 2.81096245 |
| Sedentary | Domestic | 0.53450882 | 0.11787579 | 0.95396382 | 0.62897596 | 0.00061559 | 2.96816781 |
| Sedentary | Nonhumanmammal | 3.036154 | 0.43150761 | 0.779748 | 0.52251605 | 3.72E-06 | 2.41343409 |
| Domestic | Nonhumanmammal | 20.3929539 | 0.83601822 | 0.93032214 | 0.80834525 | 0.01262889 | 2.11755253 |
| PartMigration | Migration | 2.21508717 | 0.35640484 | 0.75888425 | 0.4470211 | 0.0005076 | 2.5426762 |
| Sedentary | Migration | 36000 | 1 | 1.44051268 | 1.35975899 | 0.48396807 | 2.58706142 |
| Domestic | Migration | 1.43128677 | 0.26352628 | 0.83338968 | 0.52381554 | 5.85E-05 | 2.68385089 |
| Nonhumanmammal | Migration | 0.50275138 | 0.11165426 | 0.89103885 | 0.58370738 | 8.10E-05 | 2.77166192 |
| Sedentary | PartMigration | 5.62416466 | 0.58437951 | 0.7707354 | 0.57152876 | 5.10E-05 | 2.17327905 |
| Domestic | PartMigration | 10.9891757 | 0.73314076 | 0.75017909 | 0.58957738 | 0.00020665 | 1.96900017 |
| Nonhumanmammal | PartMigration | 140.594378 | 0.97233641 | 0.89119182 | 0.80402919 | 0.03583859 | 1.84688931 |
| Domestic | Sedentary | 4.18272727 | 0.51116543 | 0.88474002 | 0.68014107 | 0.00151883 | 2.41678285 |
| Nonhumanmammal | Sedentary | 510.342857 | 0.99222309 | 1.19024247 | 1.11177603 | 0.24962462 | 2.29915023 |
| Nonhumanmammal | Domestic | 4.69661836 | 0.54005111 | 0.64654853 | 0.41283259 | 0.00046807 | 2.10029534 |

**Table S3:** Results of BSSVS discrete trait analysis for Migratory Behavior. Table is in order of highest mean transition rate to lowest with rates having BF > 100 bolded. Rates with at least a BF > 3 are shown.

| From | To | mean | median | l95hpd | h95hpd | BAYES FACTOR | POSTERIOR PROB | mean Markov Jumps | Markov jumps lower 95%hpd | Markov jumps upper 95%hpd | Markov Jumps / tree height | Markov jumps / tree hieght lower 95%hpd | Markov jumps / tree hiegh t upper 95% hpd | Markov Jump / tree length | Markov Jump / tree length lower 95% hpd | Markov Jump / tree length upper 95% hpd |
|---|---|---|---|---|---|---|---|---|---|---|---|---|---|---|---|---|
| **Anseriformes** | **Galliformes** | **4.494** | **4.374** | **1.842** | **7.208** | **1691.250** | **0.996** | **49.950** | **26.000** | **73.000** | **17.808** | **9.269** | **26.025** | **0.493** | **0.257** | **0.721** |
| **Shorebirds** | **Nonhuman-Mammal** | **2.089** | **1.970** | **0.640** | **3.620** | **537.120** | **0.989** | **17.035** | **9.000** | **25.000** | **6.073** | **3.209** | **8.913** | **0.168** | **0.089** | **0.247** |
| **Shorebirds** | **Raptors** | **1.787** | **1.668** | **0.358** | **3.298** | **245.444** | **0.976** | **14.651** | **5.000** | **24.000** | **5.223** | **1.783** | **8.556** | **0.145** | **0.049** | **0.237** |
| **Anseriformes** | **Strigiformes** | **3.406** | **3.238** | **1.239** | **6.141** | **232.211** | **0.975** | **37.889** | **15.000** | **64.000** | **13.508** | **5.348** | **22.816** | **0.374** | **0.148** | **0.632** |
| **Galliformes** | **Strigiformes** | **3.015** | **2.871** | **0.853** | **5.365** | **193.676** | **0.970** | **28.517** | **4.000** | **47.000** | **10.166** | **1.426** | **16.756** | **0.282** | **0.039** | **0.464** |
| **Galliformes** | **Nonhuman-Mammal** | **2.648** | **2.524** | **0.779** | **4.771** | **170.338** | **0.966** | **24.653** | **5.000** | **38.000** | **8.789** | **1.783** | **13.547** | **0.243** | **0.049** | **0.375** |
| **Shorebirds** | **Passeriformes** | **1.219** | **1.103** | **0.268** | **2.428** | **150.069** | **0.962** | **9.351** | **3.000** | **14.000** | **3.334** | **1.070** | **4.991** | **0.092** | **0.030** | **0.138** |
| **Galliformes** | **Anseriformes** | **2.472** | **2.296** | **0.453** | **4.882** | **146.562** | **0.961** | **22.044** | **2.000** | **42.000** | **7.859** | **0.713** | **14.973** | **0.218** | **0.020** | **0.415** |
| **Anseriformes** | **Shorebirds** | **1.775** | **1.662** | **0.330** | **3.411** | **143.209** | **0.960** | **18.978** | **5.000** | **34.000** | **6.766** | **1.783** | **12.121** | **0.187** | **0.049** | **0.336** |
| **Anseriformes** | **Raptors** | **2.527** | **2.344** | **0.383** | **4.865** | **127.118** | **0.955** | **27.702** | **6.000** | **51.000** | **9.876** | **2.139** | **18.182** | **0.273** | **0.059** | **0.504** |
| **Galliformes** | **Passeriformes** | **2.043** | **1.967** | **0.469** | **3.785** | **117.436** | **0.951** | **18.563** | **5.000** | **30.000** | **6.618** | **1.783** | **10.695** | **0.183** | **0.049** | **0.296** |
| Strigiformes | Passeriformes | 1.574 | 1.455 | 0.183 | 3.095 | 81.600 | 0.932 | 11.313 | 2.000 | 20.000 | 4.033 | 0.713 | 7.130 | 0.112 | 0.020 | 0.197 |
| Raptors | Nonhuman-Mammal | 1.797 | 1.633 | 0.240 | 3.673 | 71.148 | 0.922 | 14.656 | 3.000 | 27.000 | 5.225 | 1.070 | 9.626 | 0.145 | 0.030 | 0.267 |
| Strigiformes | Raptors | 1.919 | 1.753 | 0.235 | 3.942 | 61.218 | 0.911 | 13.627 | 3.000 | 25.000 | 4.858 | 1.070 | 8.913 | 0.135 | 0.030 | 0.247 |
| Nonhuman-Mammal | Anseriformes | 2.236 | 2.039 | 0.207 | 4.517 | 53.815 | 0.900 | 15.836 | 2.000 | 29.000 | 5.646 | 0.713 | 10.339 | 0.156 | 0.020 | 0.286 |
| Raptors | Strigiformes | 1.710 | 1.539 | 0.178 | 3.444 | 51.534 | 0.896 | 13.861 | 2.000 | 25.000 | 4.941 | 0.713 | 8.913 | 0.137 | 0.020 | 0.247 |
| Raptors | Shorebirds | 1.681 | 1.555 | 0.169 | 3.486 | 47.881 | 0.889 | 13.376 | 1.000 | 24.000 | 4.769 | 0.357 | 8.556 | 0.132 | 0.010 | 0.237 |
| Galliformes | Raptors | 2.009 | 1.857 | 0.195 | 4.059 | 41.642 | 0.874 | 19.161 | 1.000 | 35.000 | 6.831 | 0.357 | 12.478 | 0.189 | 0.010 | 0.346 |
| Raptors | Anseriformes | 1.873 | 1.719 | 0.182 | 3.942 | 38.518 | 0.865 | 14.530 | 1.000 | 26.000 | 5.180 | 0.357 | 9.269 | 0.143 | 0.010 | 0.257 |
| Galliformes | Shorebirds | 1.479 | 1.311 | 0.129 | 3.152 | 33.818 | 0.849 | 12.235 | 2.000 | 23.000 | 4.362 | 0.713 | 8.200 | 0.121 | 0.020 | 0.227 |
| Shorebirds | Anseriformes | 1.209 | 0.952 | 0.013 | 3.149 | 17.014 | 0.739 | 7.472 | 1.000 | 15.000 | 2.664 | 0.357 | 5.348 | 0.074 | 0.010 | 0.148 |
| Strigiformes | Shorebirds | 1.487 | 1.187 | 0.016 | 3.692 | 11.588 | 0.659 | 9.110 | 1.000 | 18.000 | 3.248 | 0.357 | 6.417 | 0.090 | 0.010 | 0.178 |
| Strigiformes | Nonhuman-Mammal | 1.321 | 1.040 | 0.005 | 3.421 | 10.660 | 0.640 | 7.089 | 1.000 | 14.000 | 2.527 | 0.357 | 4.991 | 0.070 | 0.010 | 0.138 |
| Raptors | Galliformes | 1.314 | 1.030 | 0.002 | 3.302 | 9.807 | 0.620 | 8.312 | 1.000 | 17.000 | 2.963 | 0.357 | 6.061 | 0.082 | 0.010 | 0.168 |
| Nonhuman-Mammal | Galliformes | 1.553 | 1.262 | 0.009 | 3.804 | 8.727 | 0.593 | 7.869 | 1.000 | 16.000 | 2.805 | 0.357 | 5.704 | 0.078 | 0.010 | 0.158 |
| Anseriformes | Nonhuman-Mammal | 1.574 | 1.334 | 0.010 | 3.769 | 8.399 | 0.583 | 16.845 | 1.000 | 45.000 | 6.005 | 0.357 | 16.043 | 0.166 | 0.010 | 0.444 |

| | | | | | | | | | | | | | | | | |
|---|---|---|---|---|---|---|---|---|---|---|---|---|---|---|---|---|
| Passeriformes | Raptors | 1.494 | 1.206 | 0.006 | 3.750 | 8.353 | 0.582 | 5.655 | 1.000 | 12.000 | 2.016 | 0.357 | 4.278 | 0.056 | 0.010 | 0.118 |
| Nonhuman-Mammal | Shorebirds | 1.369 | 1.048 | 0.009 | 3.769 | 6.080 | 0.503 | 6.318 | 1.000 | 14.000 | 2.252 | 0.357 | 4.991 | 0.062 | 0.010 | 0.138 |
| Nonhuman-Mammal | Raptors | 1.383 | 1.076 | 0.004 | 3.619 | 6.069 | 0.503 | 6.761 | 1.000 | 15.000 | 2.410 | 0.357 | 5.348 | 0.067 | 0.010 | 0.148 |
| Nonhuman-Mammal | Strigiformes | 1.370 | 1.037 | 0.002 | 3.771 | 6.059 | 0.502 | 6.575 | 1.000 | 14.000 | 2.344 | 0.357 | 4.991 | 0.065 | 0.010 | 0.138 |
| Passeriformes | Anseriformes | 1.512 | 1.156 | 0.037 | 4.284 | 5.963 | 0.498 | 5.226 | 1.000 | 11.000 | 1.863 | 0.357 | 3.922 | 0.052 | 0.010 | 0.109 |
| Strigiformes | Galliformes | 1.434 | 1.136 | 0.005 | 3.733 | 5.487 | 0.478 | 8.253 | 1.000 | 18.000 | 2.942 | 0.357 | 6.417 | 0.081 | 0.010 | 0.178 |
| Anseriformes | Passeriformes | 1.441 | 1.124 | 0.014 | 3.748 | 5.066 | 0.458 | 13.073 | 1.000 | 28.000 | 4.660 | 0.357 | 9.982 | 0.129 | 0.010 | 0.276 |

**Table S4:** Results of BSSVS discrete trait analysis for combined results of equal host schema. Table is in order of highest mean transition rate to lowest with rates having BF > 100 bolded. Rates with at least a BF > 3 are shown.

| From | To | mean | median | l95hpd | h95hpd | BAYES FACTOR | POSTERIOR PROBABILITY | mean Markov Jumps | Markov Jumps 95% lhpd | Markov Jumps 95% hhpd | mean Markov Jump / tree height | Markov jumps / tree hieght lower 95%hpd | Markov jumps / tree hieght upper 95%hpd | Markov Jump / tree length | Markov Jump / tree length lower 95% hpd | Markov Jump / tree length upper 95% hpd |
|---|---|---|---|---|---|---|---|---|---|---|---|---|---|---|---|---|
| **Anseriformes** | **Galliformes** | **4.427** | **4.291** | **2.201** | **6.781** | **18006.000** | **1.000** | **47.738** | **30.000** | **61.000** | **21.275** | **13.370** | **27.186** | **0.345** | **0.217** | **0.441** |
| **Anseriformes** | **Nonhuman-Mammal** | **0.911** | **0.856** | **0.183** | **1.659** | **321.491** | **0.982** | **12.112** | **1.000** | **27.000** | **5.398** | **0.446** | **12.033** | **0.088** | **0.007** | **0.195** |
| **Anseriformes** | **Passeriformes** | **0.680** | **0.626** | **0.158** | **1.296** | **714.480** | **0.992** | **14.041** | **5.000** | **23.000** | **6.258** | **2.228** | **10.250** | **0.102** | **0.036** | **0.166** |
| **Anseriformes** | **Raptors** | **5.885** | **5.711** | **2.805** | **9.178** | **18006.000** | **1.000** | **85.675** | **69.000** | **99.000** | **38.183** | **30.751** | **44.122** | **0.620** | **0.499** | **0.716** |
| **Anseriformes** | **Shorebirds** | **1.647** | **1.565** | **0.359** | **3.094** | **1379.538** | **0.996** | **25.412** | **11.000** | **43.000** | **11.325** | **4.902** | **19.164** | **0.184** | **0.080** | **0.311** |
| **Anseriformes** | **Strigiformes** | **1.508** | **1.436** | **0.594** | **2.531** | **18006.000** | **1.000** | **27.070** | **12.000** | **40.000** | **12.064** | **5.348** | **17.827** | **0.196** | **0.087** | **0.289** |
| Galliformes | Anseriformes | 1.391 | 1.101 | 0.002 | 3.668 | 6.371 | 0.515 | 7.856 | 1.000 | 19.000 | 3.501 | 0.446 | 8.468 | 0.057 | 0.007 | 0.138 |
| Galliformes | Nonhuman-Mammal | 1.131 | 0.842 | 0.001 | 3.323 | 3.880 | 0.393 | 4.239 | 1.000 | 14.000 | 1.889 | 0.446 | 6.239 | 0.031 | 0.007 | 0.101 |
| Galliformes | Raptors | 1.241 | 0.924 | 0.000 | 3.456 | 3.555 | 0.372 | 8.394 | 1.000 | 18.000 | 3.741 | 0.446 | 8.022 | 0.061 | 0.007 | 0.130 |
| Passeriformes | Raptors | 1.106 | 0.775 | 0.002 | 3.219 | 3.627 | 0.377 | 4.334 | 1.000 | 10.000 | 1.931 | 0.446 | 4.457 | 0.031 | 0.007 | 0.072 |
| **Raptors** | **Anseriformes** | **2.951** | **2.753** | **0.880** | **5.633** | **299.288** | **0.980** | **26.316** | **3.000** | **54.000** | **11.728** | **1.337** | **24.066** | **0.190** | **0.022** | **0.391** |
| Raptors | Passeriformes | 1.151 | 0.744 | 0.001 | 3.658 | 3.897 | 0.394 | 11.379 | 2.000 | 22.000 | 5.071 | 0.891 | 9.805 | 0.082 | 0.014 | 0.159 |
| Raptors | Shorebirds | 1.455 | 1.265 | 0.012 | 3.293 | 11.253 | 0.652 | 18.281 | 7.000 | 36.000 | 8.147 | 3.120 | 16.044 | 0.132 | 0.051 | 0.261 |
| Raptors | Strigiformes | 1.210 | 0.998 | 0.005 | 2.960 | 13.121 | 0.686 | 19.129 | 6.000 | 29.000 | 8.525 | 2.674 | 12.925 | 0.138 | 0.043 | 0.210 |
| Shorebirds | Anseriformes | 1.200 | 0.941 | 0.005 | 3.160 | 9.474 | 0.612 | 11.610 | 1.000 | 25.000 | 5.174 | 0.446 | 11.142 | 0.084 | 0.007 | 0.181 |
| Shorebirds | Raptors | 1.226 | 0.904 | 0.000 | 3.462 | 3.199 | 0.348 | 8.708 | 1.000 | 22.000 | 3.881 | 0.446 | 9.805 | 0.063 | 0.007 | 0.159 |

**Table S5:** Results of BSSVS discrete trait analysis for combined results of proportional schema. Table is in order of highest mean transition rate to lowest with rates having BF > 100 bolded. Rates with at least a BF > 3 are shown.

| From | To | mean | median | l95hpd | h95hpd | BAYES FACTOR | POSTERIOR PROBABILITY | mean Markov Jumps | Markov Jumps 95% lhpd | Markov Jumps 95% hhpd | mean Markov Jump / tree height | Markov jumps / tree hieght lower 95%hpd | Markov jumps / tree hieght upper 95%hpd | Markov Jump / tree length | Markov Jump / tree length lower 95% hpd | Markov Jump / tree length upper 95% hpd |
|---|---|---|---|---|---|---|---|---|---|---|---|---|---|---|---|---|
| **Anseriformes** | **Galliformes** | **4.267** | **4.138** | **1.981** | **6.494** | **702.000** | **0.992** | **69.609** | **53.000** | **81.000** | **24.315** | **18.513** | **28.294** | **0.674** | **0.513** | **0.784** |
| **Anseriformes** | **Strigiformes** | **3.977** | **3.748** | **1.408** | **6.777** | **702.000** | **0.992** | **64.918** | **48.000** | **78.000** | **22.676** | **16.767** | **27.246** | **0.629** | **0.465** | **0.755** |
| **Galliformes** | **Strigiformes** | **2.863** | **2.712** | **0.467** | **4.893** | **151.333** | **0.962** | **8.935** | **1.000** | **19.000** | **3.121** | **0.349** | **6.637** | **0.087** | **0.010** | **0.184** |
| **Galliformes** | **Anseriformes** | **2.761** | **2.501** | **0.598** | **5.132** | **1410.000** | **0.996** | **10.775** | **1.000** | **21.000** | **3.764** | **0.349** | **7.335** | **0.104** | **0.010** | **0.203** |
| **Shorebirds** | **Nonhuman-mammal** | **2.643** | **2.531** | **0.991** | **4.629** | **277.200** | **0.979** | **21.281** | **14.000** | **30.000** | **7.434** | **4.890** | **10.479** | **0.206** | **0.136** | **0.291** |
| **Anseriformes** | **Raptors** | **2.628** | **2.475** | **0.984** | **4.545** | **196.286** | **0.970** | **45.234** | **26.000** | **60.000** | **15.801** | **9.082** | **20.958** | **0.438** | **0.252** | **0.581** |
| Galliformes | Passeriformes | 2.365 | 2.181 | 0.873 | 4.169 | 95.143 | 0.941 | 7.368 | 1.000 | 16.000 | 2.574 | 0.349 | 5.589 | 0.071 | 0.010 | 0.155 |
| **Galliformes** | **Nonhuman-mammal** | **2.267** | **2.122** | **0.492** | **4.185** | **151.333** | **0.962** | **6.759** | **1.000** | **14.000** | **2.361** | **0.349** | **4.890** | **0.065** | **0.010** | **0.136** |
| Raptors | Anseriformes | 2.175 | 2.063 | 0.355 | 4.230 | 64.800 | 0.915 | 11.995 | 3.000 | 22.000 | 4.190 | 1.048 | 7.685 | 0.116 | 0.029 | 0.213 |
| Anseriformes | Nonhuman-mammal | 2.148 | 2.008 | 0.198 | 4.164 | 41.200 | 0.873 | 43.700 | 29.000 | 56.000 | 15.265 | 10.130 | 19.561 | 0.423 | 0.281 | 0.542 |
| **Anseriformes** | **Shorebirds** | **2.024** | **1.879** | **0.387** | **3.799** | **102.923** | **0.945** | **33.690** | **19.000** | **47.000** | **11.768** | **6.637** | **16.417** | **0.326** | **0.184** | **0.455** |
| **Nonhuman - mammal** | **Galliformes** | **1.943** | **1.788** | **0.374** | **4.135** | **135.600** | **0.958** | **7.364** | **1.000** | **13.000** | **2.572** | **0.349** | **4.541** | **0.071** | **0.010** | **0.126** |
| Strigiformes | Raptors | 1.925 | 1.768 | 0.237 | 4.202 | 26.930 | 0.818 | 9.093 | 1.000 | 19.000 | 3.176 | 0.349 | 6.637 | 0.088 | 0.010 | 0.184 |
| **Passeriformes** | **Raptors** | **1.922** | **1.748** | **0.276** | **3.671** | **196.286** | **0.970** | **6.607** | **1.000** | **13.000** | **2.308** | **0.349** | **4.541** | **0.064** | **0.010** | **0.126** |
| Raptors | Shorebirds | 1.843 | 1.761 | 0.021 | 3.381 | 48.462 | 0.890 | 10.215 | 1.000 | 20.000 | 3.568 | 0.349 | 6.986 | 0.099 | 0.010 | 0.194 |
| Galliformes | Raptors | 1.767 | 1.585 | 0.195 | 3.549 | 35.647 | 0.856 | 6.841 | 1.000 | 14.000 | 2.390 | 0.349 | 4.890 | 0.066 | 0.010 | 0.136 |
| Strigiformes | Shorebirds | 1.719 | 1.350 | 0.082 | 4.279 | 10.465 | 0.636 | 7.913 | 1.000 | 16.000 | 2.764 | 0.349 | 5.589 | 0.077 | 0.010 | 0.155 |
| Shorebirds | Raptors | 1.676 | 1.553 | 0.358 | 3.249 | 68.526 | 0.919 | 12.752 | 4.000 | 23.000 | 4.454 | 1.397 | 8.034 | 0.123 | 0.039 | 0.223 |
| Strigiformes | Passeriformes | 1.645 | 1.491 | 0.246 | 3.428 | 72.667 | 0.924 | 6.307 | 1.000 | 13.000 | 2.203 | 0.349 | 4.541 | 0.061 | 0.010 | 0.126 |
| Raptors | Strigiformes | 1.630 | 1.400 | 0.227 | 3.355 | 34.457 | 0.852 | 12.966 | 2.000 | 21.000 | 4.529 | 0.699 | 7.335 | 0.126 | 0.019 | 0.203 |
| Shorebirds | Anseriformes | 1.609 | 1.390 | 0.188 | 3.173 | 88.400 | 0.936 | 12.370 | 4.000 | 23.000 | 4.321 | 1.397 | 8.034 | 0.120 | 0.039 | 0.223 |
| Nonhuman - mammal | Anseriformes | 1.578 | 1.257 | 0.070 | 4.015 | 9.733 | 0.619 | 6.065 | 1.000 | 13.000 | 2.118 | 0.349 | 4.541 | 0.059 | 0.010 | 0.126 |
| Strigiformes | Galliformes | 1.569 | 1.155 | 0.032 | 4.223 | 3.077 | 0.339 | 4.346 | 1.000 | 10.000 | 1.518 | 0.349 | 3.493 | 0.042 | 0.010 | 0.097 |
| Raptors | Nonhuman-mammal | 1.563 | 1.217 | 0.043 | 3.509 | 18.414 | 0.754 | 5.693 | 1.000 | 12.000 | 1.989 | 0.349 | 4.192 | 0.055 | 0.010 | 0.116 |

| | | | | | | | | | | | | | | |
|---|---|---|---|---|---|---|---|---|---|---|---|---|---|---|
| Nonhuman - mammal | Shorebirds | 1.550 | 1.199 | 0.099 | 3.939 | 6.991 | 0.538 | 4.844 | 1.000 | 11.000 | 1.692 | 0.349 | 3.842 | 0.047 | 0.010 | 0.107 |
| **Shorebirds** | **Passeriformes** | **1.467** | **1.377** | **0.432** | **2.889** | **196.286** | **0.970** | **10.579** | **5.000** | **17.000** | **3.695** | **1.747** | **5.938** | **0.102** | **0.048** | **0.165** |
| Nonhuman - mammal | Raptors | 1.450 | 1.293 | 0.011 | 3.363 | 20.222 | 0.771 | 7.361 | 1.000 | 14.000 | 2.571 | 0.349 | 4.890 | 0.071 | 0.010 | 0.136 |
| Galliformes | Shorebirds | 1.448 | 1.241 | 0.009 | 3.263 | 25.467 | 0.809 | 4.992 | 1.000 | 11.000 | 1.744 | 0.349 | 3.842 | 0.048 | 0.010 | 0.107 |
| Strigiformes | Nonhuman-mammal | 1.398 | 0.998 | 0.005 | 3.770 | 7.234 | 0.547 | 3.981 | 1.000 | 9.000 | 1.391 | 0.349 | 3.144 | 0.039 | 0.010 | 0.087 |
| Raptors | Galliformes | 1.393 | 0.950 | 0.014 | 3.926 | 3.316 | 0.356 | 3.613 | 1.000 | 9.000 | 1.262 | 0.349 | 3.144 | 0.035 | 0.010 | 0.087 |
| Shorebirds | Strigiformes | 1.370 | 1.019 | 0.003 | 4.025 | 9.226 | 0.606 | 5.959 | 1.000 | 12.000 | 2.082 | 0.349 | 4.192 | 0.058 | 0.010 | 0.116 |
| Nonhuman - mammal | Strigiformes | 1.350 | 1.024 | 0.023 | 3.472 | 9.391 | 0.610 | 5.693 | 1.000 | 13.000 | 1.989 | 0.349 | 4.541 | 0.055 | 0.010 | 0.126 |
| Anseriformes | Passeriformes | 1.304 | 0.970 | 0.100 | 3.500 | 7.358 | 0.551 | 24.674 | 11.000 | 35.000 | 8.619 | 3.842 | 12.226 | 0.239 | 0.107 | 0.339 |

**Table S6:** Results of BSSVS discrete trait analysis for Taxonomic host order (equal order sample 1). Table is in order of highest mean transition rate to lowest with rates having BF > 100 bolded. Rates with at least a BF > 3 are shown.

| From | To | mean | median | l95hpd | h95hpd | BAYES FACTOR | POSTERIOR PROBABILITY | mean Markov Jumps | Markov Jumps 95% lhpd | Markov Jumps 95% hhpd | mean Markov Jump / tree height | Markov jumps / tree hieght lower 95%hpd | Markov jumps / tree hieght upper 95%hpd | Markov Jump / tree length | Markov Jump / tree length lower 95% hpd | Markov Jump / tree length upper 95% hpd |
|---|---|---|---|---|---|---|---|---|---|---|---|---|---|---|---|---|
| **Anseriformes** | **Galliformes** | **4.451** | **4.350** | **2.028** | **7.004** | **5400.000** | **0.999** | **49.279** | **30.000** | **67.000** | **15.519** | **9.448** | **21.100** | **0.523** | **0.319** | **0.712** |
| **Anseriformes** | **Strigiformes** | **2.987** | **2.886** | **1.293** | **5.258** | **194.222** | **0.970** | **33.088** | **15.000** | **49.000** | **10.420** | **4.724** | **15.431** | **0.351** | **0.159** | **0.520** |
| **Galliformes** | **Strigiformes** | **2.791** | **2.672** | **0.936** | **4.954** | **194.222** | **0.970** | **28.210** | **10.000** | **44.000** | **8.884** | **3.149** | **13.856** | **0.300** | **0.106** | **0.467** |
| **Galliformes** | **Nonhuman-mammal** | **2.712** | **2.620** | **1.173** | **4.723** | **409.846** | **0.986** | **26.481** | **13.000** | **37.000** | **8.339** | **4.094** | **11.652** | **0.281** | **0.138** | **0.393** |
| **Galliformes** | **Raptors** | **2.455** | **2.288** | **0.932** | **4.453** | **194.222** | **0.970** | **24.571** | **9.000** | **39.000** | **7.738** | **2.834** | **12.282** | **0.261** | **0.096** | **0.414** |
| **Nonhuman - mammal** | **Anseriformes** | **2.138** | **1.968** | **0.463** | **4.173** | **129.150** | **0.956** | **15.442** | **3.000** | **27.000** | **4.863** | **0.945** | **8.503** | **0.164** | **0.032** | **0.287** |
| **Galliformes** | **Passeriformes** | **2.137** | **2.046** | **0.716** | **3.505** | **239.727** | **0.976** | **20.990** | **10.000** | **30.000** | **6.610** | **3.149** | **9.448** | **0.223** | **0.106** | **0.319** |
| **Raptors** | **Nonhuman-mammal** | **2.106** | **2.009** | **0.333** | **3.880** | **174.200** | **0.967** | **17.918** | **5.000** | **29.000** | **5.643** | **1.575** | **9.133** | **0.190** | **0.053** | **0.308** |
| **Shorebirds** | **Nonhuman-mammal** | **2.001** | **1.900** | **0.740** | **3.467** | **534.600** | **0.989** | **17.302** | **9.000** | **24.000** | **5.449** | **2.834** | **7.558** | **0.184** | **0.096** | **0.255** |
| Galliformes | Anseriformes | 1.892 | 1.758 | 0.495 | 3.653 | 96.000 | 0.941 | 17.963 | 4.000 | 30.000 | 5.657 | 1.260 | 9.448 | 0.191 | 0.042 | 0.319 |
| **Shorebirds** | **Raptors** | **1.863** | **1.783** | **0.522** | **3.374** | **444.500** | **0.987** | **15.574** | **7.000** | **25.000** | **4.905** | **2.204** | **7.873** | **0.165** | **0.074** | **0.266** |
| Anseriformes | Raptors | 1.855 | 1.739 | 0.383 | 3.679 | 72.348 | 0.923 | 19.340 | 5.000 | 34.000 | 6.091 | 1.575 | 10.707 | 0.205 | 0.053 | 0.361 |
| **Raptors** | **Strigiformes** | **1.851** | **1.742** | **0.337** | **3.512** | **153.000** | **0.962** | **15.209** | **4.000** | **26.000** | **4.790** | **1.260** | **8.188** | **0.162** | **0.042** | **0.276** |
| Raptors | Anseriformes | 1.846 | 1.704 | 0.220 | 3.615 | 69.083 | 0.920 | 15.037 | 2.000 | 26.000 | 4.735 | 0.630 | 8.188 | 0.160 | 0.021 | 0.276 |
| **Strigiformes** | **Raptors** | **1.790** | **1.643** | **0.235** | **3.447** | **132.615** | **0.957** | **12.926** | **3.000** | **23.000** | **4.070** | **0.945** | **7.243** | **0.137** | **0.032** | **0.244** |
| Raptors | Shorebirds | 1.688 | 1.566 | 0.260 | 3.486 | 64.208 | 0.915 | 14.188 | 3.000 | 27.000 | 4.468 | 0.945 | 8.503 | 0.151 | 0.032 | 0.287 |
| **Strigiformes** | **Passeriformes** | **1.595** | **1.470** | **0.330** | **3.120** | **109.021** | **0.948** | **12.416** | **2.000** | **21.000** | **3.910** | **0.630** | **6.613** | **0.132** | **0.021** | **0.223** |
| **Anseriformes** | **Shorebirds** | **1.566** | **1.455** | **0.342** | **3.037** | **111.522** | **0.949** | **16.447** | **5.000** | **27.000** | **5.179** | **1.575** | **8.503** | **0.175** | **0.053** | **0.287** |
| Nonhuman - mammal | Galliformes | 1.430 | 1.142 | 0.009 | 3.580 | 6.121 | 0.505 | 7.415 | 1.000 | 16.000 | 2.335 | 0.315 | 5.039 | 0.079 | 0.011 | 0.170 |
| Passeriformes | Raptors | 1.421 | 1.141 | 0.006 | 3.501 | 5.803 | 0.492 | 6.209 | 1.000 | 12.000 | 1.955 | 0.315 | 3.779 | 0.066 | 0.011 | 0.127 |
| Strigiformes | Galliformes | 1.372 | 1.038 | 0.014 | 3.833 | 3.636 | 0.377 | 7.221 | 1.000 | 17.000 | 2.274 | 0.315 | 5.354 | 0.077 | 0.011 | 0.181 |
| Raptors | Passeriformes | 1.347 | 1.040 | 0.010 | 3.471 | 7.250 | 0.547 | 8.488 | 1.000 | 17.000 | 2.673 | 0.315 | 5.354 | 0.090 | 0.011 | 0.181 |
| Nonhuman - mammal | Raptors | 1.322 | 1.044 | 0.004 | 3.295 | 5.310 | 0.469 | 6.728 | 1.000 | 15.000 | 2.119 | 0.315 | 4.724 | 0.071 | 0.011 | 0.159 |
| Shorebirds | Strigiformes | 1.320 | 0.793 | 0.003 | 4.003 | 3.086 | 0.340 | 4.692 | 1.000 | 10.000 | 1.478 | 0.315 | 3.149 | 0.050 | 0.011 | 0.106 |
| Galliformes | Shorebirds | 1.317 | 1.175 | 0.132 | 2.690 | 30.527 | 0.836 | 11.262 | 1.000 | 20.000 | 3.547 | 0.315 | 6.298 | 0.120 | 0.011 | 0.212 |

| | | | | | | | | | | | | | | | | |
|---|---|---|---|---|---|---|---|---|---|---|---|---|---|---|---|---|
| Strigiformes | Shorebirds | 1.311 | 1.074 | 0.032 | 3.117 | 11.327 | 0.654 | 8.375 | 1.000 | 16.000 | 2.637 | 0.315 | 5.039 | 0.089 | 0.011 | 0.170 |
| Nonhuman - mammal | Strigiformes | 1.303 | 0.949 | 0.002 | 3.716 | 5.576 | 0.482 | 6.347 | 1.000 | 14.000 | 1.999 | 0.315 | 4.409 | 0.067 | 0.011 | 0.149 |
| Nonhuman - mammal | Shorebirds | 1.291 | 0.974 | 0.011 | 3.753 | 6.013 | 0.501 | 5.966 | 1.000 | 12.000 | 1.879 | 0.315 | 3.779 | 0.063 | 0.011 | 0.127 |
| Passeriformes | Anseriformes | 1.255 | 1.026 | 0.039 | 2.911 | 29.801 | 0.832 | 5.402 | 1.000 | 10.000 | 1.701 | 0.315 | 3.149 | 0.057 | 0.011 | 0.106 |
| Raptors | Galliformes | 1.240 | 0.988 | 0.002 | 3.038 | 11.495 | 0.657 | 8.312 | 1.000 | 16.000 | 2.618 | 0.315 | 5.039 | 0.088 | 0.011 | 0.170 |
| Strigiformes | Nonhuman-mammal | 1.239 | 0.899 | 0.006 | 3.383 | 6.810 | 0.532 | 6.466 | 1.000 | 14.000 | 2.036 | 0.315 | 4.409 | 0.069 | 0.011 | 0.149 |
| **Shorebirds** | **Passeriformes** | **1.152** | **1.023** | **0.222** | **2.282** | **140.108** | **0.959** | **9.277** | **3.000** | **14.000** | **2.922** | **0.945** | **4.409** | **0.099** | **0.032** | **0.149** |
| Shorebirds | Anseriformes | 1.118 | 0.873 | 0.002 | 3.217 | 11.439 | 0.656 | 6.597 | 1.000 | 13.000 | 2.077 | 0.315 | 4.094 | 0.070 | 0.011 | 0.138 |

**Table S7:** Results of BSSVS discrete trait analysis for Taxonomic host order (equal order sample 2). Table is in order of highest mean transition rate to lowest with rates having BF > 100 bolded. Rates with at least a BF > 3 are shown.

| From | To | mean | median | l95hpd | h95hpd | BAYES FACTOR | POSTERIOR PROBABILITY | mean Markov Jumps | Markov Jumps 95% lhpd | Markov Jumps 95% hhpd | mean Markov Jump / tree height | Markov jumps / tree hieght lower 95%hpd | Markov jumps / tree hieght upper 95%hpd | Markov Jump / tree length | Markov Jump / tree length lower 95% hpd | Markov Jump / tree length upper 95% hpd |
|---|---|---|---|---|---|---|---|---|---|---|---|---|---|---|---|---|
| **Anseriformes** | **Galliformes** | **4.593** | **4.473** | **1.842** | **7.736** | **1075.2** | **0.994** | **48.298** | **24** | **72** | **20.316** | **10.095** | **30.286** | **0.454** | **0.225** | **0.676** |
| **Anseriformes** | **Strigiformes** | **3.674** | **3.455** | **1.467** | **6.572** | **210.24** | **0.972** | **39.473** | **17** | **61** | **16.604** | **7.151** | **25.659** | **0.371** | **0.16** | **0.573** |
| **Galliformes** | **Strigiformes** | **3.278** | **3.148** | **0.974** | **5.738** | **219.25** | **0.973** | **30.129** | **8** | **49** | **12.674** | **3.365** | **20.611** | **0.283** | **0.075** | **0.46** |
| **Anseriformes** | **Raptors** | **3.171** | **3.014** | **1.071** | **5.517** | **312** | **0.981** | **33.503** | **14** | **53** | **14.093** | **5.889** | **22.294** | **0.315** | **0.132** | **0.498** |
| **Galliformes** | **Anseriformes** | **2.974** | **2.821** | **0.685** | **5.363** | **201.923** | **0.971** | **27.024** | **6** | **48** | **11.368** | **2.524** | **20.191** | **0.254** | **0.056** | **0.451** |
| **Galliformes** | **Nonhuman-mammal** | **2.682** | **2.533** | **0.617** | **4.865** | **116.864** | **0.951** | **24.272** | **7** | **39** | **10.21** | **2.944** | **16.405** | **0.228** | **0.066** | **0.366** |
| Nonhuman - mammal | Anseriformes | 2.505 | 2.361 | 0.304 | 4.87 | 64.208 | 0.915 | 17.048 | 3 | 32 | 7.171 | 1.262 | 13.461 | 0.16 | 0.028 | 0.301 |
| Strigiformes | Raptors | 2.045 | 1.884 | 0.246 | 4.204 | 48.06 | 0.889 | 14.44 | 3 | 26 | 6.074 | 1.262 | 10.937 | 0.136 | 0.028 | 0.244 |
| **Shorebirds** | **Nonhuman-mammal** | **2.03** | **1.915** | **0.733** | **3.622** | **766.286** | **0.992** | **16.093** | **8** | **23** | **6.769** | **3.365** | **9.675** | **0.151** | **0.075** | **0.216** |
| **Anseriformes** | **Shorebirds** | **1.917** | **1.799** | **0.551** | **3.74** | **264.3** | **0.978** | **19.757** | **6** | **32** | **8.311** | **2.524** | **13.461** | **0.186** | **0.056** | **0.301** |
| Galliformes | Passeriformes | 1.865 | 1.766 | 0.165 | 3.479 | 81.194 | 0.931 | 16.82 | 3 | 28 | 7.075 | 1.262 | 11.778 | 0.158 | 0.028 | 0.263 |
| Raptors | Anseriformes | 1.821 | 1.63 | 0.012 | 3.84 | 23.867 | 0.799 | 13.928 | 1 | 26 | 5.859 | 0.421 | 10.937 | 0.131 | 0.009 | 0.244 |
| **Shorebirds** | **Raptors** | **1.74** | **1.615** | **0.329** | **3.322** | **380.143** | **0.984** | **13.668** | **4** | **22** | **5.749** | **1.683** | **9.254** | **0.128** | **0.038** | **0.207** |
| Anseriformes | Nonhuman-mammal | 1.664 | 1.508 | 0.096 | 3.539 | 30.282 | 0.835 | 16.275 | 1 | 32 | 6.846 | 0.421 | 13.461 | 0.153 | 0.009 | 0.301 |
| Galliformes | Shorebirds | 1.65 | 1.488 | 0.215 | 3.383 | 41.009 | 0.872 | 13.486 | 2 | 24 | 5.673 | 0.841 | 10.095 | 0.127 | 0.019 | 0.225 |
| Raptors | Shorebirds | 1.63 | 1.499 | 0.172 | 3.494 | 36.905 | 0.86 | 12.477 | 2 | 22 | 5.248 | 0.841 | 9.254 | 0.117 | 0.019 | 0.207 |
| Galliformes | Raptors | 1.624 | 1.416 | 0.039 | 3.523 | 21.303 | 0.78 | 13.357 | 2 | 26 | 5.619 | 0.841 | 10.937 | 0.125 | 0.019 | 0.244 |
| Strigiformes | Shorebirds | 1.602 | 1.328 | 0.016 | 3.97 | 12.706 | 0.679 | 9.599 | 1 | 18 | 4.038 | 0.421 | 7.572 | 0.09 | 0.009 | 0.169 |
| Raptors | Strigiformes | 1.588 | 1.406 | 0.152 | 3.424 | 31.542 | 0.84 | 11.929 | 2 | 23 | 5.018 | 0.841 | 9.675 | 0.112 | 0.019 | 0.216 |
| Nonhuman - mammal | Galliformes | 1.574 | 1.237 | 0.031 | 4.031 | 9.101 | 0.603 | 8.156 | 1 | 17 | 3.431 | 0.421 | 7.151 | 0.077 | 0.009 | 0.16 |
| Raptors | Nonhuman-mammal | 1.548 | 1.387 | 0.296 | 3.337 | 77.169 | 0.928 | 11.618 | 2 | 20 | 4.887 | 0.841 | 8.413 | 0.109 | 0.019 | 0.188 |
| Strigiformes | Passeriformes | 1.535 | 1.425 | 0.183 | 3.086 | 66.08 | 0.917 | 10.874 | 2 | 19 | 4.574 | 0.841 | 7.992 | 0.102 | 0.019 | 0.178 |
| Passeriformes | Strigiformes | 1.473 | 1.095 | 0.004 | 4.282 | 3.758 | 0.385 | 4.923 | 1 | 11 | 2.071 | 0.421 | 4.627 | 0.046 | 0.009 | 0.103 |
| Strigiformes | Galliformes | 1.46 | 1.19 | 0.005 | 3.587 | 8.975 | 0.599 | 8.907 | 1 | 19 | 3.747 | 0.421 | 7.992 | 0.084 | 0.009 | 0.178 |
| Passeriformes | Raptors | 1.454 | 1.137 | 0.026 | 3.939 | 7.686 | 0.562 | 4.365 | 1 | 9 | 1.836 | 0.421 | 3.786 | 0.041 | 0.009 | 0.085 |
| Nonhuman - mammal | Strigiformes | 1.442 | 1.137 | 0.007 | 3.951 | 5.429 | 0.475 | 6.615 | 1 | 14 | 2.782 | 0.421 | 5.889 | 0.062 | 0.009 | 0.132 |
| Nonhuman - mammal | Raptors | 1.424 | 1.054 | 0.024 | 3.917 | 5.381 | 0.473 | 6.571 | 1 | 14 | 2.764 | 0.421 | 5.889 | 0.062 | 0.009 | 0.132 |

| | | | | | | | | | | | | | | | | |
|---|---|---|---|---|---|---|---|---|---|---|---|---|---|---|---|---|
| Anseriformes | Passeriformes | 1.413 | 1.186 | 0.015 | 3.276 | 18.912 | 0.759 | 12.582 | 1 | 25 | 5.293 | 0.421 | 10.516 | 0.118 | 0.009 | 0.235 |
| Nonhuman - mammal | Shorebirds | 1.399 | 1.077 | 0.009 | 3.757 | 5.676 | 0.486 | 6.42 | 1 | 15 | 2.701 | 0.421 | 6.31 | 0.06 | 0.009 | 0.141 |
| Strigiformes | Nonhuman-mammal | 1.382 | 1.165 | 0.008 | 3.225 | 20.116 | 0.77 | 7.754 | 1 | 15 | 3.262 | 0.421 | 6.31 | 0.073 | 0.009 | 0.141 |
| Raptors | Galliformes | 1.367 | 1.114 | 0.005 | 3.302 | 10.634 | 0.639 | 8.329 | 1 | 17 | 3.504 | 0.421 | 7.151 | 0.078 | 0.009 | 0.16 |
| Shorebirds | Strigiformes | 1.227 | 0.806 | 0.004 | 3.752 | 5.881 | 0.495 | 4.89 | 1 | 10 | 2.057 | 0.421 | 4.206 | 0.046 | 0.009 | 0.094 |
| **Shorebirds** | **Passeriformes** | **1.22** | **1.147** | **0.221** | **2.328** | **162.938** | **0.964** | **9.431** | **4** | **14** | **3.967** | **1.683** | **5.889** | **0.089** | **0.038** | **0.132** |
| Shorebirds | Anseriformes | 1.196 | 0.933 | 0.013 | 2.953 | 20.762 | 0.776 | 7.195 | 1 | 14 | 3.026 | 0.421 | 5.889 | 0.068 | 0.009 | 0.132 |

**Table S8:** Results of BSSVS discrete trait analysis for Taxonomic host order (equal order sample 3). Table is in order of highest mean transition rate to lowest with rates having BF > 100 bolded. Rates with at least a BF > 3 are shown.

| From | To | mean | median | l95hpd | h95hpd | BAYES FACTOR | POSTERIOR PROBABILITY | mean Markov Jumps | Markov Jumps 95% lhpd | Markov Jumps 95% hhpd | mean Markov Jump / tree height | Markov jumps / tree hieght lower 95%hpd | Markov jumps / tree hieght upper 95%hpd | Markov Jump / tree length | Markov Jump / tree length lower 95% hpd | Markov Jump / tree length upper 95% hpd |
|---|---|---|---|---|---|---|---|---|---|---|---|---|---|---|---|---|
| **Anseriformes** | **Raptors** | **5.127** | **5.039** | **2.671** | **7.870** | **6000.000** | **1.000** | **85.678** | **72.000** | **99.000** | **44.299** | **37.227** | **51.187** | **0.647** | **0.543** | **0.747** |
| **Anseriformes** | **Galliformes** | **4.187** | **4.061** | **2.120** | **6.373** | **6000.000** | **1.000** | **47.280** | **36.000** | **59.000** | **24.445** | **18.613** | **30.505** | **0.357** | **0.272** | **0.445** |
| **Raptors** | **Anseriformes** | **2.938** | **2.736** | **0.880** | **5.486** | **347.294** | **0.983** | **27.786** | **11.000** | **43.000** | **14.366** | **5.687** | **22.233** | **0.210** | **0.083** | **0.324** |
| Raptors | Shorebirds | 1.710 | 1.581 | 0.029 | 3.281 | 42.435 | 0.876 | 15.135 | 7.000 | 23.000 | 7.825 | 3.619 | 11.892 | 0.114 | 0.053 | 0.174 |
| **Anseriformes** | **Strigiformes** | **1.456** | **1.405** | **0.606** | **2.306** | **6000.000** | **1.000** | **31.811** | **22.000** | **40.000** | **16.448** | **11.375** | **20.681** | **0.240** | **0.166** | **0.302** |
| Galliformes | Anseriformes | 1.302 | 1.035 | 0.002 | 3.313 | 7.903 | 0.568 | 8.462 | 1.000 | 19.000 | 4.375 | 0.517 | 9.824 | 0.064 | 0.008 | 0.143 |
| Galliformes | Raptors | 1.178 | 0.908 | 0.017 | 3.183 | 4.391 | 0.423 | 9.857 | 1.000 | 18.000 | 5.096 | 0.517 | 9.307 | 0.074 | 0.008 | 0.136 |
| **Anseriformes** | **Shorebirds** | **1.082** | **0.989** | **0.288** | **2.083** | **456.000** | **0.987** | **20.930** | **11.000** | **29.000** | **10.822** | **5.687** | **14.994** | **0.158** | **0.083** | **0.219** |
| Shorebirds | Raptors | 1.067 | 0.797 | 0.003 | 3.008 | 4.128 | 0.408 | 6.214 | 1.000 | 13.000 | 3.213 | 0.517 | 6.721 | 0.047 | 0.008 | 0.098 |
| Shorebirds | Anseriformes | 1.063 | 0.762 | 0.005 | 3.049 | 4.611 | 0.435 | 10.599 | 4.000 | 17.000 | 5.480 | 2.068 | 8.790 | 0.080 | 0.030 | 0.128 |
| Raptors | Nonhuman-Mammal | 1.016 | 0.677 | 0.007 | 3.116 | 3.766 | 0.386 | 7.885 | 2.000 | 13.000 | 4.077 | 1.034 | 6.721 | 0.059 | 0.015 | 0.098 |
| Galliformes | Nonhuman-Mammal | 0.983 | 0.765 | 0.003 | 2.571 | 10.410 | 0.634 | 3.114 | 1.000 | 6.000 | 1.610 | 0.517 | 3.102 | 0.024 | 0.008 | 0.045 |
| **Anseriformes** | **Nonhuman-Mammal** | **0.710** | **0.658** | **0.183** | **1.277** | **255.130** | **0.977** | **7.070** | **1.000** | **11.000** | **3.655** | **0.517** | **5.687** | **0.053** | **0.008** | **0.083** |
| Raptors | Passeriformes | 0.664 | 0.550 | 0.040 | 1.454 | 47.150 | 0.887 | 8.598 | 2.000 | 15.000 | 4.445 | 1.034 | 7.756 | 0.065 | 0.015 | 0.113 |
| **Anseriformes** | **Passeriformes** | **0.569** | **0.530** | **0.138** | **1.024** | **1195.200** | **0.995** | **13.836** | **7.000** | **22.000** | **7.154** | **3.619** | **11.375** | **0.104** | **0.053** | **0.166** |

**Table S9:** Results of BSSVS discrete trait analysis for Taxonomic host order (case proportional sample 1). Table is in order of highest mean transition rate to lowest with rates having BF > 100 bolded. Rates with at least a BF > 3 are shown.

| From | To | mean | median | l95hpd | h95hpd | BAYES FACTOR | POSTERIOR PROBABILITY | mean Markov Jumps | Markov Jumps 95% lhpd | Markov Jumps 95% hhpd | mean Markov Jump / tree height | Markov jumps / tree hieght lower 95%hpd | Markov jumps / tree hieght upper 95%hpd | Markov Jump / tree length | Markov Jump / tree length lower 95% hpd | Markov Jump / tree length upper 95% hpd |
|---|---|---|---|---|---|---|---|---|---|---|---|---|---|---|---|---|
| **Anseriformes** | **Raptors** | **6.506** | **6.299** | **3.435** | **9.973** | **6000.000** | **1.000** | **87.856** | **75.000** | **99.000** | **34.776** | **29.687** | **39.187** | **0.603** | **0.515** | **0.680** |
| **Anseriformes** | **Galliformes** | **4.443** | **4.280** | **2.520** | **7.024** | **6000.000** | **1.000** | **42.613** | **30.000** | **52.000** | **16.867** | **11.875** | **20.583** | **0.293** | **0.206** | **0.357** |
| **Raptors** | **Anseriformes** | **3.009** | **2.830** | **0.986** | **5.906** | **208.500** | **0.972** | **34.512** | **14.000** | **54.000** | **13.661** | **5.542** | **21.375** | **0.237** | **0.096** | **0.371** |
| **Anseriformes** | **Shorebirds** | **2.133** | **2.023** | **0.931** | **3.447** | **6000.000** | **1.000** | **29.608** | **14.000** | **43.000** | **11.720** | **5.542** | **17.020** | **0.203** | **0.096** | **0.295** |
| Galliformes | Anseriformes | 1.696 | 1.451 | 0.050 | 4.017 | 16.664 | 0.735 | 7.348 | 1.000 | 17.000 | 2.908 | 0.396 | 6.729 | 0.050 | 0.007 | 0.117 |
| **Anseriformes** | **Strigiformes** | **1.528** | **1.437** | **0.533** | **2.554** | **6000.000** | **0.999** | **27.804** | **17.000** | **38.000** | **11.006** | **6.729** | **15.041** | **0.191** | **0.117** | **0.261** |
| Galliformes | Raptors | 1.358 | 1.036 | 0.035 | 3.744 | 3.734 | 0.384 | 6.912 | 1.000 | 13.000 | 2.736 | 0.396 | 5.146 | 0.047 | 0.007 | 0.089 |
| Shorebirds | Anseriformes | 1.357 | 1.048 | 0.005 | 3.559 | 5.708 | 0.488 | 8.537 | 1.000 | 18.000 | 3.379 | 0.396 | 7.125 | 0.059 | 0.007 | 0.124 |
| Raptors | Strigiformes | 1.355 | 1.173 | 0.114 | 3.012 | 32.500 | 0.844 | 17.417 | 6.000 | 28.000 | 6.894 | 2.375 | 11.083 | 0.120 | 0.041 | 0.192 |
| Raptors | Nonhuman-Mammal | 1.329 | 0.942 | 0.003 | 3.853 | 3.032 | 0.336 | 4.321 | 1.000 | 10.000 | 1.710 | 0.396 | 3.958 | 0.030 | 0.007 | 0.069 |
| Raptors | Shorebirds | 1.278 | 0.990 | 0.006 | 3.764 | 3.326 | 0.357 | 21.252 | 7.000 | 36.000 | 8.412 | 2.771 | 14.250 | 0.146 | 0.048 | 0.247 |
| Galliformes | Nonhuman-Mammal | 1.237 | 0.922 | 0.007 | 3.621 | 3.059 | 0.338 | 2.925 | 1.000 | 6.000 | 1.158 | 0.396 | 2.375 | 0.020 | 0.007 | 0.041 |
| Strigiformes | Raptors | 1.203 | 0.862 | 0.001 | 3.389 | 4.145 | 0.409 | 12.006 | 1.000 | 22.000 | 4.752 | 0.396 | 8.708 | 0.082 | 0.007 | 0.151 |
| Shorebirds | Nonhuman-Mammal | 1.140 | 0.872 | 0.003 | 2.965 | 6.307 | 0.512 | 3.665 | 1.000 | 6.000 | 1.451 | 0.396 | 2.375 | 0.025 | 0.007 | 0.041 |
| Passeriformes | Galliformes | 1.118 | 0.793 | 0.003 | 3.318 | 5.440 | 0.476 | 3.573 | 1.000 | 7.000 | 1.414 | 0.396 | 2.771 | 0.025 | 0.007 | 0.048 |
| **Anseriformes** | **Nonhuman-Mammal** | **0.995** | **0.921** | **0.198** | **1.709** | **181.688** | **0.968** | **11.096** | **6.000** | **15.000** | **4.392** | **2.375** | **5.937** | **0.076** | **0.041** | **0.103** |
| **Anseriformes** | **Passeriformes** | **0.740** | **0.694** | **0.220** | **1.367** | **394.400** | **0.985** | **15.569** | **8.000** | **23.000** | **6.163** | **3.167** | **9.104** | **0.107** | **0.055** | **0.158** |

**Table S10:** Results of BSSVS discrete trait analysis for Taxonomic host order (case proportional sample 2). Table is in order of highest mean transition rate to lowest with rates having BF > 100 bolded. Rates with at least a BF > 3 are shown.

| From | To | mean | median | l95hpd | h95hpd | BAYES FACTOR | POSTERIOR PROBABILITY | mean Markov Jumps | Markov Jumps 95% lhpd | Markov Jumps 95% hhpd | mean Markov Jump / tree height | Markov jumps / tree hieght lower 95%hpd | Markov jumps / tree hieght upper 95%hpd | Markov Jump / tree length | Markov Jump / tree length lower 95% hpd | Markov Jump / tree length upper 95% hpd |
|---|---|---|---|---|---|---|---|---|---|---|---|---|---|---|---|---|
| **Anseriformes** | **Raptors** | **6.016** | **5.933** | **3.202** | **9.178** | **6000.000** | **1.000** | **83.491** | **69.000** | **99.000** | **36.763** | **30.382** | **43.592** | **0.612** | **0.506** | **0.726** |
| **Anseriformes** | **Galliformes** | **4.648** | **4.525** | **2.393** | **7.064** | **6000.000** | **1.000** | **53.319** | **42.000** | **61.000** | **23.478** | **18.494** | **26.860** | **0.391** | **0.308** | **0.447** |
| **Raptors** | **Anseriformes** | **2.904** | **2.684** | **0.915** | **5.356** | **423.000** | **0.986** | **16.650** | **3.000** | **31.000** | **7.331** | **1.321** | **13.650** | **0.122** | **0.022** | **0.227** |
| **Anseriformes** | **Shorebirds** | **1.724** | **1.632** | **0.566** | **2.983** | **6000.000** | **1.000** | **25.697** | **13.000** | **37.000** | **11.315** | **5.724** | **16.292** | **0.188** | **0.095** | **0.271** |
| **Anseriformes** | **Strigiformes** | **1.540** | **1.472** | **0.659** | **2.660** | **6000.000** | **1.000** | **21.595** | **12.000** | **29.000** | **9.509** | **5.284** | **12.769** | **0.158** | **0.088** | **0.213** |
| Raptors | Shorebirds | 1.376 | 1.197 | 0.084 | 3.069 | 15.761 | 0.724 | 18.455 | 8.000 | 28.000 | 8.126 | 3.523 | 12.329 | 0.135 | 0.059 | 0.205 |
| Shorebirds | Anseriformes | 1.181 | 0.988 | 0.043 | 2.996 | 64.659 | 0.915 | 15.695 | 5.000 | 25.000 | 6.911 | 2.202 | 11.008 | 0.115 | 0.037 | 0.183 |
| **Raptors** | **Strigiformes** | **1.155** | **1.012** | **0.053** | **2.409** | **116.571** | **0.951** | **20.145** | **11.000** | **29.000** | **8.870** | **4.844** | **12.769** | **0.148** | **0.081** | **0.213** |
| **Anseriformes** | **Nonhuman-Mammal** | **1.029** | **0.986** | **0.413** | **1.852** | **6000.000** | **1.000** | **18.171** | **9.000** | **27.000** | **8.001** | **3.963** | **11.889** | **0.133** | **0.066** | **0.198** |
| Passeriformes | Raptors | 0.961 | 0.720 | 0.014 | 2.696 | 11.014 | 0.647 | 4.772 | 1.000 | 10.000 | 2.101 | 0.440 | 4.403 | 0.035 | 0.007 | 0.073 |
| **Anseriformes** | **Passeriformes** | **0.731** | **0.696** | **0.186** | **1.341** | **1195.200** | **0.995** | **12.717** | **5.000** | **20.000** | **5.599** | **2.202** | **8.806** | **0.093** | **0.037** | **0.147** |

**Table S11:** Results of BSSVS discrete trait analysis for Taxonomic host order (case proportional sample 3). Table is in order of highest mean transition rate to lowest with rates having BF > 100 bolded. Rates with at least a BF > 3 are shown.

| Analysis | ratio | Transitions to Domestic | Transitions to Wild |
|---|---|---|---|
| 2-state | 1:1 | 46 | 40 |
| 2-state | 1:1.5 | 66 | 29 |
| 2-state | 1:2 | 79 | 16 |
| 2-state | 1:2.5 | 99 | 4 |
| 2-state | 1:3 | 106 | 4 |
| 2-state with turkey | 1:1 | 61 | 51 |
| 2-state with turkey | 1:1.5 | 109 | 26 |

| | | | |
|---|---|---|---|
| 2-state with turkey | 1:2 | 113 | 4 |

**Table S12:** Transitions into Domestic and Wild for each ratio dataset for two state rarefaction and two state rarefaction with turkey sequences.

| **From** | **To** | **Transition count** |
|---|---|---|
| Wild | Domestic | 48 |
| Wild | Turkey | 42 |
| Domestic | Wild | 5 |
| Domestic | Turkey | 18 |
| Turkey | Domestic | 38 |
| Turkey | Wild | 1 |

**Table S13**: Number of transitions between states in three state rarefaction of wild, domestic, and turkey for the 1:2 domestic:wild with turkey sequence dataset.

| From | To | mean | median | l95hpd | h95hpd | BAYES_FACTOR | POSTERIOR PROBABILITY |
|---|---|---|---|---|---|---|---|
| **Wild** | **Backyard_bird** | **1.86774162** | **1.70511383** | **0.37454847** | **3.68662003** | **18000** | **1** |
| **Wild** | **Commercial** | **1.46084104** | **1.31711214** | **0.26387335** | **3.00326823** | **18000** | **1** |
| Backyard_bird | Commercial | 0.95115148 | 0.81135468 | 0.07030257 | 2.20310188 | 87.5621891 | 0.97766915 |
| Backyard_bird | Wild | 0.59728953 | 0.41999835 | 0.00047605 | 1.71248402 | 7.84792123 | 0.79691145 |
| Commercial | Backyard_bird | 0.58676046 | 0.29058374 | 3.52E-05 | 2.18585542 | 2.50726089 | 0.55627153 |
| Commercial | Wild | 0.51687069 | 0.26402531 | 0.00010671 | 1.99546369 | 3.66278704 | 0.64681702 |

**Table S14**. Discrete trait rates for backyard bird wild bird titration analysis (25% of all wild bird sequence). Table is in order of highest mean transition rate to lowest with rates having BF > 100 bolded. Rates with at least a BF > 3 are shown.

| From | To | mean | median | l95hpd | h95hpd | BAYES_FACTOR | POSTERIOR PROBABILITY |
|---|---|---|---|---|---|---|---|
| **Wild** | **Backyard_bird** | **1.91959767** | **1.75610576** | **0.39212384** | **3.81857375** | **18000** | **1** |
| **Wild** | **Commercial** | **1.71435411** | **1.5565693** | **0.3141297** | **3.48816005** | **18000** | **1** |
| Backyard_bird | Commercial | 0.72271333 | 0.56016402 | 0.00049205 | 1.90186488 | 12.5529507 | 0.86257083 |
| Commercial | Wild | 0.57838557 | 0.25387721 | 1.88E-05 | 2.25248367 | 1.92799476 | 0.49083435 |
| Backyard_bird | Wild | 0.56892638 | 0.36543149 | 0.00035904 | 1.81943486 | 5.7561396 | 0.74213976 |
| Commercial | Backyard_bird | 0.49228375 | 0.24128113 | 1.86E-05 | 1.8770157 | 3.86575432 | 0.65903788 |

**Table S15** Discrete trait rates for backyard bird wild bird titration analysis (50% of all wild bird sequence). Table is in order of highest mean transition rate to lowest with rates having BF > 100 bolded. Rates with at least a BF > 3 are shown.

| From | To | mean | median | l95hpd | h95hpd | BAYES_FACTOR | POSTERIOR PROBABILITY |
|---|---|---|---|---|---|---|---|
| **Wild** | **Backyard_bird** | **1.84672556** | **1.68062813** | **0.2989798** | **3.81025501** | **18000** | **1** |
| **Wild** | **Commercial** | **1.58010327** | **1.43058414** | **0.24177847** | **3.28214739** | **18000** | **1** |
| Backyard_bird | Wild | 0.81861356 | 0.50058483 | 1.08E-05 | 2.66432008 | 0.72964367 | 0.26730363 |
| Backyard_bird | Commercial | 0.64099347 | 0.50335486 | 0.0056189 | 1.63180822 | 18.9569267 | 0.90456616 |
| Commercial | Wild | 0.60637747 | 0.24737791 | 1.03E-06 | 2.40045423 | 1.71865317 | 0.46217087 |
| Commercial | Backyard_bird | 0.48361884 | 0.20683227 | 9.64E-06 | 1.94383423 | 3.45680509 | 0.63348517 |

**Table S16.** Discrete trait rates for backyard bird wild bird titration analysis (75% of all wild bird sequence). Table is in order of highest mean transition rate to lowest with rates having BF > 100 bolded. Rates with at least a BF > 3 are shown.

| From | To | mean | median | l95hpd | h95hpd | BAYES FACTOR | POSTERIOR PROBABILITY | mean Markov Jumps | Markov Jumps 95% lhpd | Markov Jumps 95% hhpd | mean Markov Jump / tree height | Markov jumps / tree hieght lower 95%hpd | Markov jumps / tree hieght upper 95%hpd | Markov Jump / tree length | Markov Jump / tree length lower 95% hpd | Markov Jump / tree length upper 95% hpd |
|---|---|---|---|---|---|---|---|---|---|---|---|---|---|---|---|---|
| **Wild** | **Backyard_bird** | **1.825** | **1.644** | **0.283** | **3.787** | **18000.000** | **1.000** | **42.669** | **35.000** | **49.000** | **48.963** | **40.163** | **56.228** | **0.755** | **0.619** | **0.867** |
| **Wild** | **Commercial** | **1.601** | **1.441** | **0.233** | **3.356** | **18000.000** | **1.000** | **40.506** | **35.000** | **46.000** | **46.481** | **40.163** | **52.785** | **0.716** | **0.619** | **0.814** |
| Backyard_bird | Wild | 0.859 | 0.545 | 3.260E-05 | 2.727 | 0.424 | 0.175 | 1.748 | 1.000 | 4.000 | 2.006 | 1.148 | 4.590 | 0.031 | 0.018 | 0.071 |
| Backyard_bird | Commercial | 0.639 | 0.405 | 3.090E-06 | 2.053 | 4.584 | 0.696 | 2.334 | 1.000 | 4.000 | 2.678 | 1.148 | 4.590 | 0.041 | 0.018 | 0.071 |
| Commercial | Backyard_bird | 0.563 | 0.229 | 1.880E-06 | 2.317 | 2.137 | 0.517 | 1.257 | 1.000 | 2.000 | 1.442 | 1.148 | 2.295 | 0.022 | 0.018 | 0.035 |
| Commercial | Wild | 0.519 | 0.209 | 1.090E-05 | 2.072 | 2.649 | 0.570 | 1.262 | 1.000 | 2.000 | 1.448 | 1.148 | 2.295 | 0.022 | 0.018 | 0.035 |

**Table S17** Discrete trait rates for backyard bird wild bird titration analysis (100% of all wild bird sequence). Table is in order of highest mean transition rate to lowest with rates having BF > 100 bolded. Rates with at least a BF > 3 are shown.

| **Analysis** | **Statistic** | **observed mean** | **lower 95% CI** | **upper 95% CU** | **null mean** | **lower 95% CI** | **upper 95% CI** | **significance** |
|---|---|---|---|---|---|---|---|---|
| Geographic introduction | AI | 2.221 | 1.585 | 2.946 | 105.122 | 101.926 | 108.689 | 0.000599 |
| | PS | 19.596 | 19.000 | 21.000 | 550.334 | 544.913 | 556.562 | 0.000599 |
| | Europe | 162.900 | 160.000 | 180.000 | 2.921 | 2.486 | 3.426 | 0.000599 |
| | Asia | 82.552 | 80.000 | 80.000 | 2.778 | 2.379 | 3.181 | 0.000599 |
| | North America | 437.691 | 268.000 | 550.000 | 11.443 | 10.113 | 13.497 | 0.000599 |
| Flyway | AI | 10.563 | 9.345 | 11.880 | 78.911 | 76.250 | 82.111 | 0.00199 |
| | PS | 95.375 | 92.000 | 100.000 | 540.419 | 528.531 | 551.669 | 0.00199 |
| | Atlantic flyway | 41.305 | 41.000 | 43.000 | 3.288 | 2.808 | 4.048 | 0.00199 |
| | Mississippi flyway | 26.024 | 18.000 | 38.000 | 3.332 | 2.796 | 4.202 | 0.00199 |
| | central flyway | 18.707 | 11.000 | 23.000 | 3.316 | 2.792 | 4.056 | 0.00199 |
| | pacific flyway | 27.798 | 22.000 | 42.000 | 3.164 | 2.597 | 4.000 | 0.00199 |
| Migration | AI | 59.277 | 55.857 | 62.574 | 80.351 | 78.342 | 82.289 | 0.00399 |
| | PS | 407.488 | 397.000 | 417.000 | 510.881 | 502.818 | 519.850 | 0.00399 |
| | domestic | 8.310 | 8.000 | 10.000 | 3.784 | 3.278 | 4.916 | 0.00399 |
| | migratory | 6.544 | 6.000 | 8.000 | 4.487 | 3.924 | 5.414 | 0.00399 |
| | nonhuman-mammal | 3.232 | 3.000 | 5.000 | 1.667 | 1.324 | 2.030 | 0.00399 |
| | partially migratory | 5.656 | 5.000 | 8.000 | 2.816 | 2.378 | 3.348 | 0.00399 |
| | sedentary | 3.036 | 3.000 | 3.000 | 1.785 | 1.458 | 2.082 | 0.00399 |
| Host Order | AI | 42.505 | 39.799 | 45.411 | 61.018 | 59.291 | 62.755 | 0.00999 |
| | PS | 334.122 | 325.000 | 342.000 | 437.302 | 429.890 | 445.370 | 0.00999 |
| | Galliformes | 6.000 | 6.000 | 6.000 | 2.321 | 2.052 | 3.020 | 0.00999 |

| | | | | | | | | |
|---|---|---|---|---|---|---|---|---|
| | Anseriformes | 2.592 | 2.000 | 4.000 | 2.318 | 2.058 | 3.006 | 0.00999 |
| | nonhuman-mammal | 4.922 | 4.000 | 6.000 | 2.306 | 2.050 | 3.012 | 0.00999 |
| | raptors | 4.106 | 3.000 | 6.000 | 2.321 | 2.056 | 3.018 | 0.00999 |
| | shorebirds | 8.008 | 8.000 | 8.000 | 2.324 | 2.054 | 3.030 | 0.00999 |
| | Strigiformes | 3.664 | 3.000 | 5.000 | 2.316 | 2.048 | 3.020 | 0.00999 |
| | Passeriformes | 10.000 | 10.000 | 10.000 | 1.847 | 1.414 | 2.156 | 0.00999 |
| domestic wild turkey | AI | 20.709 | 18.436 | 23.018 | 57.149 | 54.852 | 59.336 | 0.00199 |
| | PS | 144.599 | 135.000 | 155.000 | 318.740 | 313.372 | 323.543 | 0.00199 |
| | turkey | 9.784 | 7.000 | 15.000 | 2.631 | 2.294 | 3.097 | 0.00199 |
| | domestic | 11.565 | 11.000 | 14.000 | 2.613 | 2.283 | 3.123 | 0.00199 |
| | wild | 27.688 | 19.000 | 37.000 | 9.602 | 8.294 | 12.082 | 0.00199 |
| domestic wild | AI | 15.148 | 13.665 | 16.980 | 43.950 | 41.232 | 46.378 | 0.00199 |
| | PS | 108.882 | 104.000 | 115.000 | 241.071 | 234.363 | 246.802 | 0.00199 |
| | Wild | 38.178 | 37.000 | 44.000 | 12.628 | 10.511 | 16.301 | 0.00199 |
| | domestic | 23.096 | 19.000 | 28.000 | 3.390 | 2.924 | 4.156 | 0.00199 |
| domestic wild backyard bird | AI | 10.854 | 9.014 | 12.775 | 30.252 | 28.751 | 31.921 | 0.00999 |
| | PS | 84.263 | 78.000 | 93.000 | 164.462 | 161.920 | 166.440 | 0.00999 |
| | backyard bird | 7.297 | 6.000 | 9.000 | 2.007 | 1.693 | 2.267 | 0.00999 |
| | domestic | 11.243 | 10.000 | 13.000 | 2.012 | 1.693 | 2.320 | 0.00999 |
| | wild | 64.840 | 63.000 | 68.000 | 16.674 | 14.143 | 20.300 | 0.00999 |

**Table S18:** Results of BaTs analysis for each discrete trait set used. Association Index (AI), Parsimony score (PS) and the maximum monophyletic clade size for each trait in the analysis are listed with their mean and 95% CI.

| Analysis | Trait | Original | tip shuffle | Number of taxa |
|---|---|---|---|---|
| Migration | Domestic | 0.1194 | 0.1891 | 100 |
| | Nonhuman Mammal | 0.0141 | 0.2071 | 100 |
| | Migratory | 0.4569 | 0.2171 | 100 |
| | Part Migratory | 0.4065 | 0.1917 | 100 |
| | Sedentary | 0.0031 | 0.1948 | 100 |
| Global | Asia | 0.2117 | 0.001 | 294 |
| | North America | 0.6759 | 0.998 | 1333 |
| | Europe | 0.1124 | 0.001 | 300 |
| Flyway | Atlantic | 0.9503 | 0.301 | 250 |
| | Central | 0.0232 | 0.253 | 250 |
| | Mississippi | 0.0252 | 0.247 | 250 |
| | Pacific | 0.0011 | 0.197 | 250 |
| Domestic, Wild, Turkey (first titration 1:1:1) | Domestic | 0.0263 | 0.063 | 173 |
| | Turkey | 0.0113 | 0.063 | 173 |
| | Wild | 0.9623 | 0.872 | 346 |
| Domestic, Wild (first titration, 1:1) | Domestic | 0.114 | 0.517 | 270 |
| | Wild | 0.895 | 0.483 | 270 |
| Domestic, Wild, Backyard bird (first titration, 1:1 Domestic:backyard bird 25% all wild birds) | Wild | 0.9992 | 0.919 | 193 |
| | Domestic | 0.004 | 0.0037 | 85 |
| | Backyard bird | 0.0003 | 0.0043 | 85 |
| Host orders – Equal 1 | Galliformes | 0.1189 | 0.1309 | 100 |
| | Strigiformes | 0.029 | 0.1568 | 99 |
| | Raptors | 0.1019 | 0.1838 | 100 |
| | Nonhuman mammal | 0.1409 | 0.1568 | 100 |
| | Shorebird | 0.1499 | 0.1548 | 100 |
| | Passeriformes | 0.013 | 0.02 | 57 |
| | Anseriformes | 0.4466 | 0.1968 | 100 |
| Host orders – Equal 2 | Galliformes | 0.0789 | 0.158 | 100 |
| | Strigiformes | 0.026 | 0.154 | 99 |
| | Raptors | 0.1069 | 0.158 | 100 |

| | | | | |
|---|---|---|---|---|
| | Nonhuman mammal | 0.1099 | 0.178 | 100 |
| | Shorebird | 0.1269 | 0.169 | 100 |
| | Passeriformes | 0.018 | 0.024 | 57 |
| | Anseriformes | 0.5335 | 0.154 | 100 |
| Host orders – Equal 3 | Galliformes | 0.092 | 0.162 | 100 |
| | Strigiformes | 0.0421 | 0.157 | 99 |
| | Raptors | 0.1494 | 0.155 | 100 |
| | Nonhuman mammal | 0.1696 | 0.179 | 100 |
| | Shorebird | 0.0613 | 0.156 | 100 |
| | Passeriformes | 0.0192 | 0.024 | 57 |
| | Anseriformes | 0.4674 | 0.164 | 100 |
| Host orders – Proportional 1 | Galliformes | 0.003 | 0.046 | 65 |
| | Strigiformes | 0.001 | 0.029 | 44 |
| | Raptors | 0.0759 | 0.469 | 167 |
| | Nonhuman mammal | 0.001 | 0.033 | 33 |
| | Shorebird | 0.0529 | 0.009 | 83 |
| | Passeriformes | 0.001 | 0.012 | 31 |
| | Anseriformes | 0.9691 | 0.399 | 232 |
| Host orders - Proportional 2 | Galliformes | 0.008 | 0.087 | 65 |
| | Strigiformes | 0.001 | 0.055 | 44 |
| | Raptors | 0.0709 | 0.384 | 167 |
| | Nonhuman mammal | 0.001 | 0.094 | 33 |
| | Shorebird | 0.0519 | 0.021 | 83 |
| | Passeriformes | 0.001 | 0.034 | 31 |
| | Anseriformes | 0.9681 | 0.321 | 232 |
| Host orders - Proportional 3 | Galliformes | 0.001 | 0.080 | 65 |
| | Strigiformes | 0.002 | 0.043 | 44 |
| | Raptors | 0.3387 | 0.368 | 167 |
| | Nonhuman mammal | 0.0709 | 0.128 | 33 |
| | Shorebird | 0.001 | 0.034 | 83 |
| | Passeriformes | 0.006 | 0.025 | 31 |

| | | | | |
|---|---|---|---|---|
| | Anseriformes | 0.5904 | 0.318 | 232 |

**Table S19:** Root state probabilities for discrete traits of each analysis in the study using the discrete trait shuffling test.

**Table S20.** Acknowledgments table for GISAID isolates used in these analyses. See attached table file.
